# Towards Synthetic PETtrophy: Engineering *Pseudomonas putida* for concurrent polyethylene terephthalate (PET) monomer metabolism and PET hydrolase expression

**DOI:** 10.1101/2022.04.21.489007

**Authors:** Oliver F. Brandenberg, Olga T. Schubert, Leonid Kruglyak

**Affiliations:** Department of Human Genetics, Department of Biological Chemistry, and Howard Hughes Medical Institute, University of California Los Angeles, Los Angeles, United States; Department of Environmental Microbiology, Swiss Federal Institute of Aquatic Science and Technology, Dübendorf, Switzerland

**Keywords:** polyethylene terephthalate (PET), PET hydrolase, biodegradation, metabolic engineering, adaptive laboratory evolution, membrane display, enzyme secretion, whole-cell biocatalysis

## Abstract

**Background:** Biocatalysis offers a promising path for plastic waste management and valorization, especially for hydrolysable plastics such as polyethylene terephthalate (PET). Microbial whole-cell biocatalysts for simultaneous PET degradation and growth on PET monomers would offer a one-step solution toward PET recycling or upcycling. We set out to engineer the industry-proven bacterium *Pseudomonas putida* for (i) metabolism of PET monomers as sole carbon sources, and (ii) efficient concurrent extracellular expression of PET hydrolases. We pursued this approach for both PET and the related polyester polybutylene adipate co-terephthalate (PBAT), aiming to learn about the determinants and potential applications of bacterial polyester-degrading biocatalysts.

**Results:** *P. putida* was engineered to metabolize the PET and PBAT monomer terephthalic acid (TA) through genomic integration of four tphII operon genes from *Comamonas sp*. E6. Efficient cellular TA uptake was enabled by a point mutation in the native *P. putida* membrane transporter mhpT. Metabolism of the PET and PBAT monomers ethylene glycol and 1,4-butanediol was achieved through adaptive laboratory evolution. We then used fast design-build-test-learn cycles to engineer extracellular PET hydrolase expression, including tests of (i) the three PET hydrolases LCC, HiC, and IsPETase; (ii) genomic versus plasmid-based expression, using expression plasmids with high, medium, and low cellular copy number; (iii) three different promoter systems; (iv) three membrane anchor proteins for PET hydrolase cell surface display; and (v) a 30-mer signal peptide library for PET hydrolase secretion. PET hydrolase surface display and secretion was successfully engineered but often resulted in host cell fitness costs, which could be mitigated by promoter choice and altering construct copy number. Plastic biodegradation assays with the best PET hydrolase expression constructs genomically integrated into our monomer-metabolizing *P. putida* strains resulted in various degrees of plastic depolymerization, although self-sustaining bacterial growth remained elusive.

**Conclusion:** Our results show that balancing extracellular PET hydrolase expression with cellular fitness under nutrient-limiting conditions is a challenge. The precise knowledge of such bottlenecks, together with the vast array of PET hydrolase expression tools generated and tested here, may serve as a baseline for future efforts to engineer *P. putida* or other bacterial hosts towards becoming efficient whole-cell polyester-degrading biocatalysts.

## Introduction

Polyethylene terephthalate (PET) is a linear aromatic polyester composed of repeating units of terephthalic acid (TA) and ethylene glycol (EG) and is one of the most common plastics in the world. Annual production of virgin PET amounts to > 60 million metric tons, with roughly equal amounts processed as fiber (‘polyester’) for the textile industry and as rigid containers for food and beverage packaging.^1–3^ Due to the short lifetimes of many PET products, accumulation of PET waste in landfills or terrestrial and aquatic ecosystems is a major concern.^4,5^ Therefore, better incentives for PET recycling or safe options for PET disposal are urgently needed. In addition, improved options for PET recycling or upcycling may translate into less use of crude oil and contribute to the establishment of a circular plastic (bio)economy.^6–9^

The small fraction of PET waste collected for recycling is mostly processed mechanically; however, PET can be recycled mechanically only a few times due to processing-induced chain scission and ensuing degradation of mechanical properties. ^10^ To circumvent this, chemical recycling (including glycolysis, hydrolysis, methanolysis, and pyrolysis, among others) can be used to depolymerize PET to its constituents TA and EG, followed by synthesis of new virgin PET.^11^ While this offers a truly circular recycling path, chemical depolymerization requires high energy input and harsh reaction conditions. Therefore, the use of enzymes for biocatalytic PET depolymerization under mild, aqueous, and environmentally friendly reaction conditions has gained significant interest.

In the last two decades, a growing number of enzymes from bacterial and fungal sources with PET hydrolase activity have been characterized. These enzymes typically encompass serine hydrolases which natively function as cutinases, lipases, or carboxylesterases, and show promiscuous PET hydrolyzing activity.^1,12–16^ Enzymatic PET hydrolysis by native, non-engineered enzymes typically occurs on the order of days to weeks and is severely impaired by the semi-crystalline structure of commercial PET, which consists of amorphous regions interspersed with crystalline regions that are highly refractory to enzymatic attack.^1,17,18^ PET hydrolases are frequently engineered to function near the glass transition temperature (T_g_) of PET at ≥ 70°C where amorphous regions become more accessible, with the latest generation of enzymes capable of degrading PET samples at elevated temperatures within as little as 10 hours.^19–21^

In addition to enzymatic PET depolymerization, biology can be leveraged for recycling applications downstream of polymer hydrolysis. The PET monomers TA and EG can be used as metabolic feedstocks for biochemical production pathways in engineered microbes. Recent examples include the engineering of bacterial strains capable of transforming TA and/or EG into value-added products such as β-ketoadipic acid, glycolic acid, and the bioplastics polyhydroxyalkanoate (PHA) and poly(amide urethane).^20,22–25^

Considering these two facets of biology for PET recycling, a clear continuation would be the combination of enzymatic PET depolymerization and monomer metabolism in the same microbial organism (Figure 1). Indeed, nature itself already delivered this solution: In 2016, researchers isolated the bacterium *Ideonella sakaiensis* 201-F6 from a PET recycling site and showed that it possesses the enzymatic machinery to both depolymerize PET and utilize the resulting monomers as carbon source for growth.^26^ Here, we set out to replicate this feature by engineering a bacterium with the capacity to both degrade PET and metabolize the monomers as carbon sources, a capacity we refer to as *synthetic PETtrophy*. In addition to PET, we pursued this approach for the more readily biodegradable polyester polybuylene adipate co-terephthalate (PBAT). We implemented this strategy using the soil bacterium *Pseudomonas putida,* which, in contrast to *I. sakaiensis*, is already established as a promising host for industrial biocatalytic processes due to robust growth under harsh conditions and has a vast array of tools and knowledge available for genetic engineering.^27–29^

**Figure 1:**
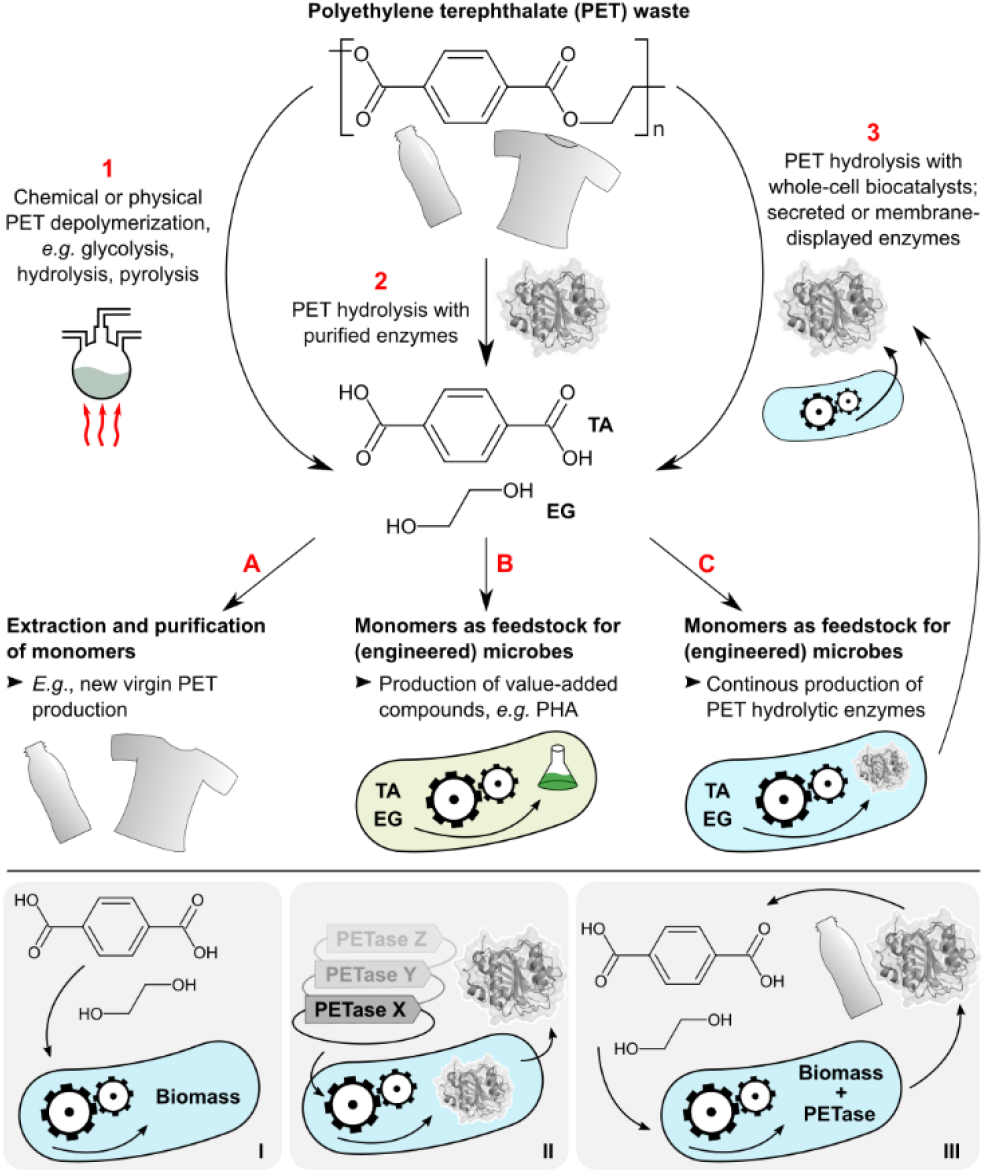
Synthetic PETtrophy in relation to other biocatalytic strategies for PET waste valorization. Top panel: PET waste can be depolymerized to TA and EG using chemical means (1), with purified PET hydrolases (2), or using whole-cell biocatalysts displaying or secreting PET hydrolases (3). TA and EG can be extracted and processed into new virgin PET (A); used as feedstocks for engineered microbes and production of value-added bioproducts (B); or used as feedstocks for engineered microbes producing PET hydrolases in a CBP-like one-pot system (C). Path 2-A was explored by Tournier *et al*.^19^ and Lu *et al*.^20^; path 1-B was explored by Kenny *et al*.^22,33^, Werner *et al*.^34^ and Kim *et al*.^25^; path 2-B was described by Tiso *et al*.^24^ and Lu *et al*.^20^; path 3-C is what *I. sakaiensis* perfoms natively^26^ and was explored in this study. Additionally, several recent studies described whole-cell biocatalysts displaying or secreting PET hydrolases (3) without attempting downstream extraction or microbial utilization of released monomers. Bottom panel: sequence of engineering steps attempted in this study. First, we installed TA, EG or BD metabolism in *P. putida* using genetic engineering and adaptive laboratory evolution (I); secondly, we tested a wide variety of expression systems for PET hydrolases (II); third, we attempted to combine these efforts to obtain a self-sustaining whole-cell PET or PBAT depolymerization system (III). Enzyme structure PDB 5xjh.^35^

Successful engineering of synthetic PETtrophy would be a step towards consolidated bioprocessing (CBP) of PET waste. The concept of consolidated bioprocessing originated in the field of lignocellulosic biomass utilization and includes synthesis of hydrolytic enzymes, polymer hydrolysis, and downstream routing of monomers into biosynthetic pathways by the same microbial organism in a single fermentation setup.^6,30,31^ A whole-cell biocatalyst for PET continuous bioprocessing would simplify the recycling workflow and may be able to deal with contaminated or mixed-plastic waste streams. It may also lend itself to decentralized *in situ* bioremediation of PET waste (*e.g*., in landfills or dedicated composting facilities), thus rendering PET a biodegradable waste and ‘ecocyclable’.^3,32^ Additionally, we were motivated to discover potential bottlenecks and dos-and-don’ts encountered in the synthetic PETtrophy engineering endeavor in order to understand the fundamental determinants of bacterial PET utilization as a carbon source.

We first engineered *P. putida* to accept TA, EG and 1,4-butanediol (BD, a component of PBAT) as sole carbon and energy sources. Next, we used rapid design-build-test-learn cycles to achieve functional expression of diverse PET hydrolases. Lastly, we tested the most promising strains in whole-cell PET or PBAT degradation experiments. While we did not obtain fully self-sustaining plastic-depolymerizing bacterial systems, our work provides detailed insights into engineering of PET monomer metabolism, PET hydrolase expression construct design, and the challenges of balancing enzyme expression and cell viability under nutrient-limiting conditions, all of which may prove useful for future efforts towards establishing synthetic PETtrophy.

## Results

### Establishing TA catabolism in Pseudomonas putida KT2440

Our first goal was to enable *Pseudomonas putida* KT2440 to use TA as sole carbon and energy source. A common aromatic acid catabolite in *P. putida* is protocatechuate (PCA), which feeds into central carbon metabolism via the β-ketoadipate pathway.^36^ However, *P. putida* KT2440 cannot convert TA to PCA; importantly though, enzyme cascades catalyzing this transformation have been found in various bacteria.^30^ We selected four genes from the *tph_II_* operon of *Comamonas sp*. E6,^37^ encoding the multi-subunit TA dioxygenase TphA_II_ that transforms TA to 1,2-dihydroxy-3,5-cyclohexadiene-1,4-dicarboxylate (DCD), and the DCD dehydrogenase TphB_II_ that transforms DCD to PCA, to endow *P. putida* with the ability to transform TA to PCA (Figure 2A). We designed a synthetic operon consisting of the *Comamonas sp*. E6 genes *tphA1_II_, tphA2_II_, tphA3_II_* and *tphB_II_* under control of the constitutive P_Em7_ promoter and endowed it with a bicistronic translational coupler (BCD2) to ensure efficient transcription and translation in *P. putida* (Supplementary Figure 1).^38,39^ We did not include a selectable antibiotic resistance marker in the knock-in cassette, as the capacity to catabolize TA is itself a selectable trait. The operon was cloned into the mini-TN5 transposon vector pBAMD1-2 (yielding vector pBAMD-tphII), enabling random integration of the operon into the *P. putida* genome.^40^

**Figure 2:**
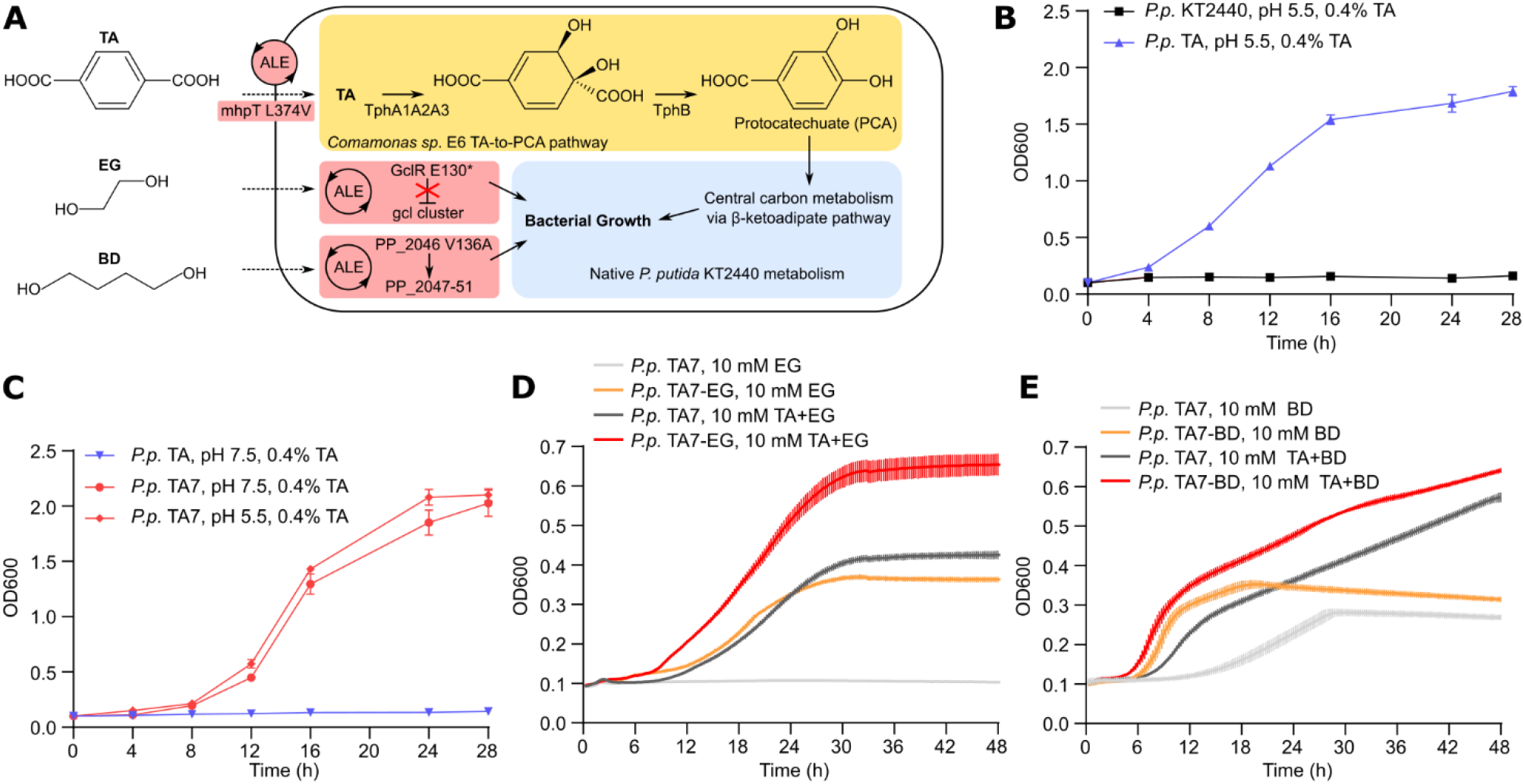
Engineering TA, EG and BD metabolism in *P. putida*. (A) Overview of the engineering steps, including genomic insertion (yellow) and ALE (red), to endow *P. putida* with the capacity to use TA, EG and BD as sole carbon sources. (B) Growth of *P. putida* KT2440 and *P. putida* TA in shake flasks with M9 minimal medium, 0.4% TA, pH 5.5. (C) Growth of *P. putida* TA and *P. putida* TA7 in shake flasks with M9 minimal medium, 0.4% TA, pH 5.5 or pH 7.5. (D) Growth of *P. putida* TA7 and *P. putida* TA7-EG analyzed in a plate reader assay with M9 minimal medium, 10 mM EG or 10 mM EG + 10 mM TA, pH 7.5. (E) Growth of *P. putida* TA7 and *P. putida* TA7-BD analyzed in a plate reader assay with M9 minimal medium, 10 mM BD or 10 mM BD + 10 mM TA, pH 7.5. Data shown are mean and SD, n=2 (B and C) or n=6 (D and E).

pBAMD-tphII was delivered to *P. putida* KT2440 by electroporation, and cells were cultured in M9 minimal medium supplemented with 0.4% TA as sole carbon source to select for cells with functional integration of the synthetic *tph*_II_ operon. We adjusted the culture medium to pH 5.5 to ensure that a certain fraction of TA molecules is fully protonated and thus able to diffuse through the cell membrane in the absence of dedicated TA transporters.^37,41^ We observed bacterial growth two days post-transformation and passaged the culture over ten days, followed by subsequent analysis of single clones. The best-performing clone was designated as strain *P. putida* TA; whole-genome sequencing revealed integration of the *tphII* cassette into the PP_4627 (cidR) locus (Supplementary Table 1). This strain showed robust growth in M9 minimal medium supplemented with TA, while no growth was observed for *P. putida* KT2440 (Figure 2B).

We next set out to modify *P. putida* TA to grow at neutral pH. At pH 7, TA is in its partially or fully deprotonated form, necessitating TA transporters for cellular uptake. Considering that *P. putida* has a variety of membrane transporters for aromatic compounds,^42^ we reasoned that adaptive laboratory evolution (ALE) may succeed in repurposing an existing *P. putida* transporter to function as a TA uptake transporter. Thus, we cultured *P. putida* TA in M9 medium with 0.4% TA adjusted to pH 6.5 for six consecutive passages over seven days, after which we observed improved bacterial growth (Supplementary Figure 2). Isolation and analysis of single clones yielded strain *P. putida* TA7, which showed strongly improved growth over parent strain *P. putida* TA at neutral pH (Figure 2C). Whole-genome sequencing (Supplementary Table 2) of both strains revealed a missense mutation in *P. putida* TA7 in a MarR-family putative transcriptional regulator for hydroxycinnamic acid degradation (PP_3359; c.151C>T, R51C)^43^ and a missense mutation in the putative 3-(3-hydroxyphenyl)proprionate transporter mhpT (PP_3349; c.1120C>G, L374V)^44^. This indicated that mhpT has promiscuous TA transport activity, which was likely enhanced by the L374V mutation. The mutation in the putative transcription factor PP_3359, located only 12 kbp upstream of mhpT, may influence mhpT expression. We confirmed the function of mhpT^L374V^ as a putative TA transporter by rescuing growth of *P. putida* TA in M9-TA medium at pH 7.5 by plasmid-based overexpression of mhpT^L374V^ (Supplementary Figure 3).

Overall, through genomic knock-in of four genes from the TA-to-PCA pathway of *Comamonas sp*. E6 followed by brief ALE, we obtained strain *P. putida* TA7 capable of robust growth on TA as sole carbon and energy source.

### Establishing EG and BD catabolism in P. putida TA7

We next aimed to establish EG metabolism in *P. putida* TA7 based on two considerations. First, high EG concentrations negatively affected growth of *P. putida* TA7 (Supplementary Figure 4A). Second, turning EG from a liability to a potential carbon source would support robust growth on PET, especially considering that PET hydrolysis is the rate-limiting step, and any released monomers should be utilized as efficiently as possible.

*P. putida* possesses the enzymatic machinery required to utilize EG as carbon and energy source; however, transcription of the genes encoding the two key enzymes Gcl and GlxR is constitutively repressed by the transcriptional repressor GclR (Supplementary Figure 4B).^45–47^. Thus, we attempted ALE of *P. putida* TA7 for growth on EG, using cultures in M9 minimal medium supplemented with 25 mM EG and 5 mM TA.

We observed improved bacterial growth after approximately one week and isolated single clones after eleven days of passaging (Supplementary Figure 5); the best-performing strain was designated *P. putida* TA7-EG. This strain showed both a shorter lag phase and higher final OD600 compared to *P. putida* TA7 when grown in M9 supplemented with equimolar amounts of TA and EG (Figure 2D). Cell dry weight measurements of *P. putida* TA7-EG cultures confirmed an improved capacity of *P. putida* TA7-EG to accumulate biomass (Supplementary Figure 6). Both strains showed comparable growth in LB medium, while *P. putida* TA7-EG showed a slight growth advantage with TA as only carbon source (Supplementary Figure 7 A, B). Whole-genome sequencing of *P. putida* TA7-EG revealed a nonsense mutation in GclR (PP_4283; c.388G>T, E130* Supplementary Table 2), likely resulting in GclR loss of function, which is consistent with previous studies on the molecular basis of improved EG catabolism.^45–47^

For the PBAT monomer BD, we set up two ALE cultures with *P. putida* TA7 in M9 minimal medium supplemented with either 25 mM BD and 5 mM TA, or 25 mM BD, 5 mM TA, and 25 mM adipic acid (AA), the second aliphatic linker in PBAT. Both ALE cultures showed similar growth profiles, and we observed improved growth after ten days, when we isolated and tested single clones (Supplementary Figure 8). The strain showing best growth characteristics was designated *P. putida* TA7-BD (Figure 2E). Both *P. putida* TA7 and *P. putida* TA7-BD showed comparable growth in LB medium, while we observed a minor detrimental effect when AA was added to the medium (Supplementary Figure 7 C, D). *P. putida* TA7-BD carried a missense mutation in PP_2046 (c.407T>C, V136A), a gene previously implicated as a transcriptional regulator of an operon involved in BD metabolism, with gain of function mutations resulting in improved growth on BD.^48^

Interestingly, across all ALE experiments and engineered strains, we did not detect any mutations in the *tph_II_* cassette (Supplementary Table 2). This indicates that the chosen operon design, including the transcriptional output from the P_EM7_ promoter, is likely close to optimal; at least, no mutations appeared to have a positive effect on bacterial growth strong enough to fix in the population during our ALE experiments.

In summary, ALE rapidly succeeded in endowing *P. putida* TA7 with the ability to use either EG or BD as additional carbon and energy sources, and the resulting *P. putida* TA7-EG and *P. putida* TA7-BD represent robust strains for engineering PET hydrolase expression.

### Engineering PET hydrolase surface display

We next aimed to endow *P. putida* with the enzymatic machinery required to depolymerize PET or PBAT. We first attempted display of PET hydrolases on the bacterial outer membrane, speculating that such surface display may provide higher local enzyme concentrations and better adsorption of the PET hydrolase to the plastic surface, thereby promoting more efficient hydrolysis.

To obtain a stable and functional PET hydrolase display system for *P. putida*, we cloned and tested three different membrane display anchors, all previously shown to function in *P. putida* (Supplementary Figure 9). First, we used a construct based on the *E. coli* autotransporter EhaA.^49^ This construct consists of the *P. putida* OprF signal peptide (37 residues), the cargo domain (*i.e*., the PET hydrolase to be displayed), a G_4_SGGS(G_4_S)_3_ linker,^50^ and the C-terminal EhaA autotransporter domain (488 residues). Second, we employed a construct based on the commonly used ice nucleation protein display anchors, comprising the N-terminal 203 residues fused to the C-terminal 97 residues of the ice nucleation protein inaV from *Pseudomonas syringae*.^51^ This anchor domain is followed by the G_4_SGGS(G_4_S)_3_ linker and the PET hydrolase cargo domain. Last, we employed the *P. putida* OprF outer membrane protein as anchor domain, as previously demonstrated using OprF from *Pseudomonas aeruginosa.^52^* This construct comprised the N-terminal 206 residues of *P. putida* OprF, followed by the G_4_SGGS(G_4_S)_3_ linker and the PET hydrolase cargo domain. Of note, full-length OprF has been shown to adopt two different conformations, with the cargo fusion site V206 predicted to be either periplasmic or extracellular.^53^ Based on reports of extracellular exposure of V206 in OprF homologs,^54,55^ we decided to try this construct for PET hydrolase display.

All display constructs were put under control of the rhamnose-inducible P_rhaB_ promoter, shown to provide low levels of leaky transcription and high induction levels in *P. putida*.^56,57^ All constructs also featured the BCD2 translational coupler used for our *tph_II_* operon described above,^39^ a hexa-His tag attached to the cargo domain opposite from the anchor domain, unique PstI and NcoI restriction sites for facile cloning of different cargo domains into the display systems, and unique AvrII and HindIII sites to transfer the entire display cassettes between vectors (Supplementary Figure 9). Furthermore, as expression of heterologous membrane proteins often leads to bacterial growth defects, we implemented an extra layer of control over display construct expression levels by modulating gene dosage using plasmid vectors with low (RK2), medium (pBBR1), and high (pRO1600/ColE1) cellular copy number.^58^ All plasmids carried a Km^R^ selectable marker and were constructed from plasmids obtained through the Standard European Vector Architecture (SEVA) repository.^59^ In total, this yielded nine display construct-vector combinations (Figure 3A).

**Figure 3:**
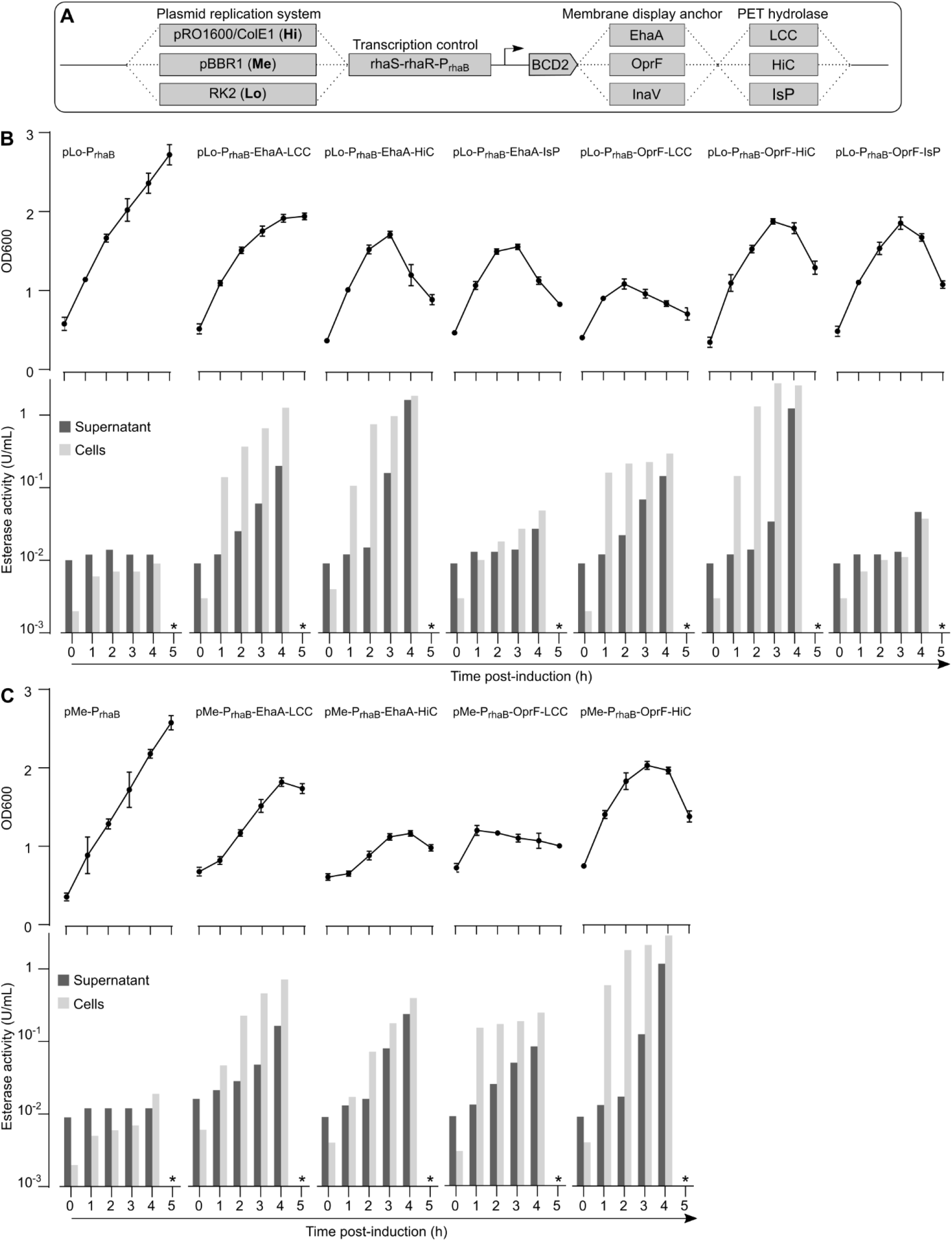
Testing PET hydrolase membrane display systems. (A) Overview of combinatorial test construct design for PET hydrolase membrane expression, including plasmids providing low, medium, and high cellular copy number, transcriptional control by the P_rhaB_ system, translation initiation via the BCD2 translational coupler, three different membrane display anchors, and three different PET hydrolases. (B) OD600 and esterase activity of *P. putida* TA7-EG shake flask cultures expressing the indicated display constructs from the RK2 low copy number plasmid, measured for 5 h post-induction (induction at t=0 h). (C) OD600 and esterase activity of *P. putida* TA7-EG shake flask cultures expressing the indicated display constructs from the pBBR1 medium copy number plasmid, measured for 5 h post-induction (induction at t=0 h). The * in (B) and (C) indicates that esterase activity was not measured at the t=5 h timepoint due to obvious cell lysis. Data shown are derived from one representative experiment with duplicate cultures.

We chose three well-known PET hydrolases for testing: the *Humicola insolens* cutinase (HiC), leaf compost cutinase (LCC), and *Ideonella sakaiensis* PETase (IsP). HiC is a thermostable fungal cutinase of 194 residues capable of almost completely degrading low-crystallinity PET film within 96 h at 70°C, with TA as predominant hydrolysis product.^60^ LCC is a 293 residue protein and was isolated through functional screening of a compost metagenomic library.^61^ LCC and variants thereof have long been recognized as some of the most promising PET hydrolases, achieving hydrolysis of amorphous PET to TA and EG on the gram to kg scale.^19,24^ IsP is a 290 residue protein that is regarded as the first identified enzyme with the native function to enable bacterial growth on PET as sole carbon and energy source. IsP stands out for its high PET hydrolysis activity at mesophilic temperatures (30-40°C), where it surpasses thermostable PET hydrolases; however, LCC and HiC are more active when used at their optimum temperatures.^26,60,61^ In contrast to HiC and LCC, IsP yields MHET as main hydrolysis product, which is further hydrolyzed to TA in *I. sakaiensis* by MHETase.^26^ However, we omitted MHETase from our approach, as recent studies indicated TA accumulation in long-term PET hydrolysis setups with IsP alone,^62^ and we wanted to obtain a clean comparison of PET hydrolase expression levels without expression of auxiliary enzymes.

We first cloned ORFs of both HiC and LCC (omitting their signal peptides) into the set of nine display vectors (IsP was included only later). Upon transformation of *P. putida* TA7-EG with these plasmids, we observed only sporadic colony formation for all pRO1600/ColE1 plasmids; presumably, leaky expression from these high copy number plasmids was sufficient to induce bacterial growth defects, and these plasmids were excluded from further analyses. Additionally, initial analyses showed no PET hydrolase expression for any of the InaV display constructs, and they were also excluded from further analyses. For the remaining plasmids, we tested cell viability and PET hydrolase expression over time in shaking cultures (Figure 3B). As a proxy for cell viability we measured OD600, while we used the commonly employed esterase model substrate *para*-nitrophenylbutyrate (pNPB) and a spectrophotometric read-out of 4-nitrophenol formation to assess enzymatic activity of expressed PET hydrolases (Supplementary Figure 10). We performed the pNPB esterase assay for both cleared culture supernatant and the cell fraction.

For *P. putida* TA7-EG transformed with empty vectors (lacking PET hydrolases), we observed stable growth and very low esterase activity both in the cell fraction and the culture supernatant following induction (Figure 3B, C). In contrast, for all strains expressing PET hydrolase display constructs, we observed strong increases in cell-associated esterase activity, indicating functional PET hydrolase surface display. The highest cell-associated esterase activity was measured for OprF-HiC at 2.68 U/mL 3 h post-induction, 330-fold above the empty-vector control. SDS-PAGE analysis of the membrane fraction of PET hydrolase-expressing cells followed by LC-MS analysis confirmed correct expression and localization of the EhaA membrane display constructs, while the observed bands for OprF constructs were less clear (Supplementary Figure 11). Approximately 1-2 h after the cell-associated esterase signal we also observed spikes in esterase activity in the culture supernatant, likely originating from cell lysis. Indeed, for all bacteria expressing PET hydrolase display constructs, we observed growth defects following induction, with most cultures collapsing (decrease in OD600 with visible cell lysis, clumping, and increase in medium viscosity) starting 3-4 h post-induction (Figure 3B, C). Plate reader analysis of OD600 for 24 h post-induction confirmed the collapse of PET hydrolase-expressing cultures (Supplementary Figure 12). Growth defects appeared to be slightly more pronounced for the medium copy number pBBR1 plasmids compared to the low copy number RK2 plasmids.

To investigate the observed cell death upon PET hydrolase display construct expression, we first tested whether cytosolic expression of the PET hydrolase ORFs (without the membrane display anchors) would cause similar growth defects. Surprisingly, we observed that cytosolic expression of LCC indeed resulted in a similar collapse of the bacterial culture. In contrast, cytosolic expression of HiC or IsP resulted in normal bacterial growth rates (Supplementary Figure 13). The reason for the observed LCC toxicity in *P. putida* remains to be determined, while surface display construct expression appears to be the major culprit for all three enzymes.

Next, we tested whether titration of the inducer, rhamnose, could mitigate the problem of expression construct toxicity. We used one of the constructs showing the highest PET hydrolase membrane expression and the lowest apparent toxicity, EhaA-LCC in the RK2 plasmid background, and varied rhamnose inducer concentration from 0.5 to 8 mM. This revealed a very limited dynamic range for expression from the P_rhaB_ promoter, with 0.5 mM rhamnose already providing high levels of induction and leading to culture collapse, as seen for all other inducer concentrations (Supplementary Figure 14).

Thus, the EhaA and OprF display anchors resulted in successful cell surface display of PET hydrolases. However, membrane display construct expression also caused cellular instability, which could not be mitigated by lower inducer concentrations.

### Engineering PET hydrolase secretion

Parallel to engineering PET hydrolase cell surface display we explored options for PET hydrolase secretion, which we speculated may result in lower cellular toxicity compared to membrane display.

Protein secretion is highly dependent on the N-terminal signal peptide (SP). Secretion levels can differ by orders of magnitude depending on the SP, and a highly efficient SP for one protein may not work well for another.^63^ In addition, extracellular protein secretion in Gram-negative bacteria such as *P. putida* is incompletely understood.^64,65^ Therefore, we used a SP library screening approach to achieve efficient PET hydrolase secretion. We tested a set of 30 SPs, 28 originating from *Pseudomonas* proteins annotated as secreted or periplasmic, and the remaining two being the SPs from the *I. sakaiensis* PETase and MHETase (Supplementary Table 3). Of these 30 SPs, 19 were predicted to be Sec-dependent, while 11 were predicted to be Tat-dependent. Instead of cloning each SP individually, we cloned the library in bulk using a specifically designed plasmid such that the LCC, HiC and IsP ORFs were provided with the SP library in-frame at the 5’ end (Figure 4A, B).

**Figure 4:**
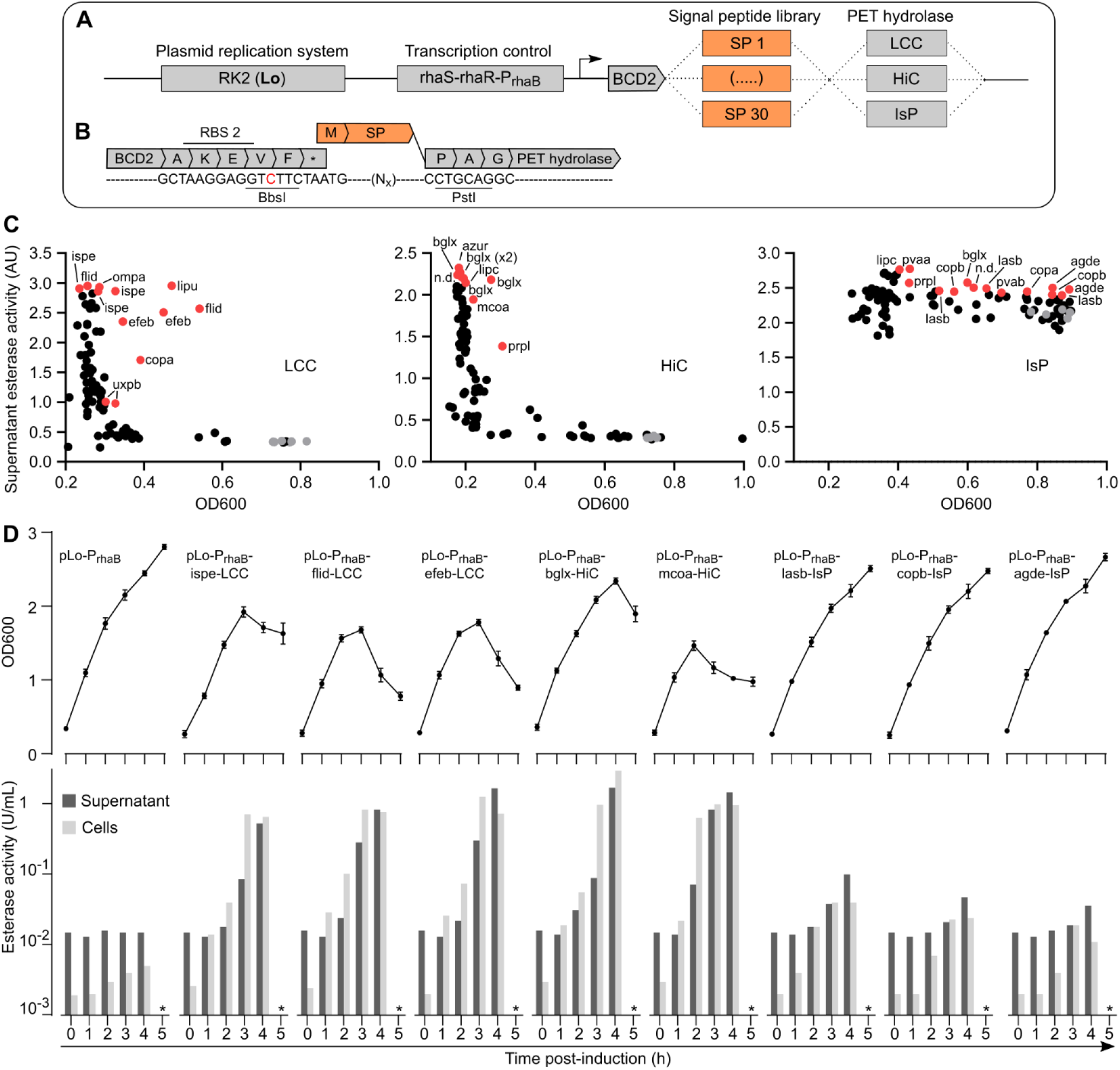
Engineering PET hydrolase secretion. (A) Overview of plasmid design to screen a 30-mer SP library for LCC, HiC, and IsP secretory expression in *P. putida* TA7-EG. We used the RK2 low copy number plasmid, the P_rhaB_ induction system, BCD2 translational coupler, and a SP library cloned at the 5’ end of the LCC, HiC, and IsP ORFs. (B) Sequence view of the 3’ end of the BCD2 translational coupler, where we introduced a silent point mutation (red) to create a unique BbsI restriction site. Together with a unique PstI site preceding the PET hydrolase ORFs, this allowed facile in-frame cloning of the SP library. (C) SP library screening results, showing culture OD600 (measured 20 h post-induction) plotted against culture supernatant esterase activity (measured 3 h post-induction). Each dot represents a single clone (one well of a 96-well culture plate); grey dots are e.v. controls, red dots are sequenced clones, with the identified SPs indicated. (D) Shake flask validation of *P. putida* TA7-EG expressing the best identified SP-PET hydrolase constructs. Data depict OD600 (top) and supernatant or cell-associated esterase activity measured for 5 h following induction (bottom). The * in (D) indicates that esterase activity was not measured at the t=5 h timepoint due to obvious cell lysis for most samples. Data shown are derived from one representative experiment with duplicate cultures.

We retained the P_rhaB_ promoter, BCD2 bicistronic translational coupler, and RK2 low copy number oriV already used for the membrane display constructs.

Using a library-based approach necessitated a screening procedure to identify the most effective SPs for each PET hydrolase. Thus, *P. putida* TA7-EG was transformed with each of the three SP-PET hydrolase libraries, and for each library 90 single clones (3-fold oversampling) were analyzed in 96-well liquid shaking cultures. To assess PET hydrolase secretion, we measured both esterase activity in the supernatant (3 h post-induction) and OD600 as proxy for cell viability (3 h and 20 h post-induction) (Figure 4C); cells transformed with empty vector served as controls. For both LCC and HiC, we observed that almost all SP library clones showed strongly reduced OD600 at 20 h post-induction; the few clones showing higher, control-like OD600 showed no PET hydrolase expression (*i.e.*, no esterase activity in the culture supernatant). Furthermore, for both the LCC and HiC libraries, we observed a wide range of esterase activities in the culture supernatants 3 h post-induction, indicative of different PET hydrolase secretion efficiencies of the different SPs. For IsP, the low intrinsic activity of this enzyme in the pNPB assay resulted in a lower resolution between the library clones and controls, although it remained possible to identify clones showing higher supernatant esterase activity. At the same time, more clones retained high OD600 levels compared to LCC and HiC (Figure 4C).

For all three libraries, we extracted and sequenced plasmids from the 10 to 14 clones showing the best combination of supernatant esterase activity and OD600. For all three enzymes, we observed recurring hits (*i.e*., SPs identified more than once across the sequenced set), and the SP hits across the three libraries differed from each other, indicating that the assay was specific for each PET hydrolase and that the libraries did not suffer from substantial cloning bias towards certain SPs. For LCC, out of 12 sequenced clones one SP was identified in 3 clones (ispe, the SP from IsP) and three SPs were identified twice (flid, efeb, and uxpb). For HiC, the dominant SP with 5 hits out of 10 sequenced clones was bglx. For IsP, out of 14 sequenced clones one SP was identified three times (lasb) and two SPs were identified twice (copb and agde) (Supplementary Table 4).

We next tested the most promising SP-PET hydrolase constructs (featuring all SPs identified multiple times per library) in shake flask cultures by monitoring PET hydrolase activity and OD600 over time. For all LCC and HiC constructs, we observed strong increases in supernatant and cell-associated esterase activity over the first 4 h following rhamnose induction. One of the best variants, bglx-HiC, showed an esterase signal in the supernatant of 1.71 U/mL 4 h post-induction, a 114-fold increase over the control. However, for both LCC and HiC, this was again accompanied by a subsequent collapse of the bacterial cultures, similar to the growth defects observed for the membrane-displayed constructs (Figure 4D and Supplementary Figure 15). For both LCC and HiC, we also observed a strong cell-associated esterase signal; by comparing the esterase activity of secreted and cytosolic PET hydrolase expression in culture supernatant, cell fraction, and cell lysate, we concluded that this is likely an artefact of PET hydrolase expression in the secretory pathway (Supplementary Figure 16). For IsP, we observed comparably small increases in supernatant esterase activity (either due to low IsP secretory expression or low activity of the enzyme in the pNPB assay) with the best variant, lasb-IsP, reaching 0.1 U/mL 4 h post-induction, 6.5-fold above the control. Importantly, the OD600 of all IsP expression cultures remained relatively stable (Figure 4D and Supplementary Figure 15).

In summary, our strategy of screening a 30-mer SP library resulted in the identification of SPs successfully directing the secretion of each of the three PET hydrolases. However, for LCC and HiC, secretory construct expression again resulted in cellular growth defects. For IsP, we observed only low esterase activity in the supernatant, but cell fitness remained high.

### Fine-tuning PET hydrolase expression with heat-inducible promoters

Since both surface display and secretion of LCC, HiC and IsP by our *P. putida* chassis was achievable but often incurred fitness costs, we turned our attention towards mitigating this problem. We reasoned that in both cases a likely culprit is an overload of the cellular machinery for protein folding, protein membrane insertion or protein translocation across inner and outer membranes.^66,67^ Thus, reducing PET hydrolase expression levels may result in improved cellular fitness.

As we observed for P_rhaB_, promoter systems relying on chemical inducers often show close to an all-or-none response, cannot be easily switched off again, or require genetic modifications to achieve a finely tunable expression response.^68,69^ Thus, we turned our attention to temperature-inducible promoters, where expression can be precisely steered by the intensity and duration of a heat shock. Using such a tightly controllable physical stimulus has the additional advantage of not requiring any chemical inducers, which could potentially be metabolized by the cells and thereby confound plastic growth assays.

We tested two heat-inducible promoter systems, both cloned into the RK2 low copy number plasmid. First, we used the cI857/P_L_ system, which is frequently used in *E. coli* and was recently shown to function efficiently in *P. putida*.^70^ The system consists of the P_L_ promoter blocked by the constitutively expressed heat-labile cI857 repressor, which is inactivated at elevated temperatures (ideally, above 40°C), thereby allowing transcription from P_L_. We designed this system as previously described,^70^ adding the BCD2 linker at the 5’ end preceding the PET hydrolase surface display or secretion cassettes (Figure 5A). Second, we used a short transcriptional-translational control unit natively regulating expression of the heat shock protein IbpA in *P. putida* and *P. aeruginosa*.^71^ This system consists of a promoter under control of the alternative heat shock sigma factor σ^32^. Furthermore, the *IbpA* mRNA 5’ UTR forms an RNA thermometer (RNAT) consisting of two hairpins that sequester the ribosome binding site. Upon heat shock (ideally, above 37°C), the hairpins melt, releasing the RBS and enabling initiation of translation. We utilized the IbpA unit of *P. aeruginosa*, providing 3-fold lower expression upon induction compared to IbpA from *P. putida*. We further introduced a 5’ UTR mutation (C40A) shown to further lower basal expression and provide a 10-fold induction upon heat shock,^71^ and mutated the last three nucleotides preceding the start codon to introduce a unique NdeI restriction site (Figure 5B). The BCD2 coupler was omitted, as its two strong RBSs would interfere with this system. The entire resulting transcriptional-translational regulator has a length of only 115 bp. In the following, we refer to this modified *P. aeruginosa* P_Ibpa_-5’ UTR construct as P_IbpA_. Notably, the mere presence of the three promoter systems (P_rhaB_, P_L_ or P_IbpA_) on empty vector control plasmids did not influence *P. putida* fitness (Supplementary Figure 17).

**Figure 5:**
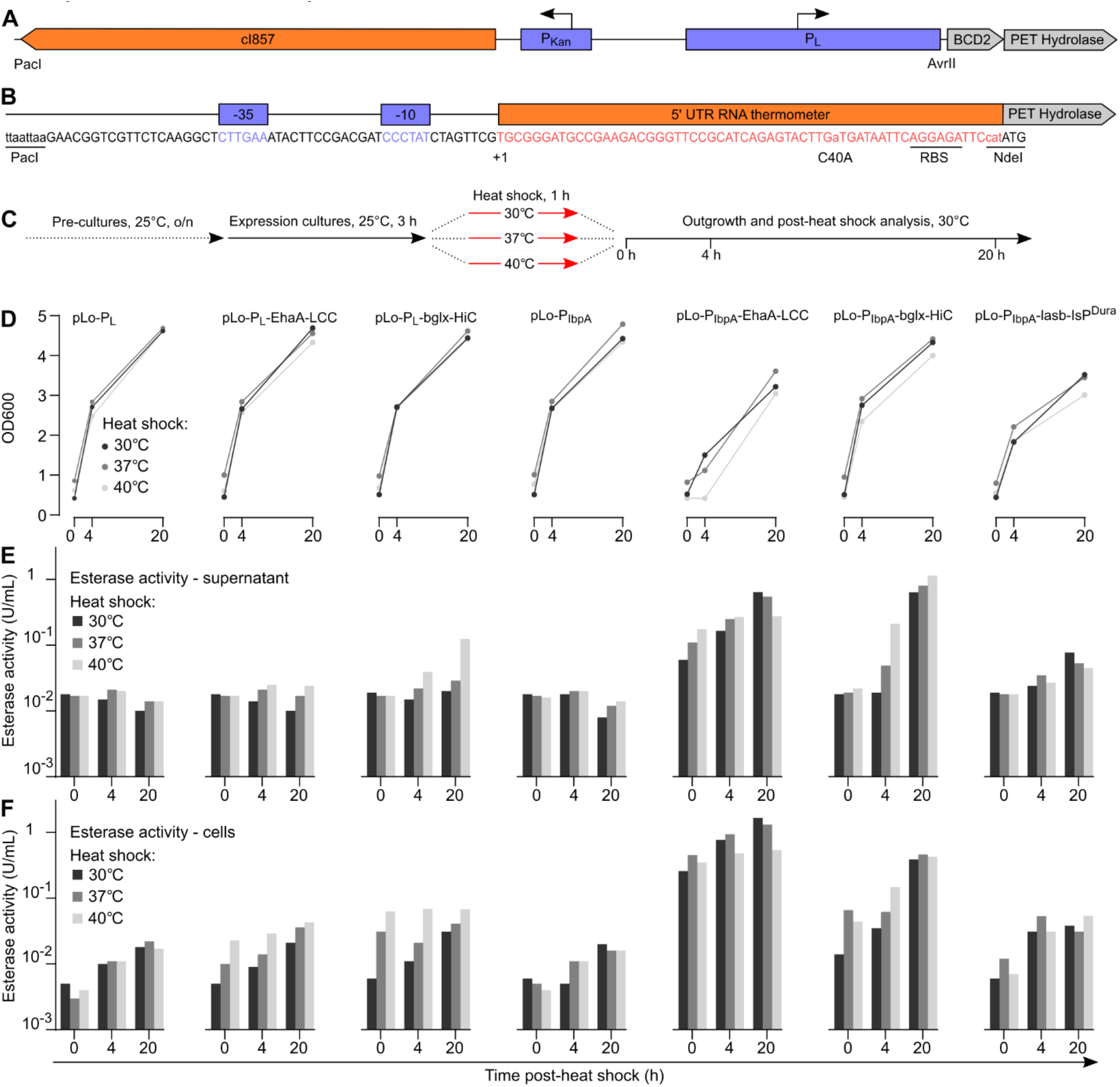
Testing P_L_ and P_IbpA_ heat shock inducible promoters. (A) Overview of the CI857/P_L_ system; this transcriptional regulatory unit spans 1410 bp from the PacI to the AvrII site. (B) Sequence overview and relevant features of the P_IbpA_ system used here. (C) Experimental protocol to test PET hydrolase expression from P_L_ and P_IbpA_ upon heat shock. (D) OD600 of cultures expressing PET hydrolases in dependance on the heat shock regime. (E) Esterase activity of cultures expressing PET hydrolases in dependance on the heat shock regime; top, esterase activity in the supernatant; bottom, cell-associated esterase activity. Data shown are derived from one representative experiment with duplicate cultures.

We initially tested P_L_ and P_IbpA_ with the two most promising PET hydrolase expression constructs: the surface display construct EhaA-LCC and the secreted bglx-HiC construct. For P_IbpA_ only, we also tested the secreted lasb-IsP, but substituted for IsP the recently described, more thermostable DuraPETase (in the following referred to as IsP^Dura^).^21^ We reasoned that this variant may tolerate the necessary heat shocks better while also showing enhanced catalytic activity on PET. To first test the expression behavior of the constructs, we subjected *P. putida* TA7-EG transformed with the plasmids to 1 h heat shocks at 30°, 37° or 40°C, followed by a 20 h outgrowth phase at 30°C during which we measured culture OD600 and esterase activity in the supernatant and the cell fraction (Figure 5C).

For P_L_, all cultures regardless of the heat shock regime showed good cell growth (Figure 5D). However, we also observed relatively low PET hydrolase expression levels, in agreement with previous work reporting longer and higher heat shocks required for optimal expression from P_L_.^70^ For pLo-P_L_-EhaA-LCC, we recorded a 2.5-fold induction of cell-associated esterase activity 20 h after the 40°C heat-shock, while we observed a 9-fold induction of supernatant esterase activity for pLo-P_L_-bglx-HiC under the same conditions (Figure 5E, F).

For P_IbpA_, we observed mild growth defects for EhaA-LCC and lasb-Isp^Dura^, regardless of heat shock regime, while bglx-HiC showed unimpaired cell growth. At the same time, we observed good PET hydrolase expression (Figure 5E, F). For pLo-P_IbpA_-EhaA-LCC, we observed a 104-fold induction of cell-associated esterase activity 20 h after the 30°C heat shock; interestingly, this was 3-fold higher than the esterase activity following the 40°C heat shock. For pLo-P_IbpA_-bglx-HiC, we recorded an 80-fold increase in supernatant esterase activity 20 h after the 40°C heat shock, with induction levels for the 30°C and 37°C heat shocks just slightly lower. For pLo-P_IbpA_-lasb-IsP^Dura^, we observed a 5-fold induction of supernatant esterase activity. Of note, across all three P_IbpA_ constructs, it appeared that maintaining the culture at 30°C (without a dedicated heat shock at 37°C or 40°C) already resulted in relatively high expression levels. Overall, these data pointed towards a promising function of P_Ibpa_ for achieving good PET hydrolase expression while maintaining cell fitness, especially for the secreted bglx-HiC construct.

Next, we further defined the temperature-dependent expression behaviour of P_IbpA_. Cultures of *P. putida* TA7-EG transformed with empty vector or pLo-P_IbpA_-bglx-HiC were shifted from 25°C to either a constant expression temperature of 30°C or a heat shock at 37°C for 2, 4, 6, or 20 h, followed by an outgrowth phase at 30°C; all cultures were analyzed 20 h after the start of the heat shock (Figure 6A). We observed a constant increase in supernatant esterase activity from 0.85 U/mL to 1.6 U/mL (empty vector: 0.025 U/mL) from the constant 30°C culture to the 6 h heat shock culture, while the 20 h heat shock culture showed 2.5 U/mL (Figure 6). However, the 20 h heat shock culture also showed a strongly reduced OD600, indicating that a large part of this supernatant esterase activity may originate from lysed cells. For the 6 h heat shock culture we observed a mildly decreased OD600 value, while the 0 h, 2 h and 4 h cultures showed a growth behavior just slightly below empty vector controls. Thus, it appeared that shifting *P. putida* cultures from 25°C to 30°C is sufficient to induce good expression from P_IbpA_, while a 2 to 4 h heat shock at 37°C provides an additional small boost to expression levels without affecting cell fitness.

**Figure 6:**
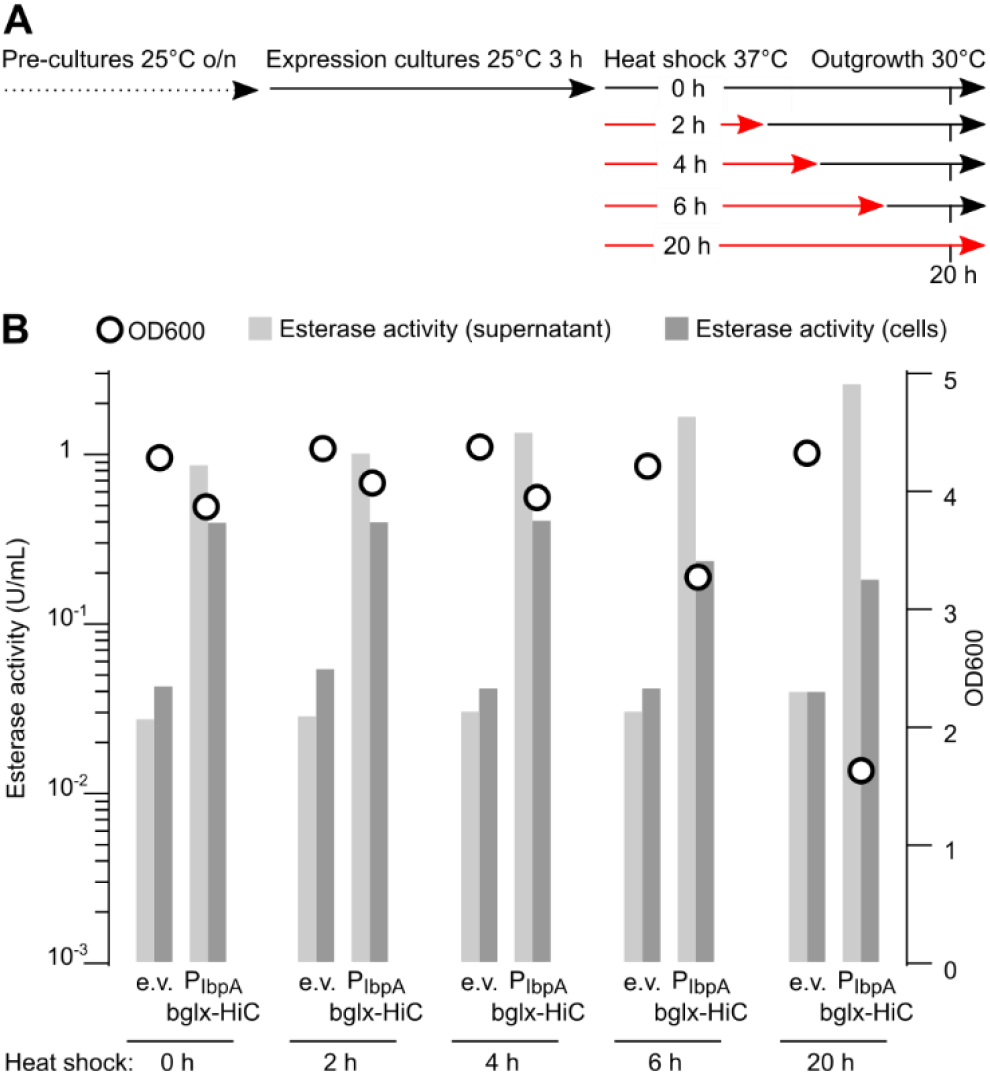
Defining P_IbpA_ temperature-dependent expression output. (A) Experimental protocol to test PET hydrolase expression from P_IbpA_ upon different heat shock regimes. (B) OD600 and esterase activities of *P. putida* TA7-EG cultures transformed with pLo-P_IbpA_ (empty vector, e.v.). or pLo-P_IbpA_-bglx-HiC in dependance on the heat shock regime. Data shown are derived from one representative experiment with duplicate cultures.

### Genomic integration of P_IbpA_ PET hydrolase expression constructs

In a final engineering step, we genomically integrated the P_IbpA_-PET hydrolase expression constructs into our *P. putida* strains. We reasoned that this may lessen expression burden by reducing construct copy number to one, may lead to better culture stability as potential plasmid loss is avoided, and circumvents constant antibiotic selection for plasmid maintenance, all factors that may contribute to better plastic degradation performance of long-term cultures. We chose the pBAMD system for genomic integration and constructed plasmids encoding integration cassettes for P_IbpA_-EhaA-LCC, P_IbpA_-bglx-HiC, and P_IbpA_-lasb-IsP^Dura^. The cassettes were knocked into *P. putida* KT2440 and strains TA7-EG and TA7-BD. The best-performing clones were whole-genome sequenced to identify the expression cassette’s integration site and to ensure absence of undesired mutations (Supplementary Tables 1, 2).

First, we investigated the induction behavior of the genomically integrated P_IbpA_ constructs, using *P. putida* KT2440 and the corresponding P_IbpA_-EhaA-LCC, P_IbpA_-bglx-HiC and P_IbpA_-lasb-Isp knock-in strains. Shake flask cultures in LB medium were subjected to constant 30°C or heat shocks at 37°C for 2 or 6 h, followed by a recovery phase at 30°C and subsequent analysis of OD600 and esterase activity (Figure 7A). As for plasmid-based expression, we observed the best supernatant esterase signal for P_IbpA_-bglx-HiC, showing a 4-fold induction over *P. putida* KT2440 for the constant 30°C culture, 16-fold for the 2 h heat shock, and over 60-fold (up to 0.95 U/mL) for the 6 h heat shock, while maintaining growth comparable to the wt control (Figure 7B). For P_IbpA_-EhaA-LCC and P_IbpA_-lasb-Isp^Dura^, we observed lower induction levels (2 to 4-fold over wt) while also recording slightly reduced OD600. Similar results were obtained for the equivalent P_IbpA_-PET hydrolase knock-ins in strains *P. putida* TA7-EG and TA7-BD (data not shown).

**Figure 7:**
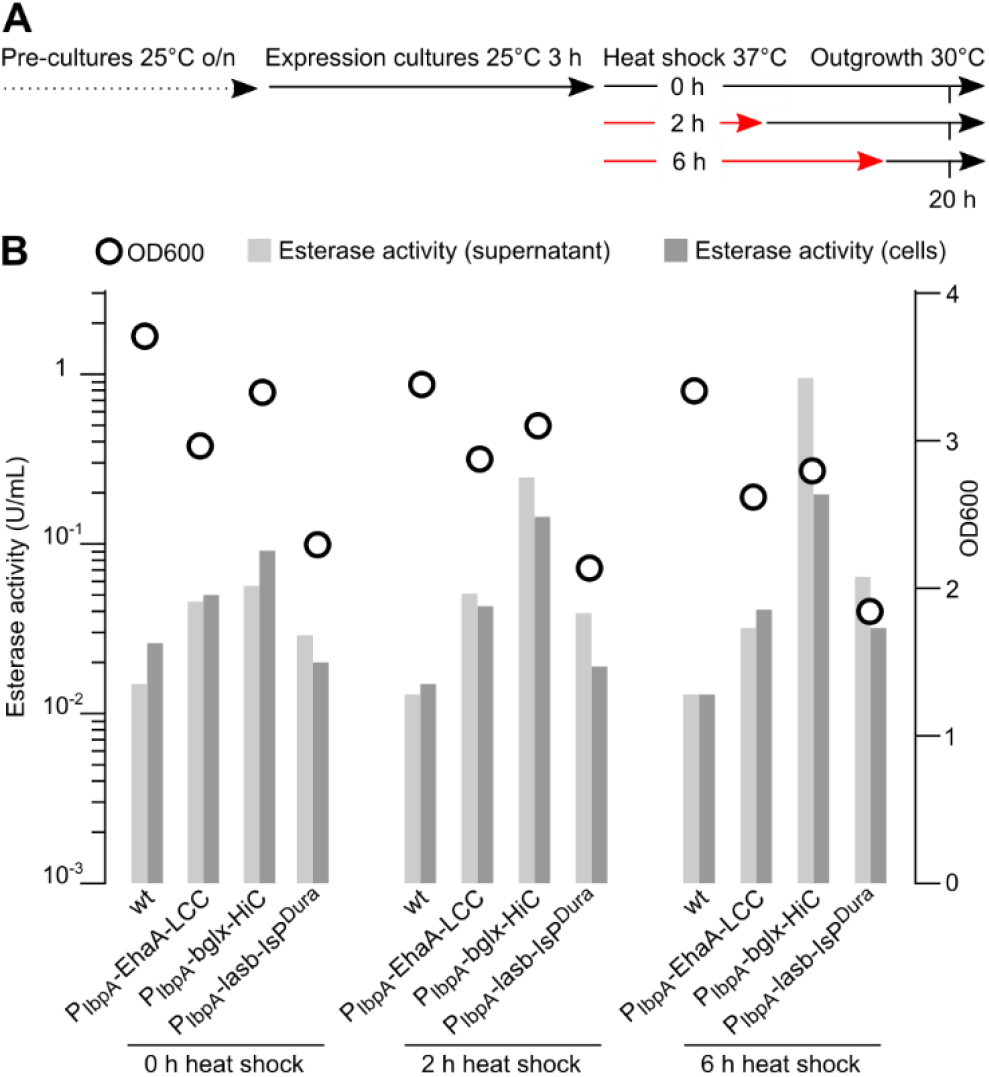
Testing expression of *P. putida* KT2440 P_IbpA_-PET hydrolase knock-in strains. (A) Experimental protocol to test the expression of genomically integrated P_IbpA_-PET hydrolase constructs upon different heat shock regimes. (B) OD600 and esterase activities of *P. putida* KT2440 (wt) or the corresponding P_IbpA_-EhaA-LCC, P_IbpA_-bglx-HiC, and P_IbpA_-lasb-IsP^Dura^ knock-in strains in dependance on the heat shock regime. Data depict one representative experiment with duplicate cultures.

We confirmed secretory HiC expression from the plasmid-based pLo-P_IbpA_-bglx-HiC construct by SDS-PAGE and LC-MS analysis (Supplementary Figure 18). However, SDS-PAGE was not sensitive enough to unambiguously confirm secretory expression of HiC from the genomically integrated P_IbpA_-bglx-HiC cassette or plasmid-based expression of IsP from pLo-P_IbpA_-lasb-IsP^Dura^. This underscores the high secretory activity seen for the bglx-HiC construct, as well as the desired effect of modulating expression levels through plasmid-based versus genomic expression.

Next, we directly compared absolute PET hydrolase expression levels across constructs, using the surface display construct EhaA-LCC (Supplementary Figure 19). EhaA-LCC expression from pLo-P_rhaB_-EhaA-LCC yielded a cell-associated esterase activity of 1.4 U/mL (100-fold over empty vector), measured 3 h post-induction (before culture collapse). EhaA-LCC expression from pLo-P_IbpA_-EhaA-LCC after a 2 h, 37°C heat shock followed by outgrowth at 30°C for 18 h delivered 0.26 U/mL of cell-associated esterase activity, 5-fold lower than the expression output from pLo-P_rhaB_-EhaA-LCC. EhaA-LCC expression from the genomically integrated P_IbpA_ construct under the same induction regime delivered 0.08 U/mL, 18-fold lower compared to plasmid-based P_rhaB_-dependent expression and 3-fold lower than plasmid-based expression from the same P_IbpA_ promoter. At the same time, while plasmid-based P_rhaB_ expression cultures collapsed a few hours after induction, plasmid-based P_IbpA_-EhaA-LCC cultures showed reduced but stable OD600, while the genomically integrated P_IbpA_-EhaA-LCC constructs showed an OD600 not far below the control strain. Thus, our engineering strategy succeeded in balancing PET hydrolase expression levels with cell viability, but at the cost of lower absolute PET hydrolase expression output.

A central feature of synthetic PETtrophy would be the bacterial ability to efficiently express PET hydrolases with only TA and EG (or TA and BD in case of PBAT) as carbon sources. Thus, we compared PET hydrolase expression of strains *P. putida* KT2440, TA7-EG and TA7-BD with the integrated P_IbpA_-bglx-HiC cassette in either LB medium or M9 minimal medium (Figure 8). Both overall cell growth (OD600) and esterase activity were dramatically reduced in M9 minimal medium, especially for strains TA7-EG and TA7-BD, where we recorded only 2-fold inductions of supernatant esterase activity (compared to ca. 30 to 50-fold induction in LB medium) over the control strains. Thus, PET hydrolase expression in minimal medium is detectable but strongly reduced, likely reflecting the lower energetic value of the polyester monomers compared to complete medium.

**Figure 8:**
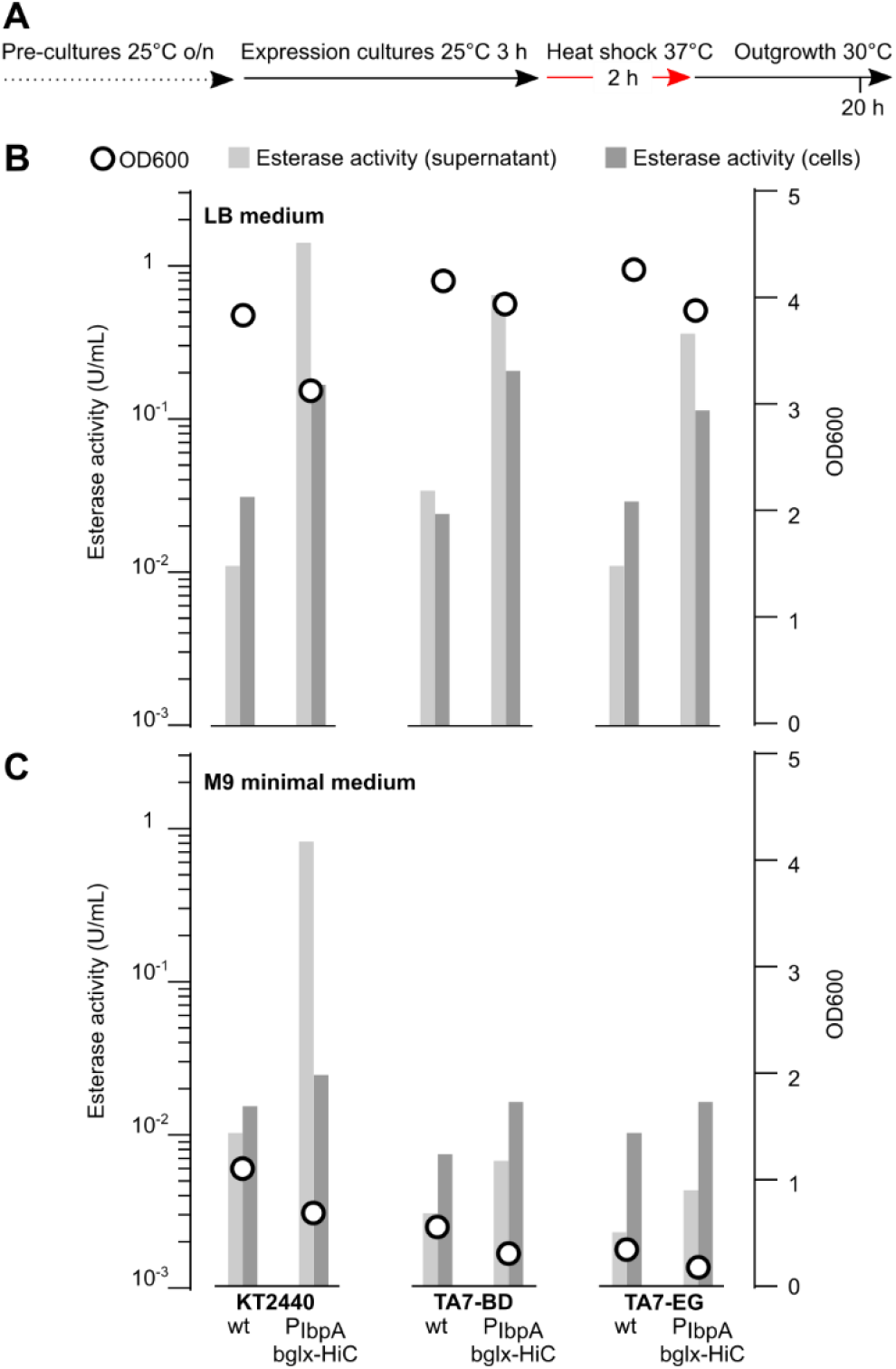
Comparing P_IbpA_-PET hydrolase expression in LB versus M9 minimal medium. (A) Experimental protocol to test genomically integrated P_IbpA_-bglx-HiC construct expression in different culture media. (B) OD600 and esterase activities of cultures in LB medium including *P. putida* KT2440, TA7-EG, and TA7-BD, either as the non-hydrolase expressing strains (labelled wt) or with the genomically integrated P_IbpA_-bglx-HiC cassette. (C) OD600 and esterase activities of *P. putida* KT2440, TA7-EG, and TA7-BD, either as the wt strains or with the genomically integrated P_IbpA_-bglx-HiC cassette, in M9 minimal medium with 20 mM citrate (KT2440), 10 mM TA + 10 mM BD (TA7-BD) or 10 mM TA + 10 mM EG (TA7-EG). Data depict one representative experiment with duplicate cultures.

### Assessing PET and PBAT degradation by P. putida P_IbpA_-PET hydrolase strains

In a final step, we assessed the capacity of our engineered strains to degrade PET or PBAT plastics and maintain synthetic PETtrophy. Plastic samples included amorphous PET film, commercial PET bottle chips, high-crystallinity PET powder, polyester garment fibers, and a commercial PBAT-based copolymer film. We tested different culture formats to bring plastic samples in contact with bacterial cultures, including static cultures in 14-mL tubes, 10-cm Petri dishes, or 6-well plates, and shaking cultures in 14-mL tubes or 125-mL flasks. Overall, we obtained the most promising results with 15-mL shaking cultures in 125-mL flasks at a shake rate of 75 rpm. We also tested different culture media, including LB medium (to assess maximal PET hydrolase expression and plastic degradation capacity), M9 minimal medium supplemented with 20 mM TA (to mimic high monomer supply through efficient plastic degradation), and M9 minimal medium without added carbon source (to test for synthetic PETtrophy). Typical PET hydrolase culture induction regimes as previously established for the P_IbpA_ promoter consisted of a 2 h heat shock at 37°C per day, with the remaining time at 25°C.

We maintained cultures for up to 6 weeks; in this timeframe, we were unable to reliably detect synthetic PETtrophy, *i.e*., the bacterial capacity to proliferate on polyester plastics as sole carbon source. We predominantly used OD600 measurements to assess bacterial growth, supplemented with visual culture inspection by eye, stereo microscopy, and optical microscopy. While we occasionally observed biofilm-like structures and live bacteria in cultures that were several weeks old (potentially stemming from the inoculum), this typically did not manifest as reliably detectable OD600 increases and did not result in reproducible weight loss of plastic substrates.

In contrast, in nutrient-supplemented cultures (either LB or M9-TA), we detected modifications of the plastic substrates. For PET, we typically observed only minor changes: for example, after six days of culturing in LB medium with strain *P. putida* KT2440 P_IbpA_-bglx-HiC, we observed fraying along the edges of an amorphous PET film and a weight loss in the range of 1.5%, neither of which was observed in control cultures of *P. putida* KT2440 (Supplementary Figure 20). For PBAT copolymer film, results were more pronounced: after six days of culturing in LB medium with strain *P. putida* KT2440 P_IbpA_-bglx-HiC, we observed fraying and tearing of the film and a weight loss of approximately 25%, neither of which was observed in control cultures of *P. putida* KT2440 (Figure 9A). Furthermore, in a 4-week culture in M9-TA medium with strain *P. putida* TA7-BD P_IbpA_-bglx-HiC, we observed a striking decay of the film to parts smaller than 5 mm alongside a weight loss in the range of 40% (Figure 9B, C). Scanning electron microscopy (SEM) of film samples showed surface erosion and development of cracks and fissures in the *P. putida* TA7-BD P_IbpA_-bglx-HiC treated samples, matching the observed increase in film brittleness and decay (Figure 9D, E). Despite the apparent activity of the secreted HiC hydrolase on the PBAT copolymer film, we did not detect sustained growth of *P. putida* TA7-BD P_IbpA_-bglx-HiC in M9 medium supplied with PBAT copolymer film as only carbon source. Potentially, this is due to the reported mediocre activity of HiC on PBAT oligomers, which may fail to supply TA and BD monomers for cellular catabolism.^72,73^ Indeed, we did not detect TA in the culture medium by high performance liquid chromatography, likely because the small amounts of potentially released TA were quickly consumed by the engineered strains. While these observations hint towards the polyester plastic degradation potential of our strains, they also highlight future work required to fine-tune and improve PET hydrolase expression levels, culture conditions, and cellular fitness

**Figure 9:**
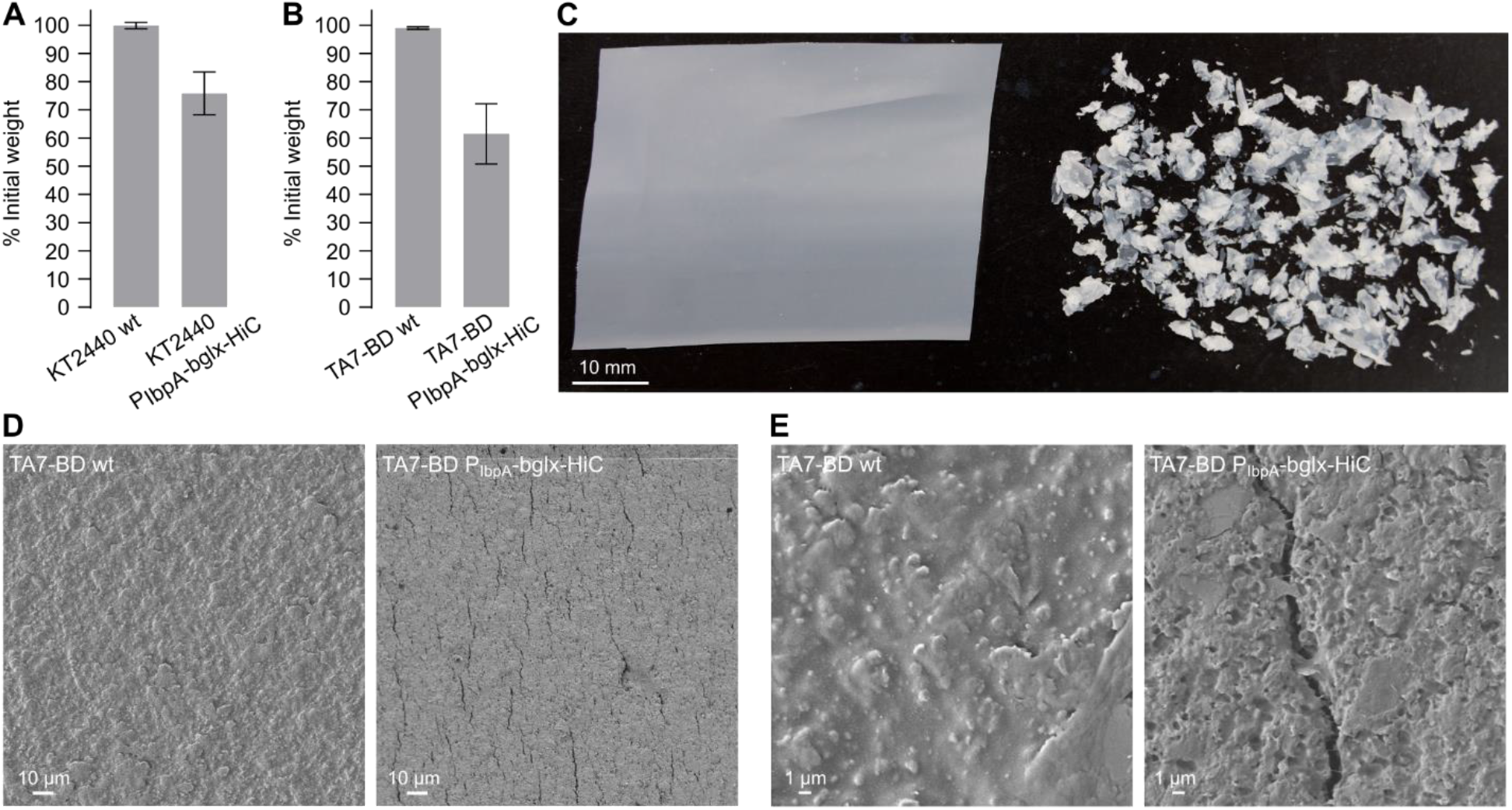
PBAT copolymer film degradation by engineered *P. putida* strains. (A) Weight loss of a PBAT copolymer film after 6 days of incubation in LB medium with *P. putida* KT2440 or *P. putida* KT2440 P_IbpA_-bglx-HiC. (B) Weight loss of a PBAT copolymer film after 4 weeks of incubation in M9 minimal medium supplemented with 20 mM TA with *P. putida* TA7-BD or *P. putida* TA7-BD P_IbpA_-bglx-HiC. (C) Image of the PBAT copolymer film after the treatment shown in (B), with the *P. putida* TA7-BD-treated film left and the *P. putida* TA7-BD P_IbpA_-bglx-HiC-treated film right. (D) and (E) SEM images of the same films shown in (C) at different magnifications.

## Discussion

In this work, we followed a two-tiered strategy towards endowing the bacterium *Pseudomonas putida* with the ability to utilize PET or PBAT plastics as sole carbon sources. First, we used metabolic engineering and adaptive laboratory evolution to allow *P. putida* to catabolize the PET or PBAT monomers TA, EG and BD. Second, we tested various expression systems to achieve stable and efficient secretion or membrane display of three PET hydrolases. The resulting monomer-metabolizing and PET hydrolase-expressing *P. putida* strains were tested in different culture conditions to assess their PET or PBAT plastic degradation capacity. While this approach yielded mixed results, and we did not, to date, observe the desired bacterial ‘synthetic PETtrophy’, we derived important insights into the factors governing (and limiting) the efficiency of a potential whole-cell biocatalytic system for polyester plastic degradation and assimilation.

The engineering of *P. putida* to allow catabolism of polyester monomers TA, EG and BD is an active field of research, and different groups recently disclosed approaches similar to the results reported here. Werner *et al*. utilized the same *tphII* operon from *Comamonas sp*. E6, in combination with a TA transporter (*tpaK*) from *Rhodococcus jostii*, to install TA metabolism in *P. putida*.^34^ Similarly, Narancic *et al*. recently transferred a homologous operon (including the TA transporter *tphK*) originating from the native TA metabolizer *Pseudomonas umsongensis* GO16 to *P. putida*, again enabling it to use TA as sole carbon source.^74^ Together, these studies and the results reported here firmly demonstrate that the *tphA1, A2, A3*, and *B* genes of *tph*-type operons are sufficient to enable TA catabolism by *P. putida*. Notably, our work differs from the other two studies by omitting a TA transporter from the genomic knock-in cassette. Instead, we found that one of the many native *P. putida* aromatic acid transporters can be repurposed to function in TA uptake with just a single amino acid mutation. The resulting putative TA transporter mhpT^L374V^ may prove useful for future engineering efforts, as native expression of mhpT^L374V^ may come with a lower fitness cost than the expression of a heterologous membrane transporter. Additional work is needed to unravel the mhpT^L374V^ TA uptake mechanism and its regulation of expression (*e.g*., the role of the R51C mutation in the marR-family transcription factor PP_3359).

For EG, previous research has firmly established that *P. putida* contains the metabolic genes required for its utilization; however, the two key enzymes Gcl and GlxR are transcriptionally repressed by GclR.^45,46^ Li *et al*. recently showed that ALE rapidly results in GclR loss-of-function mutations, thereby enabling efficient utilization of EG as sole carbon source by *P. putida*.^47^ Our observation of *P. putida* strains carrying a likely GclR E130* loss-of-function mutation after ALE for EG utilization agrees with these studies. Similarly, Li *et al*. recently demonstrated that ALE can enable BD use as sole carbon source by *P. putida*.^48^ The underlying metabolic network is less clearly understood; however, apparent key mutations occur in the transcription factor PP_2046, which likely regulates expression of the PP_2047-2051 operon involved in BD metabolism.^48^ In line with this, upon performing ALE with *P. putida* TA7 for BD utilization, we obtained a strain carrying a mutation, V136A, in PP_2046. We did not attempt to engineer metabolism of adipic acid, the second PBAT aliphatic monomer; however, Ackermann *et al*. recently demonstrated metabolic engineering to allow *P. putida* to metabolize this compound as well.^75^ In summary, this body of work highlights the potential of *P. putida* as a bacterial chassis for polyester plastic monomer metabolization and upcycling, as currently pursued by several research groups.

To install polyester plastic depolymerization capabilities into our *P. putida* strains, we explored the heterologous expression of *Ideonella sakaiensis* PETase (IsP),^26^ *Humicola insolens* cutinase (HiC),^60^ and leaf compost cutinase (LCC).^61^ These hydrolases are among the best-characterized PET-depolymerizing enzymes, originate from bacteria and fungi, and show different optimum temperatures. Thus, we reasoned that by testing this diverse set of enzymes we would maximize our chances of achieving stable expression and catalytic function in our *P. putida* strains. Both LCC and HiC have been described to catalyze the complete depolymerization of PET to TA and EG. On the other hand, IsP produces only small amounts of TA and predominantly releases the intermediate PET degradation product mono(2-hydroxyethyl) terephthalate (MHET), which is subsequently hydrolyzed to TA and EG by the *I. sakaiensis* auxiliary enzyme MHETase.^26^ Interestingly, recent studies of IsP expression in green algae found that, upon prolonged incubation, TA accumulates in PET degradation cultures.^62,76^ However, it is currently unclear whether this MHET degradation activity originates from IsP or from unknown proteins of the microalgal hosts. In this study, we chose to express IsP without MHETase to establish baseline conditions for stable IsP expression and to facilitate direct comparisons with HiC and LCC.

To explore PET hydrolase expression, we chose an approach of rapid design-build-test-learn cycles. To accommodate this strategy, we relied on assessing the activity of expressed PET hydrolases with the spectrophotometric surrogate assay of para-nitrophenyl butyrate (pNPB) cleavage. While this assay offers fast results and is easy to scale, IsP shows markedly lower activity on pNP alkanoate esters than do HiC and LCC.^26^ Thus, without precise enzyme quantification (which was not feasible for the many small-scale expression cultures typically performed here), it was occasionally unclear whether the low pNPB esterase activity of IsP expression cultures was due to low enzyme expression or low IsP activity in the pNPB assay. Regardless, the pNPB assay delivered detailed insights into PET hydrolase expression levels, especially since *P. putida* cultures (including culture supernatant, whole cells, and cell lysate) showed negligible pNPB esterase activity in the absence of heterologous PET hydrolase expression.

Our analyses of P_rhaB_-driven PET hydrolase membrane display showed that EhaA and OprF membrane anchors yielded strong PET hydrolase cell surface expression but also resulted in subsequent culture collapse. The latter was not entirely surprising, as heterologous expression of membrane proteins frequently results in compromised cellular fitness.^66,67^ Indeed, the developers of the *P. putida* EhaA display system (in their study with an esterase cargo and under control of the araBAD promoter) reported a 1000-fold reduction in cell viability following induction of expression.^49^ Furthermore, they allowed expression to proceed for only 2 h before cell analysis; we speculate that the authors also encountered impaired cell integrity at later timepoints post-induction. The developers of the *P. putida* OprF display system (in their study with a lipase cargo and under control of the tac promoter) did not report cellular growth defects following induction; however, we note that the authors used a relatively low IPTG inducer concentration (0.1 mM) and, again, a short expression window of only 4 h before cell analysis.^52^ Neither of the two studies reported bacterial growth curves (OD600) following induction.

We next explored secretory expression of the three PET hydrolases, using a library comprising 30 SPs, and succeeded in identifying SPs directing extracellular secretion of each of the three enzymes. As the joint effects of SP and enzyme cargo on secretion efficiency are incompletely understood,^63,64^ we believe that the library screening approach performed here was vital to identify suitable SPs for each PET hydrolase. Notably, the oligo library we used to clone the SP library came at a synthesis cost of only around 100 USD; thus, we believe that a screening strategy (as opposed to individual cloning and testing of selected SPs) represents a cost-effective and fast strategy to identify SPs for a given PET hydrolase (or other gene of interest), especially if a fast read-out such as the pNPB assay is available.

As with surface-displayed PET hydrolases, we observed impaired cell growth following secretion construct expression. Notably, this was more pronounced for HiC and LCC, while IsP secretion resulted in relatively mild growth defects. While we assessed cell fitness in the initial SP library screen and validated the best SP-PET hydrolase constructs in shake flask cultures, there is a small possibility that our screen did not identify the SPs resulting in the most efficient enzyme secretion, but rather those SPs inducing the fastest cell leakage. Thus, it would be interesting to repeat the entire SP library screen with the P_IbpA_ or P_L_ promoter systems instead of the strong P_rhaB_ promoter, to determine if a lower, more sustainable construct expression output would result in the same SPs coming out on top.

To ameliorate the problem of cell toxicity following PET hydrolase expression, we next substituted the P_rhaB_ promoter with the heat-inducible CI857/P_L_ and P_IbpA_ promoter systems. In agreement with the literature, P_IbpA_ was largely inactive at 25°C but provided low expression at 30°C and strong expression at 37°C, while cI857/P_L_ required induction at ≥ 40°C.^70,71^ Thus, we chose to proceed with P_IbpA_ due to its induction range within the typical growth temperature of *P. putida*, easily clonable size, *Pseudomonas* origin, and absence of auxiliary transcriptionally active proteins.

In a final engineering step, we integrated the most promising PET hydrolase expression constructs under control of the P_IbpA_ promoter into the genome of our engineered *P. putida* strains. The resulting *P. putida* P_IbpA_ knock-in strains showed good PET hydrolase expression upon induction in complex medium, as well as good viability, likely due to the reduction in expression construct copy number to a single genomic copy. On the other hand, we observed only limited PET hydrolase expression from cultures in minimal medium containing TA, EG or BD as sole carbon sources. We believe that this imperfect balance between nutrient supply and PET hydrolase expression was a decisive factor in the lack of bacterial growth on plastics as sole carbon source, as discussed further below.

Across a wide range of test cultures, including different PET or PBAT plastic samples and growth conditions, we were unable to detect bacterial PETtrophy, that is, the envisioned capacity of our strains to depolymerize polyester plastics while maintaining cell growth and PET hydrolase expression. The underlying reasons are likely manifold. First, the low culture temperature, in the range of 30-37°C, likely poses a strong limitation, especially for HiC and LCC, given the much higher activity of these enzymes (and a higher absolute activity compared to IsP) at temperatures closer to the glass transition temperature of PET. For IsP-expressing strains, we would have expected to see better activity, especially on amorphous PET; however, the omittance of MHETase may have compromised the efficiency of our strains. Furthermore, despite our promoter-tuning efforts, our strains likely still do not strike an ideal balance between PET hydrolase expression levels and the available monomer concentrations as released by PET hydrolase-catalyzed plastic depolymerization. This represents a chicken-and-egg problem: PET hydrolase expression requires plastic monomers as carbon sources; however, monomer production requires a certain minimum PET hydrolase activity. This problem may be exacerbated by the comparatively low yield of central carbon metabolites obtained from PET monomers, as compared to typical sugar feedstocks or monomers of other plastics.^8^ In this context, it is also interesting to point out that *Ideonella sakaiensis* PET degradation cultures contained low amounts (0.01%) of yeast extract.^26^ This may have served precisely the purpose outlined above, namely, to maintain a basal level of cellular metabolism ensuring PETase and MHETase expression. Lastly, we point out that we employed commonly used but rather simplistic methods to assess bacterial PETtrophy, such as measurements of culture OD600 and plastic weight loss. It has been argued that such methods are unsuited for assessing biocatalytic plastic degradation, and indeed, more fine-grained techniques capable of detecting subtle changes in plastic composition or bacterial growth rate are available.^77,78^ It would be interesting to re-assess the plastic degradation and growth performance of our strains using more sensitive techniques; however, at the same time, we argue that if such methods are necessary to detect bacterial growth and plastic degradation, the relevance of such a bacterial system for delivering tangible solutions to the plastic waste problem is likely small.

How does our work compare to recent studies aiming to create whole-cell biocatalysts for PET plastic degradation? To date, *I. sakaiensis* remains the only bacterium with thoroughly characterized biochemical machinery capable of sustained growth on PET substrates, albeit with the caveat of requiring small amounts of extraneously added complex medium. In addition, several recent studies aimed to create whole-cell biocatalysts for PET biodegradation, complementing the more traditional approach of using purified PET hydrolases. In most cases, these studies aimed for recombinant PET hydrolase secretion or membrane display by microbial hosts; unfortunately, most of these studies lack detailed information on the fitness of the resulting recombinant strains, which proved to be a key factor in our work. Seo *et al*. demonstrated IsP secretion in *E. coli* by testing five different *E. coli* SPs; no information on the growth characteristics of the recombinant strains was provided.^79^ Similarly, Shi *et al*. expressed IsP in *E. coli* with the pelB SP and variants thereof; while they do not comment on cell growth, we note that the supernatant of their expression strain shows a much higher overall concentration of proteins on SDS-PAGE compared to the empty vector control strain, which we interpret as indication of cell leakage or lysis.^80^ Cui *et al*. also expressed IsP in *E. coli* using short ‘enhancer’ peptides fused to standard SPs; they provide culture OD600 for their expression strains, but lack crucial empty vector or uninduced controls.^81^ Huang *et al*. demonstrated IsP secretion by the commonly used gram-positive secretion host *Bacillus subtilis* using the native IsP SP; growth curves of recombinant strains were provided and showed no significant growth defects compared to controls.^82^ Similarly, Wang *et al*. showed secretory expression of IsP in *B. subtilis*, using a weak constitutive promoter and testing five different SPs; they additionally performed bacterial growth tests at 22°, 28° and 37°C and documented culture OD600, but unfortunately lacked an empty vector control strain for comparison.^83^ Moog *et al*. showed secretory expression of IsP^R280A^ in the microalgae *Phaeodactylum tricornutum*, using a *P. tricornutum* SP. While not providing growth curves of the recombinant strains, they controlled for cell lysis (and reported none) by measuring the cytoplasmic marker alpha-tubulin in the culture supernatant.^62^ They report PET and PETG degradation by the microalgal cultures under certain conditions, but also provide the valuable information that several degradation setups were unsuccessful, possibly due to low reaction temperatures (21-26°C), culture agitation, or different characteristics of the PET substrates. Kim *et al*. also expressed IsP in a green algae, *Chlamydomonas reinhardtii*, but used an intracellular expression system necessitating cell lysis for PET degradation assays; they do not provide growth curves of their recombinant strains.^76^ Yan *et al*. showed secretory expression of LCC in the thermophilic bacterium *Clostridium thermocellum*, using a *C. thermocellum* SP; they show a growth curve for their engineered strain, but also lack appropriate controls.^84^ Notably, they also report that LCC is only actively produced in the first day of a 14-day culture due to subsequent medium depletion. Regarding membrane display, Chen *et al*. displayed IsP on the surface of the yeast *Pichia pastoris*, testing three *P. pastoris* glycosylphosphatidylinositol-anchored proteins as display system; they report similar growth of their recombinant strains compared to a wt control.^85^ Lastly, Heyde *et al*. recently explored IsP surface display on *E. coli*, using the Lpp-OmpA or C-IgAP display systems; they do not provide information on cell fitness following construct expression.^86^

Thus, while secretion or membrane display of PET hydrolases is a growing field, there is unfortunately often only very limited information on the effects of such expression systems on host fitness (a trend we observe across the general protein secretion or membrane display literature). While host fitness may not be relevant for many biochemical applications, such information is crucial to establishing continuous bioprocessing-like applications as aimed for in this study. Furthermore, if host growth characteristics are not controlled for, especially in enzyme secretion systems, one may confound the accumulation of the target protein in the culture supernatant as ‘secretion’, when it may originate instead from cell leakage or lysis. Thus, we strongly advocate for assessment and reporting of host fitness for any newly developed enzyme secretion or membrane display system, including comparison to the matched empty vector or wt strain, to enable scientists aiming to establish CBP-like processes to gauge the suitability of each system for their purposes.

Finally, it remains to be discussed whether pursuing a CBP-like process using a mesophilic bacterium is a promising strategy. A PET CBP process would need to incorporate both PET degradation through hydrolase expression and efficient monomer metabolization, of which the former is likely the more challenging part. We chose to use *P. putida* due to the ease of engineering PET monomer metabolization, and subsequently tackled PET hydrolase expression. However, given the strict relationship between reaction temperature and PET hydrolase degradation performance, it may be promising to first establish PET hydrolase expression in a thermophilic host, and subsequently engineer it for monomer metabolism.^84^ Recent progress in the development of genetic tools and knowledge for genetic engineering of thermophiles will undoubtedly prove useful for such endeavors.^87,88^ Nevertheless, we argue that a mesophilic, self-sustaining PET degrading bacterium as pursued here may still have significant value. First, by creating such an organism, we may learn about the determinants of bacterial growth on PET in mesophilic environments. Second, such organisms may prove useful for approaches in which speed of degradation is less decisive, for example for small-scale, decentralized PET waste composting facilities. In this regard, it would be very interesting to perform economic process modelling analyses for a putative whole-cell-based PET recycling process, as recently performed by Singh *et al*. for a recycling process based on purified enzyme,^9^ to better gauge the potential merits of establishing a whole-cell PET degradation biocatalyst. We look forward to seeing how the next generation of highly efficient PET hydrolases, such as DuraPETase,^21^ FastPETase,^20^ or the LCC ICCG variant,^19^ may be leveraged in the future to develop whole-cell biocatalytic strategies as novel solutions for the PET waste problem.

## Conclusions

What does it take to engineer a mesophilic bacterium for growth on polyester plastics? Here, we addressed this question using the industrial workhorse organism *Pseudomonas putida*. Together with other recent studies, our work shows that polyester plastic monomer metabolism is readily achievable using genetic engineering and adaptive laboratory evolution. On the other hand, attaining high-level extracellular expression and activity of PET hydrolases remains a challenge. Our work of testing a variety of expression systems to balance host cell fitness with PET hydrolase expression may prove useful for future efforts aiming to establish extracellular enzyme expression in *P. putida*, and in particular for efforts geared towards establishing synthetic PETtrophy in mesophilic bacterial hosts.

## Materials and Methods

### Data availability

The DNA sequencing read data is deposited in the NCBI Sequence Read Archive under BioProject ID PRJNA808732. The sequence of pBAMD-tphII has been deposited in GenBank (accession number ON192369). All other plasmid maps and sequences are available on request.

### General reagents, procedures, and bacterial culturing conditions

Bacterial strains, plasmids, and DNA oligos used in this study are listed in Supplementary Tables 5, 6, and 7. *Escherichia coli* and *Pseudomonas putida* strains were routinely grown in LB medium (10 g tryptone, 5 g yeast extract, 5 g NaCl per liter ddH_2_O, pH 7.0) at 37°C or 30°C, supplemented with antibiotics where needed (Ampicillin (Amp): 100 μg/mL; Kanamycin (Kan): 50 μg/mL). Adaptive laboratory evolution and assays with *P. putida* strains for growth on terephthalic acid, ethylene glycol, 1,4-butanediol, PET, and PBAT, were performed in modified M9 minimal medium: 47.7 mM Na_2_HPO_4_7H_2_O, 22 mM KH_2_PO_4_, 18.7 mM NH_4_Cl, 8.5 mM NaCl, 2 mM MgSO_4_, 0.1 mM CaCl_2_, 1x trace metals (Teknova 1000x trace metals mixture), and varying pH (adjusted with NaOH or HCl) depending on the experiment. Bacteria were routinely cultured in 125 mL shake flasks (15 mL working volume) or 14 mL snap-cap tubes (2-4 mL working volume) at 250 rpm. Bacterial culture plates were prepared with 1.5 % (w/v) agar. Expression of proteins under control of the P_rhaB_ promoter was induced by addition of a 1 M sterile rhamnose stock solution in ddH_2_O to 2 mM final concentration, unless otherwise indicated. Optical density (OD600) of cell cultures was measured in a Beckman-Coulter DU520 spectrophotometer. Bacterial growth in clear round-bottom 96-well plates was measured in a Biotek Synergy plate reader with constant shaking (set to the fastest shake rate, no rpm provided) at 30°C, 100 μL volume per well, and OD600 read-out in 15 min intervals; plates were sealed with clear Breathe-Easy membranes (Sigma-Aldrich). Plasmid DNA was prepared using Miniprep kits (Qiagen). DNA restriction digests and ligations (with T4 DNA ligase) were performed with enzymes from New England Biolabs (NEB). DNA fragment purification by gel extraction was performed using Zymoclean gel DNA recovery kits (Zymo Research). High-fidelity PCR was performed using either Phusion DNA polymerase (NEB) or PfuUltra II Fusion DNA polymerase (Agilent). Analytical colony PCR was performed with Taq DNA polymerase (Takara). Gibson assembly was performed using Gibson Assembly Master Mix (NEB). Bacterial transformation with plasmid DNA was achieved through electroporation (see below) or by chemical transformation using NEB 5-alpha *E. coli* (NEB) according to the manufacturer’s instructions. All plasmids were stored as glycerol stocks at −80°C in *E. coli* CC118 λpir (pBAMD1-2 and derivatives thereof) or *E. coli* NEB 5-alpha (all other plasmids). Synthetic DNA fragments and oligonucleotides were purchased from IDT. Sanger sequencing was performed by Laragen. Chemicals were obtained from Sigma. PBAT-starch-polylactic acid copolymer bags (Vove compostable sandwich bags) were obtained from Amazon. Commercial bottle PET was obtained from Evian water bottles. Amorphous PET film (0.25 mm thickness) and PET powder (≥ 40% crystallinity) were obtained from Goodfellow.

### Preparation and transformation of electrocompetent cells

Electrocompetent *E. coli* and *P. putida* cells were prepared by a modified standard protocol. Briefly, 250 mL of LB medium in a 1 L flask were inoculated with an *E. coli* or *P. putida* overnight culture to an OD600 of 0.05 and incubated at 250 rpm and 37°C or 30°C, respectively, until an OD600 of 0.5 was reached. Cells were cooled on ice, transferred to 250 mL centrifuge buckets, and pelleted at 4°C, 3000g, 8 min. The supernatant was removed, the cells were gently resuspended by swirling in 50 mL sterile 10% ice-cold glycerol in ddH_2_O, transferred to 50 mL tubes, and pelleted at 4°C, 3000g, 8 min. The cell pellet then underwent two additional washing steps with 50 mL 10% glycerol, before resuspending the cells in 1 mL 10% glycerol. Aliquots of 50 μL were either used directly for electroporation or frozen on dry ice in 0.7 mL tubes and stored at −80°C. For transformation, 0.5 to 1.5 μL of plasmid DNA, ligation reaction or Gibson assembly were added to a cell aliquot on ice, and electroporation was performed using a BioRad MicroPulser electroporator (Ec1 setting) and 1 mM gap width electroporation cuvettes (BioRad). SOC medium (600 μL; NEB) was added immediately following electroporation and cells were incubated at 37°C or 30°C (*E. coli* or *P. putida*, respectively) for 1 h (3-5 h in case of pBAMD genomic knock-ins) before plating on selective LB-agar plates. We found that for all *P. putida* transformations, including pBAMD-based genomic knock-ins, electroporation provided sufficient transformation rates and no conjugative plasmid transfer was necessary.

### Plasmid construction

#### 1. Cloning of the TA-to-PCA operon knock-in vector pBAMD-tphII

The mini-Tn5 transposon plasmid pBAMD1-2 was digested with EcoRI and PshAI (thereby excising the Kan^R^ selectable marker) and the vector fragment was purified by agarose gel extraction. To construct the TA-to-PCA operon, a synthetic DNA fragment containing the P_EM7_ promoter and BCD2 translational coupler^39^ (Supplementary Sequence 1) was amplified using primers OB40 and OB51. In parallel, the codon-optimized synthetic DNA fragments encoding *tphA2*_II_, *tphA3*_II_, *tphB*_II_, and *tphA1*_II_ from *Comamonas sp*. E6 (Supplementary Sequences 2-5) were amplified using primer pairs OB39 and OB54, OB55 and OB56, OB57 and OB58, and OB59 and OB53, respectively. This strategy introduced 32 bp intergenic regions between the four genes, each containing a strong ribosome binding site (ACGAGG)^89^. In addition, the outmost primers (OB51 and OB53) introduced overhangs homologous to pBAMD1-2 overlapping the EcoRI and PshAI restriction sites. Following PCR and purification of the individual gene fragments, overlap extension PCR was used to splice together the P_EM7_-BCD2, *tphA2_II_, tphA3_II_* fragments and the *thpB_II_, tphA1_II_* fragments, respectively. Next, the two resulting fragments were spliced together using overlap extension PCR to create the complete operon fragment. The fragment was cloned into EcoRI/PshAI-digested pBAMD1-2 using Gibson assembly, resulting in plasmid pBAMD-tphII. *E. coli* CC118 λpir cells were transformed with the Gibson assembly mix by electroporation and cells were plated on LB-Amp agar plates. Plasmids isolated from single clones were analyzed by Sanger sequencing to verify correct assembly of the operon and absence of undesired mutations.

#### 2. Cloning of PET hydrolase expression plasmids

##### (A) Cloning of expression vectors

To obtain a set of expression vectors harboring the rhaS-rhaR-P_rhaB_ transcriptional regulator for rhamnose-inducible protein expression and differing only in their replication system (and thus cellular copy number) we first transferred the rhaS-rhaR-P_rhaB_ element from pPS39 to pSEVA328 and pSEVA338 using PacI and SpeI restriction sites, generating vectors pSEVA328-P_rhaB_ and pSEVA338-P_rhaB_. Next, we obtained the DNA fragment encoding the Kan resistance marker from pBAMD1-2 through digest with SwaI and PshAI and inserted it into the equally digested pPS39, pSEVA338-P_rhaB_ and pSEVA328-P_rhaB_, thereby obtaining plasmids pHi-P_rhaB_, pMe-P_rhaB_ and pLo-P_rhaB_. To obtain a low copy number expression vector with the P_IbpA_ transcriptional/translational regulator, we ordered the P_IbpA_ fragment as a DNA oligo (Supplementary Sequence 6), which was then transformed into dsDNA and amplified by PCR using primers OB179 and OB192. Next, we PCR-amplified pLo-P_rhaB_ as two separate fragments, to simultaneously remove the rhaS-rhaR-P_rhaB_ element and to remove a NdeI site in the replication protein trfA (I305; primer-encoded silent codon change ATA > ATC (antisense); CATATG > CAGATG); primers used were OB191 with OB189, and OB178 with OB190. The two resulting pLo vector fragments were fused and amplified using overlap extension PCR with outer primers OB178 and OB191. Next, the vector and P_IbpA_ fragments were digested with PacI and NdeI and ligated to obtain pLo-P_IbpA_. To obtain a low copy number expression vector with the CI857/P_L_ transcriptional regulator, the CI857/P_L_ element was ordered as synthetic DNA fragment (Supplementary Sequence 7) and amplified with primers OB182 and OB183. Next, the resulting fragment was digested with PacI and AvrII and ligated into equally digested pLo-P_rhaB_ to obtain pLo-PL.

##### (B) Cloning of PET hydrolase membrane display constructs

(i) Cloning of OprF display constructs: A 100 bp BCD2 translational coupler fragment was amplified from a synthetic DNA fragment (Supplementary Sequence 8) with primers OB85 and OB91. Next, the OprF fragment including a C-terminal G_4_SG_2_S(G_4_S)_3_ linker^50^ was amplified from a synthetic DNA fragment (Supplementary Sequence 9) with primers OB92 and OB93. The BCD2 and OprF fragments were fused and further amplified by overlap extension PCR with outer primers OB85 and OB93. The resulting fragment was digested with AvrII and HindIII and ligated into equally digested pHi-P_rhaB_, pMe-P_rhaB_ and pLo-P_rhaB_, generating plasmids pHi-P_rhaB_-OprF, pMe-P_rhaB_-OprF and pLo-P_rhaB_-OprF. Next, synthetic DNA fragments encoding LCC (Supplementary Sequence 8, primers OB85 and OB86), HiC (Supplementary Sequence 10, primers OB94 and OB95) and IsP (Supplementary Sequence 11, primers OB132 and OB133) were amplified by PCR, digested with PstI and NcoI, and ligated into equally digested pHi-P_rhaB_-OprF, pMe-P_rhaB_-OprF and pLo-P_rhaB_-OprF (note, IsP was only cloned into pLo-P_rhaB_-OprF). This generated plasmids pHi-P_rhaB_-OprF-LCC, pHi-P_rhaB_-OprF-HiC, pMe-P_rhaB_-OprF-LCC, pMe-P_rhaB_-OprF-HiC, pLo-P_rhaB_-OprF-LCC, pLo-P_rhaB_-OprF-HiC, and pLo-P _rhaB_-OprF-IsP. (ii) Cloning of EhaA display constructs: The codon-optimized synthetic EhaA DNA fragment (Supplementary Sequence 12) including an N-terminal G4SG2S(G4S)3 linker was PCR-amplified with primers OB87 and OB88. A synthetic codon-optimized DNA fragment (Supplementary Sequence 8) encoding the N-terminal BCD2 translational coupler followed by residues 1-37 of *P. putida* OprF, a 6xHis tag, PstI restriction site, residues 35-293 of LCC and a NcoI restriction site was amplified by PCR using primers OB85 and OB86. Next, the EhaA and LCC fragments were fused and further amplified by overlap extension PCR with outer primers OB85 and OB88 (5% DMSO were added to the PCR mix to ensure efficient amplification across GC-rich regions). The resulting fragment was digested with AvrII and HindIII and ligated into equally digested pHi-P_rhaB_, pMe-P_rhaB_ and pLo-P_rhaB_, generating plasmids pHi-P_rhaB_-EhaA-LCC, pMe-P_rhaB_-EhaA-LCC and pLo-P_rhaB_-EhaA-LCC. To generate the corresponding EhaA-HiC and EhaA-IsP constructs, the LCC fragment was excised from these plasmids with PstI and NcoI and replaced with equally digested HiC and IsP ORF fragments (see OprF cloning above) (note, IsP was only cloned into pLo-P_rhaB_-EhaA). (iii) Cloning of InaV-NC display constructs: A 100 bp BCD2 translational coupler fragment was amplified from a synthetic DNA fragment (Supplementary Sequence 8) with primers OB85 and OB91. Next, the InaV-NC fragment including a C-terminal G_4_SG_2_S(G_4_S)_3_ linker was amplified from a synthetic DNA fragment (Supplementary Sequence 13) with primers OB89 and OB90. The BCD2 and InaV-NC fragments were fused and amplified by overlap extension PCR with outer primers OB85 and OB89. The resulting fragment was digested with AvrII and HindIII and ligated into equally digested pHi-P_rhaB_, pMe-P_rhaB_ and pLo-P_rhaB_, generating plasmids pHi-P_rhaB_-InaV, pMe-P_rhaB_-InaV and pLo-P_rhaB_-InaV. Next, synthetic DNA fragments encoding LCC (Supplementary Sequence 8, primers OB85 and OB86) and HiC (Supplementary Sequence 11, primers OB94 and OB95) were amplified by PCR, digested with PstI and NcoI, and ligated into equally digested pHi-P_rhaB_-InaV, pMe-P_rhaB_-InaV and pLo-P_rhaB_-InaV. This generated plasmids pHi-P_rhaB_-InaV-LCC, pHi-P_rhaB_-InaV-HiC, pMe-P_rhaB_-InaV-LCC, pMe-P_rhaB_-InaV-HiC, pLo-P_rhaB_-InaV-LCC, and pLo-P_rhaB_-InaV-HiC.

##### (C) Cloning of PET hydrolase secretion constructs

First, we excised the EhaA coding sequence from pLo-P_rhaB_-EhaA-LCC with NcoI and HindIII. Next, we used PCR to generate a short 85 bp linker fragment from overlapping oligonucleotides OB121 and OB122, spanning across the NcoI and HindIII sites and introducing a stop codon after the LCC coding sequence and NcoI site. Gibson assembly was used to repair the pLo-P_rhaB_-EhaA-LCC NcoI/HindIII-digested vector fragment with this linker fragment, resulting in plasmid pLo-P_rhaB_-SP(OprF)-LCC. Next, we used site-directed mutagenesis with primer pair OB134 and OB135 to introduce a silent mutation creating a unique BbsI site at the C-terminal end of the BCD2 translational coupler (V16, GTT to GTC). This generated plasmid pLo-P_rhaB_-SPswitch-LCC, allowing facile exchange of the N-terminal SP preceding the LCC coding region using the BbsI/PstI restriction sites. We next generated analogous plasmids pLo-P_rhaB_-SPswitch-HiC and pLo-P_rhaB_-SPswitch-IsP by digesting pLo-P_rhaB_-SPswitch-LCC with PstI and NcoI to excise the LCC coding sequence and replacing it with equally digested HiC and IsP coding fragments obtained from pLo-P_rhaB_-EhaA-HiC and pLo-P_rhaB_-EhaA-Isp. To clone a library of different signal peptides (SPs) into the LCC, HiC, and IsP SPswitch vectors, we ordered a 30-member SP library as oligonucleotides (oPool) from IDT (Supplementary Table 3). The library was transformed into dsDNA and further amplified by PCR with outer primers OB136 and OB137. Next, the SP library and plasmids pLo-P_rhaB_-SPswitch-LCC, pLo-P_rhaB_-SPswitch-HiC, and pLo-P_rhaB_-SPswitch-IsP were digested with BbsI and PstI and the SP library was ligated into the vector fragments. *E. coli* NEB5α cells were transformed with the ligation reactions and serial dilutions plated on LB-Km plates and incubated o/n. The next day, colonies from a dense plate (> 250 colonies) for each construct were washed off with 5 mL LB and directly processed by Miniprep; the isolated plasmid library was stored at −20C and used to transform *P. putida* for subsequent SP library screening. In parallel, plasmid DNA isolated from five single colonies per construct was sent for sequencing to verify the cloning procedure and SP library diversity. Individual SP-PET hydrolase plasmids as listed in Supplementary Table 5 were obtained after library screening.

##### (D) Cloning of PET hydrolase cytoplasmic expression plasmids

To obtain plasmids for cytoplasmic expression of PET hydrolases (without membrane anchor or signal peptide), the LCC, HiC, and IsP ORFs including a 6xHis tag at the C-terminus were PCR-amplified from pLo-P_rhaB_-OprF-LCC, pLo-P_rhaB_-OprF-HiC, and pLo-P_rhaB_-OprF-IsP with primer pairs OB139 and OB211, OB139 and OB210, and OB139 and OB209, respectively. The fwd primers introduced a BbsI restriction site matching the site in the pLo-P_rhaB_-SPswitch vectors, while the reverse primer introduced a HindIII site. The PCR fragments were digested with BbsI and HindIII and ligated into equally digested pLo-P_rhaB_-SPswitch-LCC to generate pLo-P_rhaB_-LCC, pLo-P_rhaB_-HiC, and pLo-P_rhaB_-IsP.

##### (E) Cloning of PET hydrolases into heat-shock promoter plasmids

To obtain the EhaA-LCC display construct under control of P_L_, the pLo-P_rhaB_-EhaA-LCC plasmid was digested with PacI and AvrII to excise the rhaR-rhaS-P_rhaB_ element, which was then replaced with the equally digested CI857/P_L_ element (obtained from PLo-P_L_) to generate pLo-P_L_-EhaA-LCC. To obtain the secreted bglx-HiC construct under control of P_L_, pLo-P_L_ was digested with AvrII and HindIII. The bglx-HiC fragment (including the BCD2 translational coupler) was obtained from pLo-P_rhaB_-bglx-HiC by AvrII/HindIII digest and ligated into the equally digested pLo-PL fragment to obtain pLo-P_L_-bglx-HiC. To obtain the EhaA-LCC display construct under control of P_IbpA_, the EhaA-LCC fragment was obtained by PCR with primers OB201 and OB139 using pLo-P_rhaB_-EhaA-LCC as template. pLo-P_IbpA_ was digested with NdeI and HindIII, and the EhaA-LCC fragment was ligated into the vector using Gibson assembly, providing pLo-P_IbpA_-EhaA-LCC. To obtain the secreted bglx-HiC and lasB-IsP constructs under control of P_IbpA_, both ORFs were amplified by PCR using primer pairs OB197 and OB139 and pLo-P_rhaB_-bglx-HiC as template, and primer pairs OB195 and OB139 and pLo-P_rhaB_-lasb-IsP as template, respectively. The PCR fragments were digested with NdeI and HindIII and ligated into equally digested pLo-P_IbpA_, yielding pLo-P_IbpA_-bglx-HiC and pLo-P_IbpA_-lasb-IsP. To obtain pLo-P_IbpA_-lasb-IsP^Dura^, the P_IbpA_-IsP^Dura^ fragment was amplified from a synthetic DNA fragment (Supplementary Sequence 14) using primer pair OB220 and OB221, digested with PacI and HindIII, and cloned into equally digested pLo-P_IbpA_.

#### 3. Cloning of P_IbpA_-PET hydrolase genomic knock-in vectors

To obtain plasmids for genomic knock-in of P_IbpA_-PET hydrolase expression systems we used pBAMD1-2. For secreted constructs, we first PCR-amplified the P_IbpA_-bglx-HiC and P_IbpA_-lasb-IsP^Dura^ cassettes from their pLo-based expression plasmids (see above) using primer pairs OB143 and OB205, and OFB220 and OFB205, respectively. The PCR products and pBAMD1-2 were then digested with SacI and HindIII and ligated to yield pBAMD-P_IbpA_-bglx-HiC and pBAMD-P_IbpA_-lasb-IsP^Dura^. For P_IbpA_-EhaA-LCC we used a Gibson assembly strategy due to the presence of a SacI site in LCC. The P_IbpA_-EhaA-LCC cassette was PCR_amplified from the corresponding pLO plasmid (see above) using primer pair OB119 and OB214. The resulting fragment was cloned into SacI/HindIII digested pBAMD1-2 using Gibson assembly to yield pBAMD-P_IbpA_-EhaA-LCC. In contrast to pBAMD-tphII, all P_IbpA_-PET hydrolase knock-in vectors retained the Kan resistance marker to allow selection of transformants. We also cloned the corresponding pBAMD-P_L_-PET hydrolase constructs, however, they consistently showed very low knock-in efficiencies and are not discussed in the main text; details are available on request.

#### 4. Cloning of mhpT expression vectors

mhpT^wt^ and mhpT^L374V^ ORFs were amplified from genomic DNA of *P. putida* TA and *P. putida* TA7 by PCR using primer pair OB109 and OB110. The fwd primer OB109 introduced an AvrII restriction site and a strong ribosome binding site (ACGAGG) in front of the mhpT ATG, while the rev primer OB110 introduced a HindIII restriction site after the mhpT stop codon. The fragments were digested with AvrII and HindII and ligated into equally digested pMe-P_rhaB_, generating plasmids pMe-P_rhaB_-mhpT^wt^ and pMe-P_rhaB_-mhpT^L374V^.

### Metabolic engineering and adaptive laboratory evolution of P. putida strains

#### 1. Engineering TA-to-PCA catabolism

To knock-in the TA-to-PCA cassette into P. *putida*, 50 μL of electrocompetent *P. putida* KT2440 cells were transformed with pBAMD-tphII (75 ng) and incubated in 600 μL SOC medium for 3 h at 30°C and 250 rpm. The cells were pelleted (3000g, 3 min), resuspended in 1 mL M9 medium, and added to 25 mL M9 medium, pH 5.5, 0.4 % w/v TA, in a 125 mL flask. The culture was incubated at 30°C, 250 rpm, until after ca. 36 h noticeable bacterial growth was observed. The culture was subsequently passaged daily for 10 days (dilution to OD600 = 0.01 in 15 mL fresh M9-TA medium), at which point the culture was streaked on LB-agar plates to obtain single colonies. Five clones were processed and analyzed by PCR as described by Garcia-Martinez *et al*.^90^ to determine the genomic integration site; primers used were OB96 and OB105 (5’ end, outer primers), OB97 and OB104 (5’ end, inner primers), OB96 and OB102 (3’ end, outer primers) and OB97 and OB103 (3’ end, inner primers). The resulting DNA fragments spanning over the 5’ and 3’ integration sites were sequenced using primers OB104 and OB103, respectively; all five clones showed the same integration site, indicating that the knock-in culture was dominated by a single clone likely originating from the same integration event. Growth of one selected clone on TA as sole carbon source was verified and the variant was designated as strain *P. putida* TA.

#### 2. ALE for growth of P. putida TA at pH 7

To adapt *P. putida* strain TA with the genomically integrated TA-to-PCA cassette for better growth at higher pH, 25 mL of M9 pH 7.0, 0.4% w/v TA were inoculated with 0.5 mL of the *P. putida* TA-to-PCA knock-in culture after 10 days of passaging (see above) and cultured at 30°C, 250 rpm. After 7 days (6 total passages), significantly better bacterial growth was observed and the culture was streaked out on LB-agar plates to obtain single colonies. Three clones were tested for growth in M9, 0.4% TA at different pH in comparison to *P. putida* TA; the best-performing clone was designated strain *P. putida* TA7 and subjected to whole-genome sequencing together with *P. putida* TA to identify potential genetic changes.

#### 3. ALE for co-utilization of ethylene glycol by P. putida TA7

To adapt *P. putida* TA7 for co-utilization of ethylene glycol (EG), two duplicate cultures of 20 mL M9 pH 8.0, 5 mM TA, 25 mM EG in 125 mL flasks were inoculated with strain TA7 and cultured at 30°C and 250 rpm. Once bacterial growth was observed (typically after 24-48h) cultures were passaged to a fresh flask (to OD600 0.01). After 11 days (9 passages), significantly improved bacterial growth was observed and cultures were streaked out on LB-agar plates to obtain single colonies. Four single clones from each culture (8 total) were tested in comparison to *P. putida* TA7 for growth in M9 minimal medium with varying concentrations of TA and EG; the best-performing strain was designated as *P. putida* TA7-EG and analyzed by whole-genome sequencing.

#### 4. ALE for co-utilization of 1,4-butanediol by P. putida TA7

To adapt *P. putida* TA7 for co-utilization of 1,4-butanediol (BD), two cultures of 20 mL M9 pH 8.0 with either 5 mM TA and 25 mM BD, or 5 mM TA, 25 mM BD, and 25 mM AA, in 125 mL flasks were inoculated with strain TA7 and cultured at 30°C and 250 rpm. Once bacterial growth was observed (typically after 24-48h) cultures were passaged to fresh flasks (to OD600 0.01). After 11 days (7 passages), significantly improved bacterial growth was observed and cultures were streaked out on LB-agar plates to obtain single colonies. Four single clones from each culture (8 total) were tested in comparison to *P. putida* TA7 for growth in M9 minimal medium with varying concentrations of TA, BD, and AA; the best-performing strain was designated as *P. putida* TA7-BD and analyzed by whole-genome sequencing.

#### 5. Genomic Integration of P_IbpA_-PET hydrolase cassettes

The pBAMD-derived P_IbpA_-PET hydrolase knock-in plasmids (pBAMD-P_IbpA_-bglx-HiC, pBAMD-P_IbpA_-lasb-IsP^Dura^, pBAMD-P_IbpA_-EhaA-LCC) were delivered to *P*. *putida* by electroporation and cells were recovered in LB medium at 25°C, 250 rpm, o/n. Next, serial dilutions of the cell suspension were plated on LB-Kan agar plates and incubated at 25°C for ca. 36 h. For each knock-in, sixteen single colonies were transferred using sterile toothpicks into deep 96-well plates supplemented with 250 μL LB-Kan per well and incubated at 25°C, 250 rpm, o/n; control wells with the parent strain were also included. Next, 50 μL of the cell suspension were used to inoculate a fresh deep 96-well plate supplemented with 500 μL LB-Kan per well and the plate was incubated at 25°C, 250 rpm, for 3 h; the pre-culture plate was sealed and stored at 4°C. The expression plate was then subjected to a heat shock (37°C, 250 rpm) for 2 h, followed by an outgrowth phase at 30°C, 250 rpm, o/n. We then assessed esterase activity in the cleared supernatant and cell fraction of each well using the pNBP assay. The best-performing strains were re-tested in shake flask cultures, followed by preparation of glycerol stocks and whole-genome sequencing.

### pNPB assay for esterase activity measurements

All pNPB esterase assays were carried out in Tris buffer (100 mM Tris-HCl pH 8.0, 50 mM NaCl). A calibration curve was prepared by measuring A405 of serial dilutions of 4-nitrophenol (4NP; Sigma-Aldrich) in assay buffer (95 μLTris buffer, 5 μL DMSO, in clear flat-bottom 96-well plates; Supplementary Figure 10). For measuring PET hydrolase esterase activity, 1 mL PET hydrolase expression cultures were pelleted in 1.5 mL tubes (500 g, 2 min). 5 μL (in duplicates) of the cleared supernatant (or further dilutions thereof if esterase activity was high) were directly added to clear flat-bottom 96-well plates. The remaining supernatant was discarded and the cell pellet carefully resuspended in 1 mL Tris buffer, followed by a second centrifugation. The supernatant was discarded and the pellet carefully resuspended in 1 mL Tris buffer, and 5 μL (in duplicates) added to clear flat-bottom 96-well plates. We then prepared a mastermix of Tris buffer with pNPB substrate (per well: 90 μL Tris buffer with 5 μL pNPB stock solution (10 mM in DMSO)); multiples of this mix were prepared according to the number of wells to be analyzed (and including a pipetting reserve), and 95 uL were added to the 96-well sample plate with a multichannel pipette to start the reaction (final concentrations: 0.5 mM pNPB and 5% DMSO). Each plate contained blank wells (5 μL clean culture medium or Tris buffer instead of culture supernatant or cell suspension). The plate was immediately inserted into the plate reader, and A405 of each well was measured in 31 s intervals (the shortest possible measurement interval of the plate reader). The resulting data were plotted as A405 over time (in s) and the data points showing a linear increase were included to derive the slope by linear regression (plotting and analysis were performed in GraphPad Prism 9); the slope derived from blank controls was subtracted. The resulting slope (A405/s) was then used together with the slope of the calibration curve (A405/μMol 4NP) to derive μMol 4NP/min = ((A405/s)/(A405/[4NP]))*60. The resulting value was multiplied by 200 (as we typically used 5 μL sample per reaction) to obtain μMol 4NP/min*mL. As the amount of enzyme required to release1 μMol 4NP/min is defined as one Unit (U), this term represents U/mL.

### PET hydrolase expression cultures

To test PET hydrolase expression, single colonies of *P. putida* strains (either transformed with PET hydrolase expression plasmids or strains with genomic integration of PET hydrolase expression cassettes) were inoculated into 3 mL medium (LB or M9 minimal medium, the latter supplemented with TA, EG or BD) and cultured at 250 rpm, o/n, 30°C (all P_rhaB_ constructs) or 25°C (all P_L_ and P_IbpA_ constructs). The next day, expression cultures in either 125 ml shake flasks or 14 mL tubes and containing either LB or M9 medium were inoculated with the pre-cultures to OD600 0.1 and incubated for an additional 2-3 h under the same conditions detailed above, typically to an OD600 of 0.2 to 0.5 (cultures in M9 minimal medium supplemented with TA, EG, or BD reached lower final OD600 than cultures in LB medium and were thus also induced at lower OD600). Induction of expression was performed through addition of 2 mM rhamnose (all P_rhaB_ constructs) or transferring the cultures to 30°C or 37°C (all P_IbpA_ constructs); if P_IbpA_ cultures were subjected to a 37°C heat shock, they were typically transferred back to 30°C at defined time points (see figures for details). At selected time points post-induction, culture samples were taken to analyze OD600 and pNPB esterase activity in the culture supernatant and the cell fraction.

### Signal peptide library screen

*P. putida* TA7-EG was transformed with the LCC, HiC, and IsP SP plasmid libraries and plated on LB-Kan plates, 30°C, o/n; cells transformed with pLo-P_rhaB_ (e.v.) were included as control. For each library, pre-cultures in deep 96-well plate with 500 μL LB-Kan per well were prepared and inoculated with single colonies using toothpicks; 4 wells with e.v. control colonies were included per plate. The plates were sealed with Breathe-Easy membranes and incubated at 30°C, 250 rpm, o/n. Next, expression culture deep 96-well plates with 500 μL LB-Kan were prepared and inoculated with 25 μL of the pre-cultures; the pre-culture plates were sealed and stored at 4°C. The expression plates were incubated at 30°C, 250 rpm, for 2 h and induced by addition of 2 mM rhamnose (final concentration), followed by continued incubation at 30°C, 250 rpm. 3 h and 20 h post-induction, 100 μL cell culture per well were removed and analyzed for cleared supernatant esterase activity (using an end point assay, not the kinetic measurement) and OD600. Esterase activity was plotted against OD600 and those clones showing the best combination of the two measurements were manually selected for further analysis. For selected clones, the corresponding wells from the pre-culture plates were streaked out on LB-Kan agar plates to obtain single colonies, and for each clone a single colony was used for plasmid DNA miniprep and sequencing to identify the SP. Those SPs showing up multiple times across the set of sequenced clones were tested in shake flask expression cultures and plasmid glycerol stocks in *E. coli* NEB 5-alpha were prepared.

### Plastic degradation and synthetic PETtrophy assay setups

Plastic degradation assays were typically carried out in 125 mL flasks (closed with aluminum foil) or 6-well culture plates (closed with plastic lids). To transfer sterile plastic film samples into these vessels, the plastic film was sequentially washed in beakers containing 1% SDS in ddH_2_O, ddH_2_O, and 70% EtOH. From the EtOH bath, the samples were directly placed with forceps into sterile flasks or plates, the vessel was closed again, and allowed to dry at 30°C o/n to ensure evaporation of remaining EtOH. For powdery or ground plastic samples, the washing steps were carried out in sterile 1.7 mL tubes with centrifugation to pellet particles in between washing steps, and the sample was finally taken up in 1 mL 70% EtOH (using 1 mL pipette tips with cut-off tips, when necessary) and transferred to the culture vessel. Single colonies of *P. putida* strains with genomically integrated P_IbpA_-PET hydrolase expression cassettes (and the matching control strains) were inoculated into 3 mL LB medium or M9 minimal medium supplemented with plastic monomers as carbon sources; generally, the pre-culture medium was matched to the medium used for later plastic degradation assays, to ensure full activity of the necessary metabolic pathways. Pre-cultures were incubated at 25°C, 250 rpm, o/n. The next day, fresh 3 mL cultures were inoculated with the pre-cultures to OD600 0.1 and grown for 25°C, 250 rpm, 2 h, followed by a heat shock at 37°C for 2 h and 250 rpm. Next, the degradation cultures were started by addition of medium to the plastic-containing vessels (15 mL per 125 mL flask, or 2 mL per well for 6-well plates) and cultures were inoculated with 1 or 0.1 mL of the post-heat shock cultures, respectively. For synthetic PETtrophy cultures (in M9 minimal medium without carbon source), this procedure introduced a small amount of carbon source-containing medium, and we experimented with washing the cells in M9 minimal medium prior to culture inoculation. However, especially for secreted constructs, this also implies loss of any already expressed PET hydrolases; in any case we did not observe consistent synthetic PETtrophy for neither of the two inoculation approaches. Cultures were maintained at 30°C, 75 rpm (125 mL flasks) or static (6-well plates), with a 2 h heat shock at 37°C per day. The culture medium was changed every 1 to 7 days by removing 80 % of the remaining medium and resupplying it to a volume of 15 or 2 mL, respectively. Cultures were inspected daily visually, by stereo and optical microscopy, and by measuring OD600. Following culture termination, plastic samples were collected, washed using the same washing procedure as before, and weighed.

### Whole-genome sequencing of P. putida strains

Genomic DNA was extracted from one OD600 unit of *P. putida* cells in exponential growth phase using the Qiagen DNeasy kit according to the manufacturer’s instructions. Quality of genomic DNA was checked on a 0.8 % agarose gel and samples were processed for sequencing using an Illumina DNA Prep kit (Illumina #20018704) according to the manufacturer’s instructions. Briefly, genomic DNA was diluted to approximately 10 ng/μL in H_2_Oup, 30 μL (300 ng) were subjected to the Nextera tagmentation reaction, and tagmented DNA was cleaned up using magnetic beads. Next, we ran a 5-cycle PCR to add unique dual indexes (IDT for Illumina DNA/RNA UD Indexes, Integrated DNA Technologies). The resulting PCR products were cleaned up and size-selected using magnetic beads. Equal volumes were then pooled, followed by adjustment of the concentration to 4 nM using resuspension buffer (RSB), assuming a fragment length of 600 bp. Subsequently, 5 μL library were denatured by adding 5 μL 0.2 M NaOH and incubating for 5 min at room temperature. The denatured library (10 μl) was then combined with 1323 μL hybridization buffer (HT1) to yield a 15 pM library. 600 μL of this diluted library were loaded on a MiSeq v2 reagent cartridge for 300 cycles and paired-end sequencing (2x 150-bp reads) was performed on an Illumina MiSeq instrument using a v3 flow cell.

The reference genome sequence for Pseudomonas putida KT2440 and annotations were downloaded from the Pseudomonas Genome DB (http://www.pseudomonas.com/strain/show/110, accessed on September 19, 2019).^91^ Chromosome names were adjusted to match across files and sequences of the knock-in cassettes were added to the sequence file. Paired-end sequencing reads were then aligned against the reference genome using Snippy (version 4.6.0, https://github.com/tseemann/snippy). Snippy output files (snps.csv) were further processed with custom R code in RStudio (R version 4.0.3, RStudio version 1.3). To determine cassette integration sites and look for potential mutations in the knock-in cassettes, the aligned reads were visualized with the Integrative Genomics Viewer (IGV).^92^

### SDS-PAGE of soluble and membrane fractions of PET hydrolase expression cultures

Isolation of the *P. putida* outer membrane fraction was adapted from established protocols.^93,94^ 20 OD600 units of PET hydrolase membrane display expression cultures (harvested 3 h post-induction for P_rhaB_ constructs, or 20 h post-induction for P_IbpA_ constructs) were pelleted (3000g, 3 min) in 15 mL tubes and stored at −20°C. The pellets were thawed on ice and resuspended in 3 mL PBS (Gibco) supplemented with 0.5 mg/mL lysozyme and 2 mM EDTA. The cell suspension was sonicated on ice using a Fisher Scientific Sonic Dismembrator 550 with a microtip (settings: level 3, 1 min total, 5 s on/off cycles). 2 mL of sample were transferred to a 2 mL tube and centrifuged (21000 g, 2 min, 4°C) to remove intact cells and large debris. 1.8 mL of supernatant were transferred to a fresh 2 mL tube and centrifuged (21000 g, 40 min, 4°C). The supernatant was carefully removed (and set aside as cytosolic fraction) and the pellet was resuspended in 1 mL PBS, 2 % Triton X-100. The sample was incubated at 37°C for 30 min and centrifuged (21000 g, 30 min, 4°C). The supernatant was carefully removed and the pellet resuspended in 45 μL PBS and 20 μL 4x Laemmli sample buffer (Biorad) supplemented with 2-mercaptoethanol. Likewise, 15 μL of cytosolic fraction samples were mixed with 5 μL 4x Laemmli sample buffer. The samples were heated at 95°C for 5 min and 5 to 15 μL were loaded on mini-Protean TGX precast 12% gels (Biorad). The gels were run using mini-Protean Tetra cells and Tris/Glycine/SDS buffer (Biorad) according to the manufacturer’s instructions. Gels were stained with Coomassie Brilliant Blue R-250 (1 g/L; BioRad) in 40% methanol, 10% acetic acid, in ddH_2_O; the same solution without Coomassie was used for gel destaining.

### SDS-PAGE of secreted PET hydrolase expression constructs

50 mL of PET hydrolase secretion expression cultures (harvested 3 h post-induction for P_rhaB_ constructs, or 20 h post-induction for P_IbpA_ constructs) were collected in 50 mL tubes and centrifuged (4000g, 10 min, 4°C). The supernatant was carefully transferred to a fresh tube without disturbing the pellet and concentrated (3000 g, 4°C, 15 min centrifugation steps with intermittent carful mixing) in Amicon Ultra-15 spin concentrators (Millipore) with 10 kDa molecular weight cut-off to a final volume of ca. 500 μL. 15 μL of this concentrate were mixed with 5 μL of 4x Laemmli sample buffer and analyzed by SDS-PAGE as described above.

### LC-MS analysis of SDS-PAGE bands

Protein bands of interest were excised from the gel and further processed as described by Shevchenko and colleagues with minor modifications.^95^ Briefly, gel pieces were destained, followed by in-gel protein reduction, alkylation, and digestion using MS-grade trypsin (Pierce / Thermo Scientific, #90057). After extraction, peptides were desalted using C18 tips (Pierce / Thermo Scientific #87784), dried under vacuum and reconstituted in 5% formic acid (v/v). For liquid chromatography-coupled tandem mass spectrometric (LC-MS/MS) analysis, desalted peptides were fractionated online by nano-flow reversed phase liquid chromatography coupled to an Orbitrap Fusion Lumos Tribrid mass spectrometer (Thermo Fisher Scientific). Data was acquired in Data-Dependent Acquisition mode.

Peptide identification and quantification was performed with MaxQuant version 1.6.17.0.^96^ The search was performed against a *P. putida* KT2440 protein database downloaded from UniProt (https://www.uniprot.org/proteomes/UP000000556, accessed on January 20, 2022). To this database, which contained 5527 proteins, the sequences for the three PET hydrolases and the membrane anchor EhaA were added. Only fully LysC- and trypsin-digested peptides with up to one missed cleavage were allowed. Carbamidomethylation of cysteines was defined as a fixed modification. Peptide-spectrum-match (PSM) and protein-level false discovery rates (FDRs) were set to 0.01. The MaxQuant output (evidence.txt) was then filtered for peptides belonging to the PET hydrolases and with an intensity of >1e9.

### SEM of plastic samples

SEM images were taken by the Molecular Instrumentation Center at UCLA’s Department of Chemistry and Biochemistry. Briefly, plastic samples were sequentially washed with 70 % EtOH, 1 % SDS in ddH_2_O, and ddH_2_O, and air dried. The samples were coated with an 8 nm gold layer, and images were acquired using a JEOL JSM-6700F FE-SEM using the LEI detector.

## Author Information

### Author contributions

OFB designed the research; OFB and OTS performed experiments; OFB and OTS analyzed data; LK supervised the project; OFB wrote the manuscript; all authors critically revised the manuscript and have read and approved the final version.

## Acknowledgements

We thank all members of the Kruglyak lab for helpful comments and discussions. We thank Dr. Yasaman Jami at the UCLA Department of Biological Chemistry for help with LC-MS measurements. We also thank Dr. Ignacio Martini and Dr. Gregory Khitrov at the UCLA Molecular Instrumentation Center for help with SEM and HPLC measurements, respectively.

