## Supplementary Information for "Towards Synthetic PETtrophy: Engineering *Pseudomonas putida* for concurrent polyethylene terephthalate (PET) monomer metabolism and PET hydrolase expression"

**Contents:**


| | | |
|---|---|---|
| I | Supplementary Figures | p. 2 |
| II | Supplementary Tables | p. 14 |
| III | Supplementary DNA Sequences | p. 27 |
| Iv | Supplementary References | p. 36 |

### I Supplementary Figures

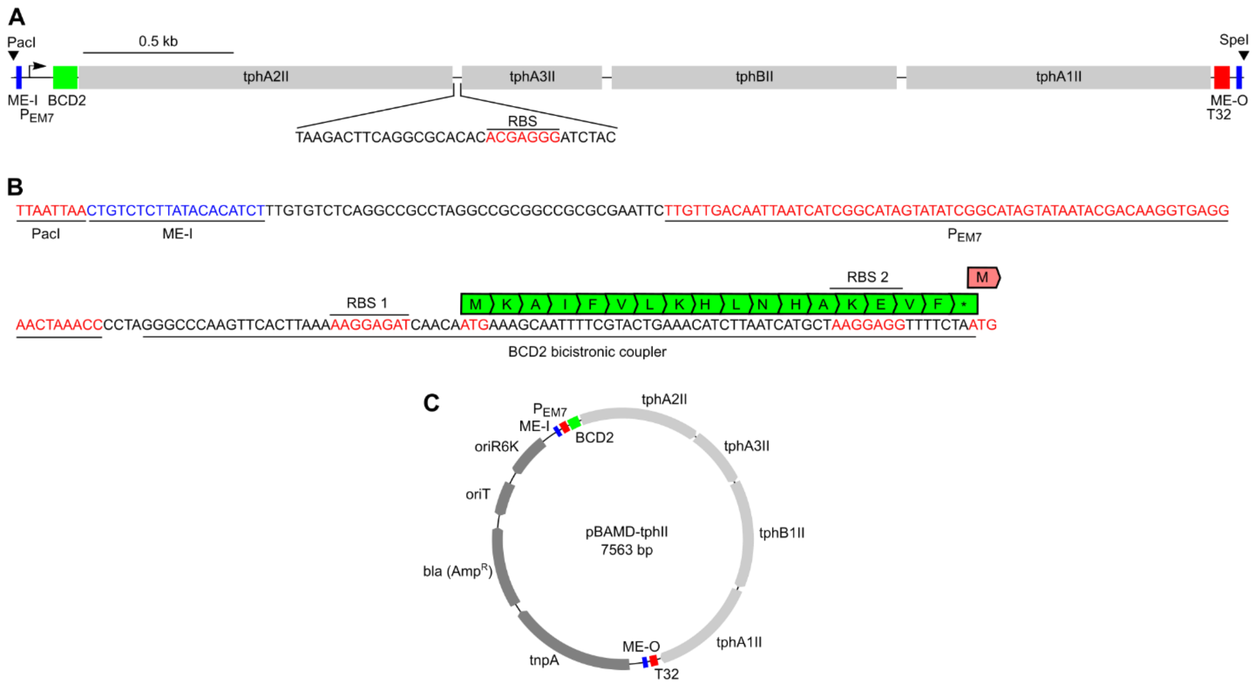


**Supplementary Figure 1: Design of pBAMD-tphII.** (A) Layout of the $tph_{II}$ synthetic operon. The knock-in cassette is flanked by unique PacI and SpeI restriction sites, followed by the ME-I and ME-O 5' and 3' end transposase recognition sites, respectively. The operon is under transcriptional control of the constitutive $P_{EM7}$ promoter and endowed with a BCD2 bicistronic translational coupler. The four ORFs are separated by 32 bp intergenic regions, each containing an additional ribosome binding site. The operon is followed by a T32 transcriptional terminator. (B) Sequence view of the knock-in cassette 5' end from the PacI site to the ATG of *tphA2$_{II}$*, showing ME-I (blue), $P_{EM7}$ (red), and the BCD2 translational coupler including the amino acid sequence of its short ORF. (C) Plasmid map of pBAMD-tphII; all functional elements outside of the $tph_{II}$ operon remained unchanged and as described for the original pBAMD1-2 plasmid.[1]

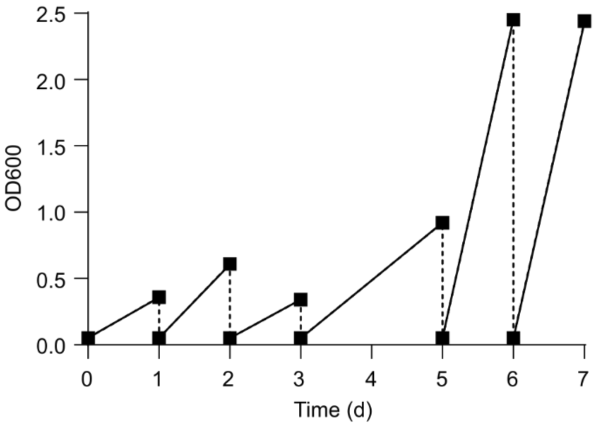


**Supplementary Figure 2: ALE with *P. putida* TA for growth at elevated pH.** *P. putida* TA was cultured in M9 minimal medium, 0.4% (w/v) TA, pH 6.5, for 7 days (6 passages). After 7 days we observed improved bacterial growth and proceeded to the isolation and analysis of single clones, which yielded strain *P. putida* TA7.

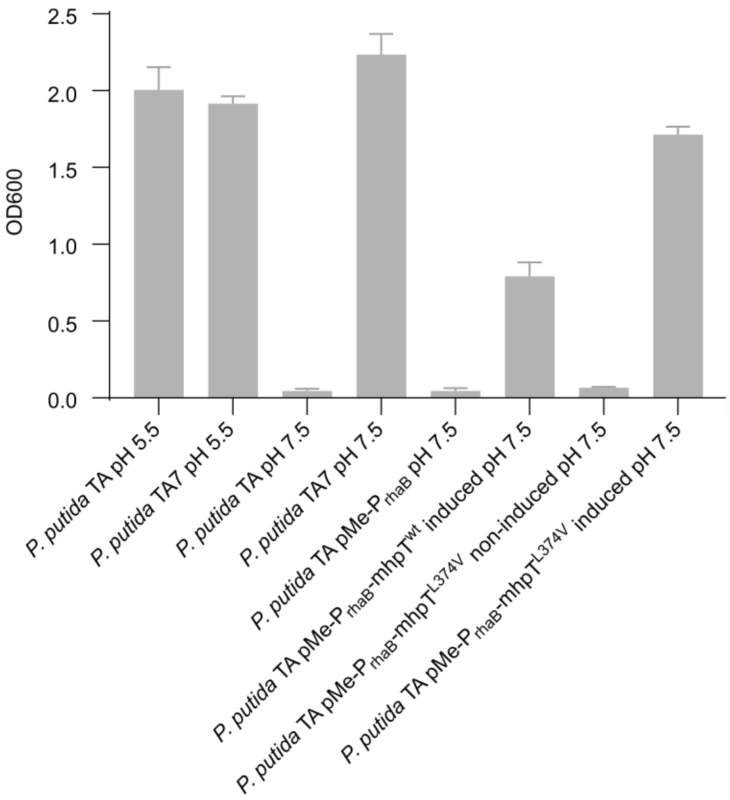


**Supplementary Figure 3: Rescue of *P. putida* TA growth at elevated pH by plasmid-based mhpT overexpression.** While both *P. putida* TA and TA7 grow well in M9 minimal medium with 0.4% TA at pH 5.5, only *P. putida* TA7 grows at pH 7.5. However, growth of *P. putida* TA at pH 7.5 can be rescued, to some extent, by plasmid-based overexpression of $mhpT^{wt}$ (from a medium copy number plasmid and under control of the $P_{rhaB}$ promoter), indicating promiscuous function of mhpT as TA transporter. Better rescue of *P. putida* TA at pH 7.5, approaching the growth level of *P. putida* TA7, can be achieved by overexpression of mhpT$^{L374V}$, while no rescue is seen for e.v. or non-induced controls. Data shown are mean and S.D., n=2.

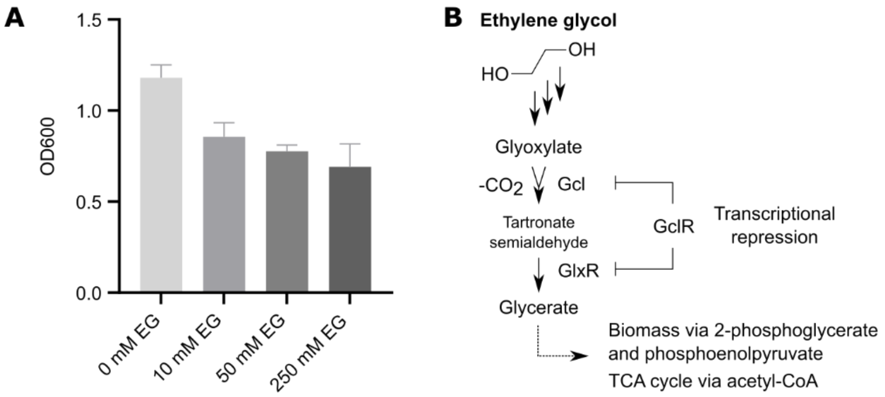


**Supplementary Figure 4: Negative effect of high EG concentrations on *P. putida* growth, and the *P. putida* EG metabolic pathway.** (A) Growth (OD600) of *P. putida* TA7 overnight cultures in M9 minimal medium, 10 mM TA, pH 7.5, with varying amounts of EG; shown are mean and S.D from n=2 cultures. (B) Pathway of EG metabolization in *P. putida*, showing the crucial function of the GclR transcriptional repressor, blocking expression of the enzymes Gcl and GlxR required for productive EG metabolism.[2–4]

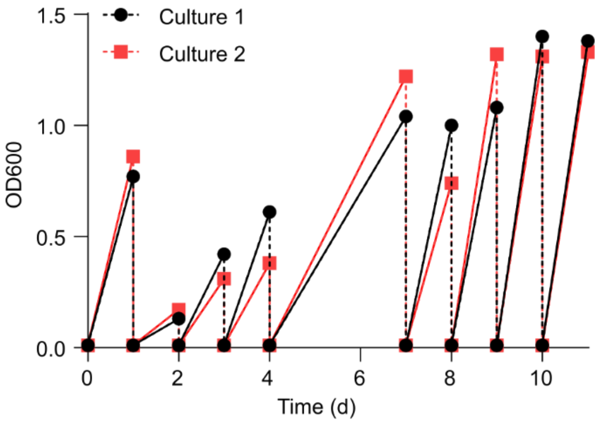


**Supplementary Figure 5: ALE with *P. putida* TA7 for EG utilization**. We performed ALE with *P. putida* TA7 in duplicate cultures using M9 minimal medium containing 25 mM EG and 5 mM TA, pH 7.5. Single clones were isolated on day 11 followed by further characterization and whole-genome sequencing, yielding strain *P. putida* TA7-EG. This strain carries a likely GclR loss-of-function mutation (c.388G>T, E130*).

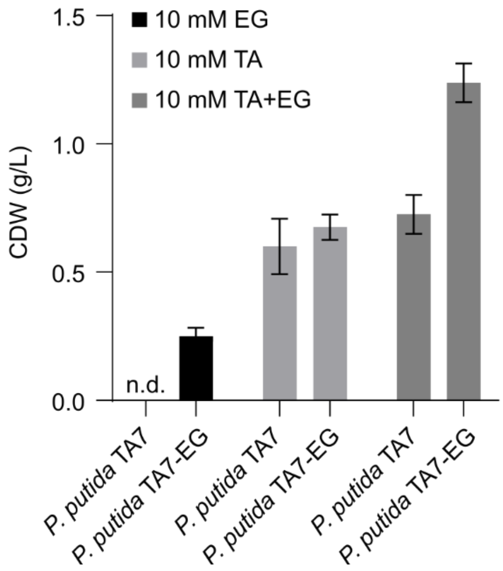


**Supplementary Figure 6: Cell dry weight measurements of *P. putida* TA7 and TA7-EG.** Both strains were cultured for 24 h in M9 minimal medium, pH 7.5, with the indicated PET monomer concentrations. Mean and S.D. of duplicate cultures are shown.

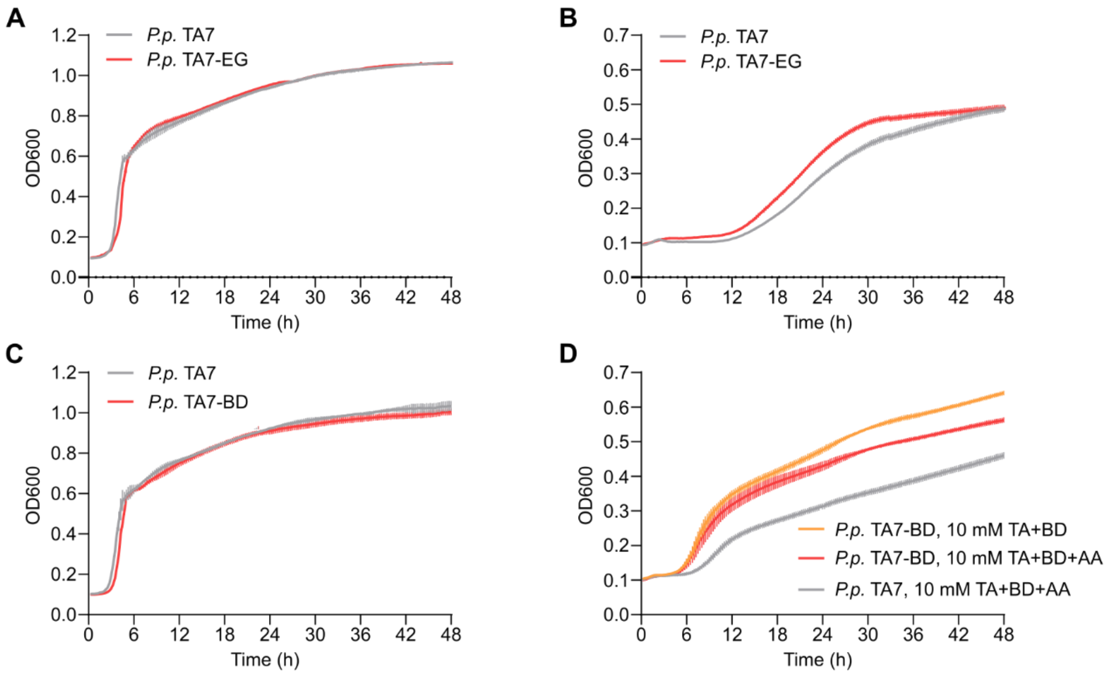


**Supplementary Figure 7: Growth comparisons of *P. putida* TA7, TA7-EG, and TA7-BD in LB and M9 minimal media.** (A) Growth comparison of *P. putida* TA7 and TA7-EG in LB medium. (B) Growth comparison of *P. putida* TA7 and TA7-EG in M9 minimal medium, 10 mM TA, pH 7.5. Despite not showing any mutations in the $tph_{II}$ cassette, strain TA7-EG showed a slight growth advantage over strain TA7 with TA as sole carbon source. (C) Growth comparison of *P. putida* TA7 and TA7-BD in LB medium. (D) Growth comparison of *P. putida* TA7 and TA7-BD in M9 minimal medium, 10 mM TA, 10 mM BD, and with / without 10 mM AA, pH 7.5. Data shown are mean and S.D. from n=6 wells per strain.

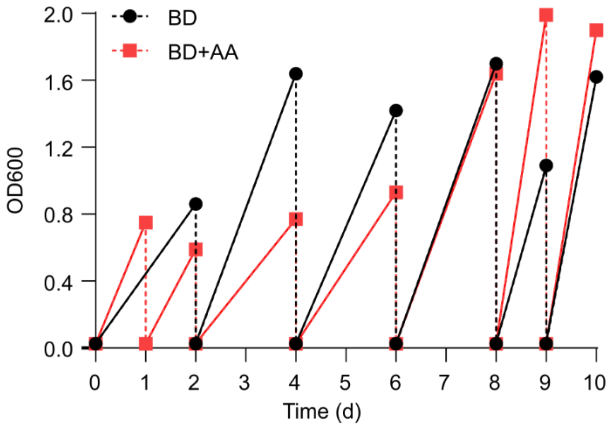


**Supplementary Figure 8: ALE with *P. putida* TA7 for BD utilization**. We performed ALE with *P. putida* TA7 in two cultures using M9 minimal medium containing either 25 mM BD and 5 mM TA, or 25 mM BD, 5 mM TA, and 25 mM AA. Single clones were isolated on day 10 followed by further characterization and whole-genome sequencing, yielding strain *P. putida* TA7-BD. This strain carries a mutation in the LysR-family transcriptional regulator PP_2046 (c.407T>C, V136A) likely influencing BD metabolism.[5]

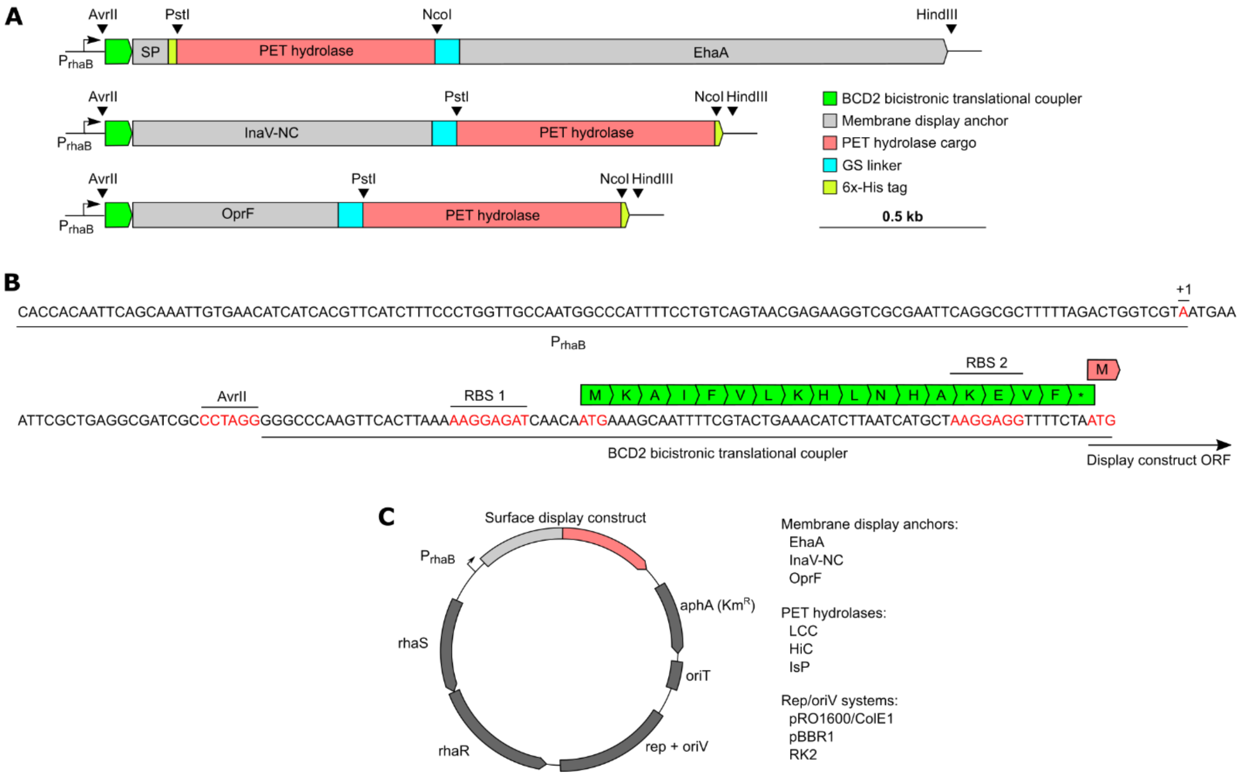


**Supplementary Figure 9: Design of PET hydrolase membrane display expression vectors.** (A) Overview of membrane display expression cassettes, including EhaA (top), InaV-NC (middle), and OprF (bottom). The display cassettes are flanked by unique AvrII and HindIII restriction sites, are under transcriptional control of the $P_{rhaB}$ promoter, and contain unique PstI and NcoI sites to easily clone cargo domains (here: PET hydrolases) into each display anchor system. Note that the cargo domain is positioned N-terminally to the display anchor domain in EhaA, while the cargo sits C-terminally in both InaV-NC and OprF. In all three constructs, cargo and display anchor are connected via a $G_4SGGS(G_4S)_3$ linker, and the PET hydrolase cargos contain a 6x-His tag (N-terminal in EhaA constructs, C-terminal in InaV-NC and OprF constructs). The PstI site was embedded in a 9 bp sequence (cctgcaggc) to maintain mostly neutral amino acids (PAG); likewise, the NcoI site was embedded in a 9 bp sequence (ggccatggg, GHG). (B) Sequence view of the $P_{rhaB}$ promoter-BCD2 region, containing $P_{rhaB}$, AvrII restriction site, BCD2 bicistronic translational coupler, and the ATG of the display construct. (C) Generalized plasmid map of the utilized membrane display vectors; we used three vector systems differing in their replication system, providing low (RK2), medium (pBBR1), and high (pRO1600/ColE1) cellular copy numbers, in combination with the three membrane display anchors and the three PET hydrolases.

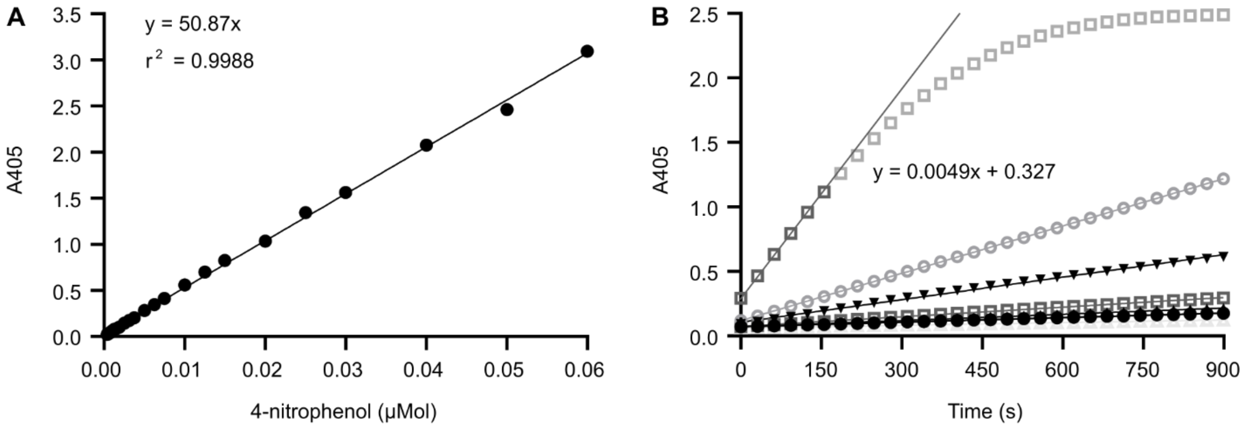


**Supplementary Figure 10: pNPB assay establishment.** (A) Calibration curve for 4-nitrophenol (4NP; see Methods for details). (B) Exemplary experimental data. 4NP formation from pNPB was measured in 96-well plates over 10 min (measurement intervals: 31 s) in a plate reader. Only those data points falling into the linear range of the assay were used to derive the slopes of the curves using linear regression. As an example, the datapoints of the topmost curve used for the calculation (darker grey) together with the resulting equation are indicated. The slope was subsequently used to derive 4NP formation over time using the slope of the calibration curve.

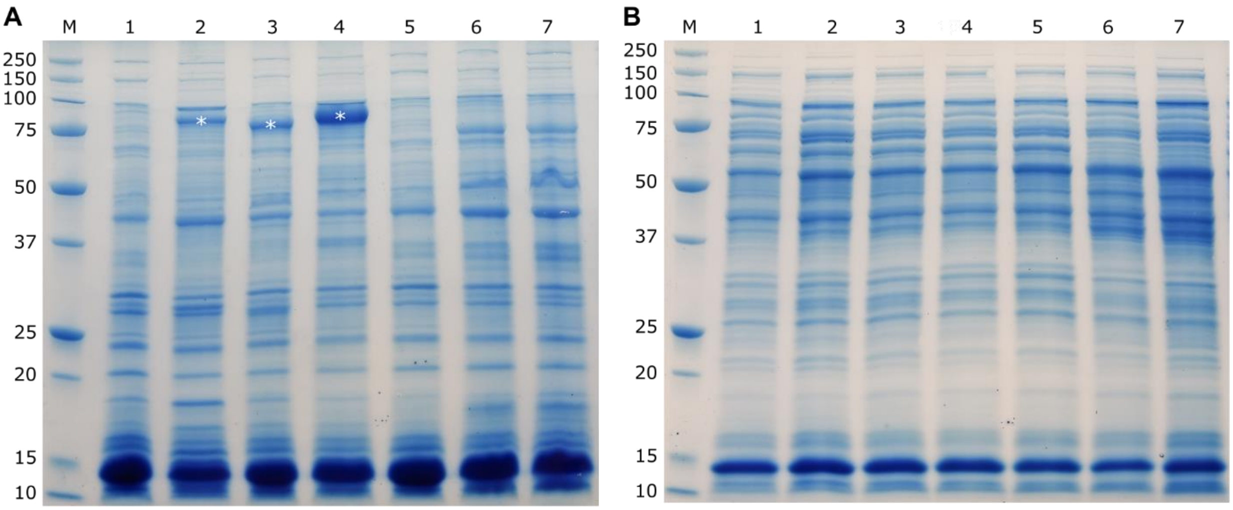


**Supplementary Figure 11: SDS-PAGE analysis of EhaA and OprF membrane display constructs.** (A) SDS-PAGE of the membrane fraction of cells expressing PET hydrolase display constructs. M, marker; 1 pLo-$P_{rhaB}$; 2 pLo-$P_{rhaB}$-EhaA-LCC (85.2 kDa); 3 pLo-$P_{rhaB}$-EhaA-HiC (77.6 kDa), 4 pLo-$P_{rhaB}$-EhaA-IsP (84.8 kDa), 5 pLo-$P_{rhaB}$-OprF-LCC (50.3 kDa), 6 pLo-$P_{rhaB}$-OprF-HiC (42.6 kDa), 7 pLo-$P_{rhaB}$-OprF-IsP (50.0 kDa). For calculating the shown theoretical molecular weights, we assumed that the OprF SP (present in both EhaA and OprF constructs) is cleaved off at position 24 (signal peptidase recognition site AMA). Asterisks mark the putative bands belonging to the EhaA expression constructs, which were excised, and their identity confirmed by mass spectrometry. (B) SDS-PAGE of the cytosolic fraction of cells expressing PET hydrolase display constructs; loading as in (A). No substantial accumulation of the display constructs in the cytosolic fraction was observed.

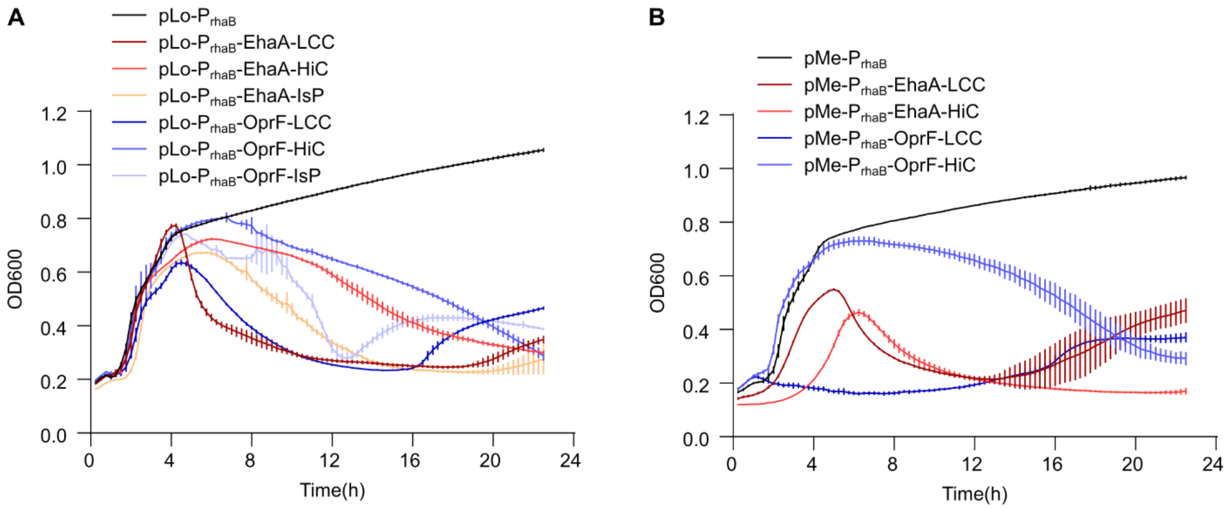


**Supplementary Figure 12: Plate reader analysis of EhaA and OprF PET hydrolase membrane display construct expression growth curves.** Cultures of *P. putida* TA7-EG transformed with the indicated plasmids were grown in LB medium in 96-well plates and OD600 was measured in 15 min intervals for 24 h post-induction (induction with 2 mM rhamnose at t=0 h). (A) Membrane display constructs expressed from low copy number vectors (RK2). (B) Membrane display constructs expressed from medium copy number vectors (pBBR1). Mean and S.D. from a representative experiment with n=6 wells per construct are shown.

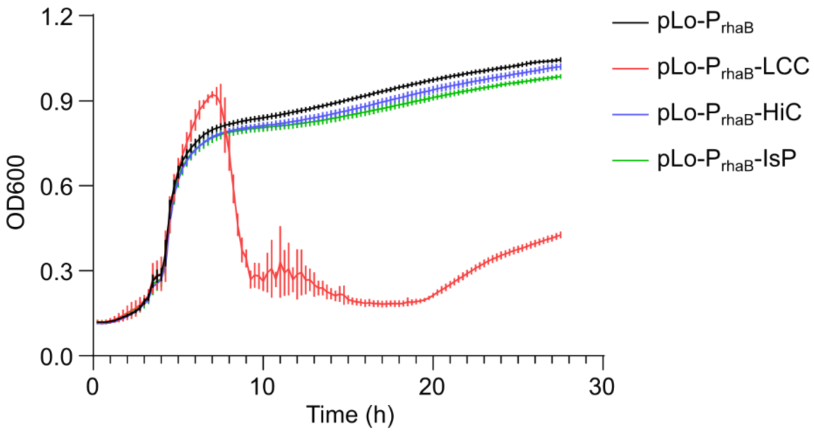


**Supplementary Figure 13: Plate reader analysis of LCC, HiC, and IsP cytosolic expression.** *P. putida* TA7-EG cultures expressing cytosolic constructs of the three PET hydrolases were grown in 96-well plates and OD600 was measured in 15 min intervals for 24 h post-induction (induction at t=0 h). Mean and S.D. from a representative experiment with n=10 wells per construct are shown.

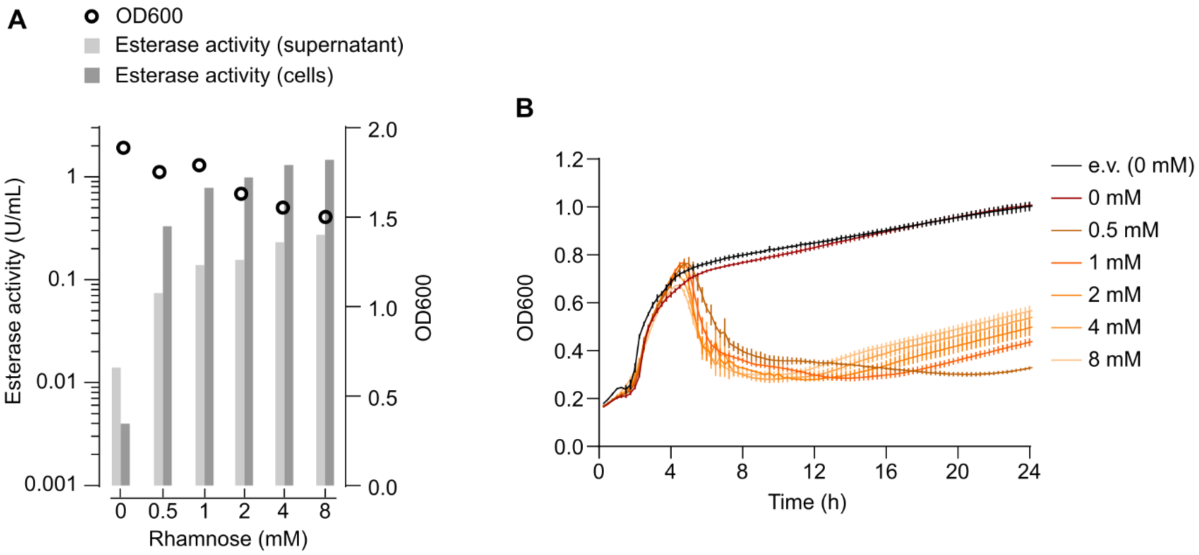


**Supplementary Figure 14: Rhamnose inducer titration with pLo-P$_{rhaB}$-EhaA-LCC.** (A) *P. putida* TA7-EG cultures transformed with pLo-P$_{rhaB}$-EhaA-LCC were induced with the indicated amounts of rhamnose, and OD600 and esterase activity were measured 3 h post-induction (before onset of culture collapse). (B) Plate reader analysis of the same expression culture setup (and including pLo-P$_{rhaB}$ as e.v. control) induced with the indicated amounts of rhamnose; culture OD600 was measured for 24 h post-induction (induction at t=0 h). Data in (A) are derived from a single representative experiment performed with duplicate cultures, while data shown in (B) are derived from a single representative experiment with n=6 wells per strain.

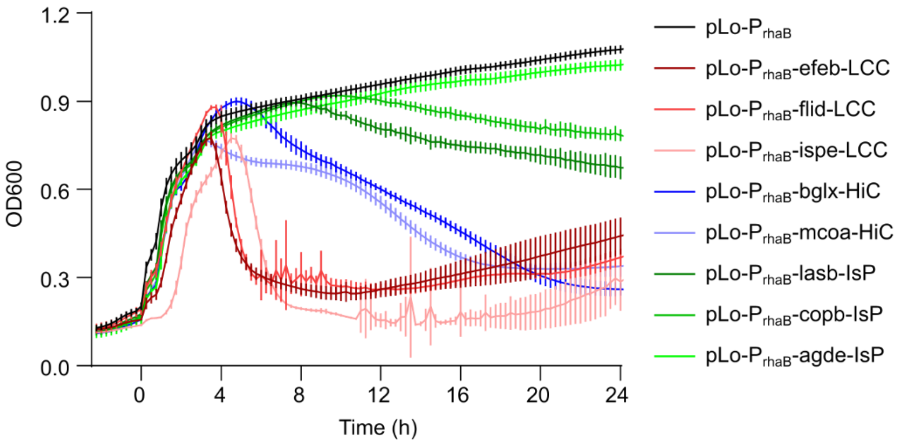

**Supplementary Figure 15: Plate reader analysis of PET hydrolase secretion construct expression growth curves.** *P. putida* TA7-EG cultures transformed with the indicated plasmids were grown in LB medium in 96-well plates and OD600 was measured in 15 min intervals for 24 h post-induction (induction at t=0 h). Mean and S.D. from a representative experiment with n=6 wells per construct are shown.

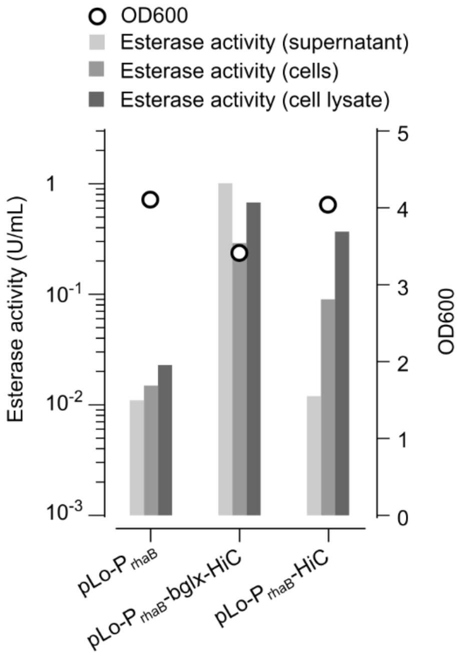


**Supplementary Figure 16: Assessing HiC esterase activity in cell culture supernatant, whole-cell fraction, and cell lysate.** *P. putida* TA7-EG cultures transformed with the indicated plasmids were cultured for 3 h post-induction, and esterase activity in the supernatant, cell fraction, and cell lysate was assessed. We assume that the strong cell-associated esterase signal for the secreted pLo-$P_{rhaB}$-bglx-HiC construct originates from the PET hydrolase *en route* to secretion but still in the cell's periplasm, where it may have access to the pNPB substrate through outer membrane porins. Cytosolic expression does not result in apparent HiC accumulation in the supernatant and a lower cell-associated esterase signal.

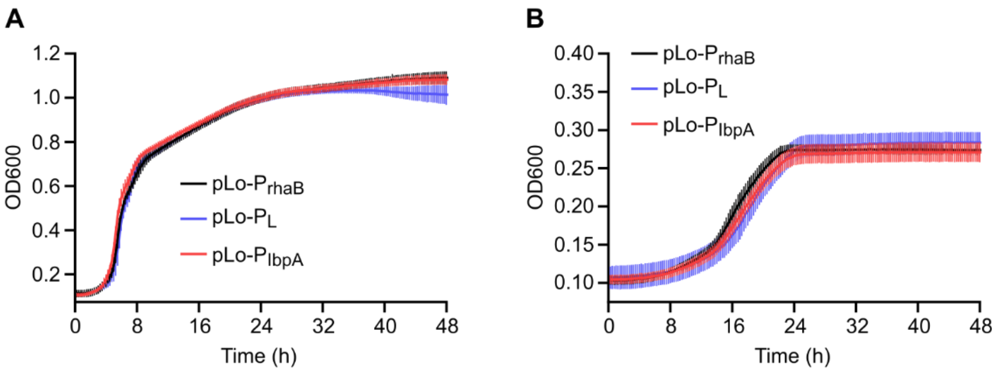

**Supplementary Figure 17: Comparing growth of *P. putida* TA7-EG transformed with different promoter system empty vector plasmids.** (A) Plate reader growth profiles of *P. putida* TA7-EG, transformed with the three promoter systems e.v. plasmids, in LB medium. (B) Same as (A), with growth performed in M9 minimal medium, 10 mM TA, 10 mM EG. Mean and S.D. of n=10 wells per strain are shown.

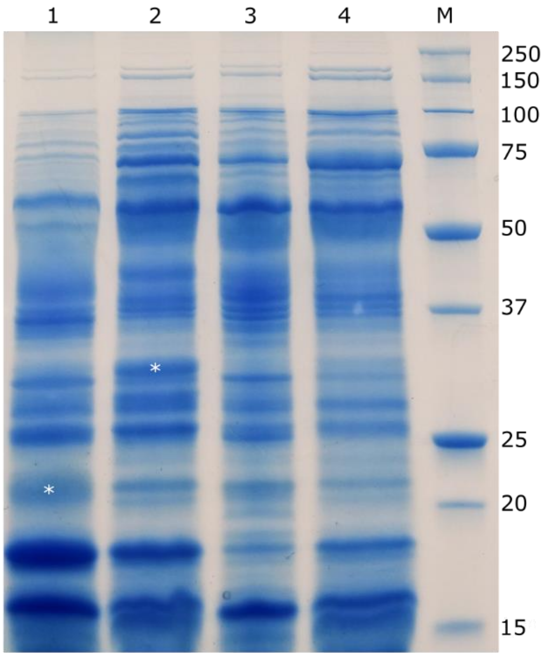


**Supplementary Figure 18: SDS-PAGE analysis of $P_{IbpA}$-bglx-HiC and $P_{IbpA}$-lasb-IsP$^{Dura}$ secretion constructs**. Plasmid-based or genomically integrated $P_{IbpA}$ constructs were evaluated by SDS-PAGE and LC-MS analysis. Cultures grown in LB medium were induced by a 37°C heat-shock for 2 h and subsequently grown at 30°C for 20 h. 50 mL of culture were cleared by centrifugation and the supernatant concentrated in spin filters to ca. 500 µL volume, of which 15 µL were loaded per well. Lane 1: *P. putida* TA7-EG transformed with pLo-$P_{IbpA}$-bglx-HiC (expected MW 22.8 / 20.7 kDa with/without the bglx SP); Lane 2: *P. putida* TA7-EG transformed with pLo-$P_{IbpA}$-bglx-IsP$^{Dura}$ (expected MW 30.4 / 27.9 kDa with/without the lasb SP); Lane 3: *P. putida* TA7-EG with $P_{IbpA}$-bglx-HiC genomic knock-in; Lane 4: *P. putida* TA7-EG wt; M, molecular weight marker. We observed putative bands for the plasmid-based bglx-HiC and lasb-IsP$^{Dura}$ secretion constructs (marked by asterisks). Those bands were excised and analyzed by LC-MS, which confirmed the identity of bglx-HiC but did not unambiguously confirm the identity of lasb-IsP$^{Dura}$. The genomically integrated $P_{IbpA}$-bglx-HiC construct (lane 3) did not show a clear band in the expected size range, likely due to lower expression output compared to plasmid-based expression.

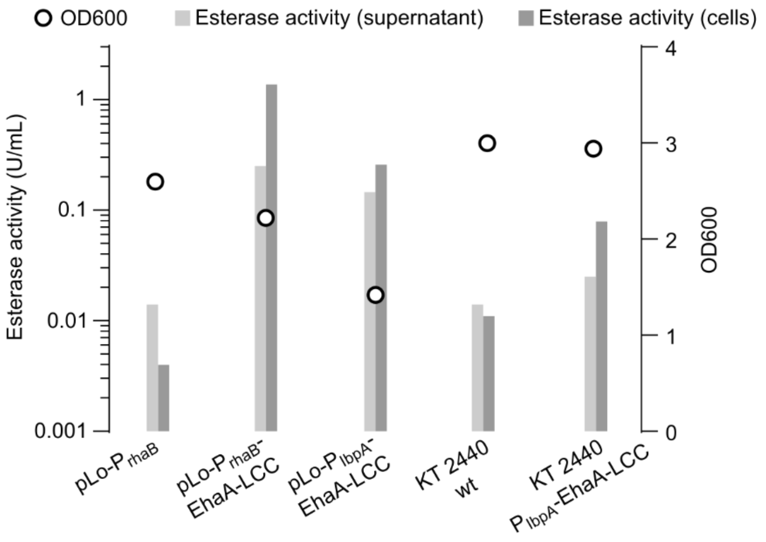


**Supplementary Figure 19: Comparing EhaA-LCC expression output from different promoter systems.** *P. putida* TA7-EG cultures transformed with the indicated plasmids (first three entries) or *P. putida* KT2440 wt and the matching $P_{IbpA}$-EhaA-LCC knock-in strain were compared for EhaA-LCC expression and cell fitness. The two cultures with the $P_{rhaB}$ plasmids were induced with rhamnose and cultured for 3 h before analysis. The $P_{IbpA}$ strains (both plasmid and genomic knock-in, and the KT2440 wt control) were cultured for 2 h at 37° C, followed by a 20 h outgrowth phase at 30° C before analysis. Data shown depict the mean from a single representative experiment with duplicate cultures.

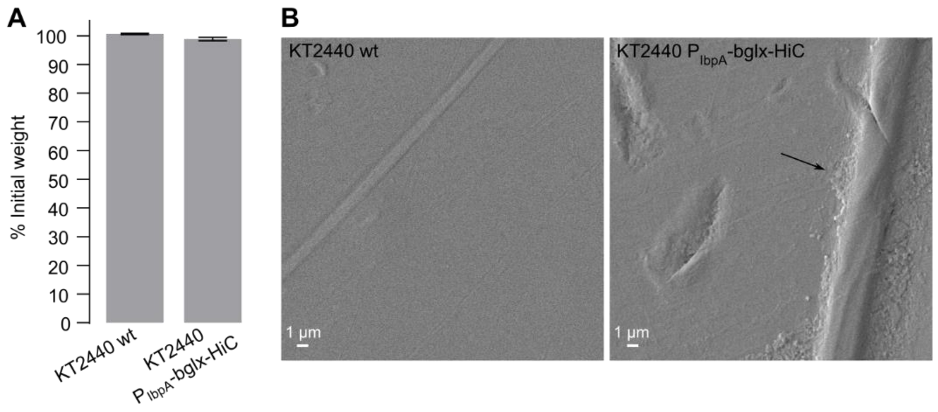


**Supplementary Figure 20: Testing for aPET plastic degradation with *P. putida* KT2440 $P_{IbpA}$-bglx-HiC.** *P. putida* KT2440 wt and the corresponding $P_{IbpA}$-bglx-HiC knock-in strain were cultured in LB medium (15 mL in a 125 mL shake flask) for 6 days with daily medium change and supplemented with aPET film (ca. 3x3 cm). The induction regime consisted of a 2 h heat shock per day at 37° C followed by outgrowth at 25° C. (A) After 6 days, we observed slight fraying of the aPET film along the edges and a weight loss in the range of 1.5% for those samples cultured with *P. putida* $P_{IbpA}$-bglx-HiC. (B) SEM analysis of the film did not reveal major changes in surface structure compared to the wt control, albeit some potential pitting of the surface especially along preexisting edges and scratches.

#### II Supplementary Tables

##### Supplementary Table 1: Genomic landing sites of knock-in cassettes.

| Strain | tphII cassette | $P_{IbpA}$-PET hydrolase cassette | Gene(s) affected |
|---|---|---|---|
| KT2440 | n.a. | n.a. | n.a. |
| TA | 5,251,766 - 5,251,774 | n.a. | PP_4627 (cidR) |
| TA7 | 5,251,766 - 5,251,774 | n.a. | PP_4627 (cidR) |
| KT2440 $P_{IbpA}$-bglx-HiC | n.a. | 6,055,981 - 6,055,989 | PP_5309 (oxyR) |
| KT2440 $P_{IbpA}$-EhaA-LCC | n.a. | 1,310,650 - 1,310,658 | PP_1144 |
| KT2440 $P_{IbpA}$-Iasb-IsP$^{Dura}$ | n.a. | 4,242,500- 4,242,508 | PP_3719 |
| TA7-BD $P_{IbpA}$-bglx-HiC | 5,251,766 - 5,251,774 | 256,383 - 256,391 | PP_4627, PP_0205 |
| TA7-BD $P_{IbpA}$-EhaA-LCC | 5,251,766 - 5,251,774 | 866,370 - 866,378 | PP_4627, PP_0747 |
| TA7-BD $P_{IbpA}$-Iasb-IsP$^{Dura}$ | 5,251,766 - 5,251,774 | 2,401,350 - 2,401,358 | PP_4627, PP_2102 |
| TA7-EG $P_{IbpA}$-bglx-HiC | 5,251,766 - 5,251,774 | 5,323,278 - 5,323,286 | PP_4627, intergenic between PP_4683 and PP_4684 |
| TA7-EG $P_{IbpA}$-EhaA-LCC | 5,251,766 - 5,251,774 | 4,505,576 - 4,505,584 | PP_4627, PP_3997 |

#### Supplementary Table 2: Whole-genome sequencing of engineered *P. putida* strains.

| POS | TYPE | REF | ALT | FTYPE | STRAND | NT_POS | AA_POS | EFFECT | LOCUS | PRODUCT | 1 | 2 | 3 | 4 | 5 | 6 | 7 | 8 | 9 | 10 | 11 | 12 |
|---|---|---|---|---|---|---|---|---|---|---|---|---|---|---|---|---|---|---|---|---|---|---|
| 278742 | ins | T | TA | NA | NA | NA | NA | NA | NA | NA | + | + | + | + | + | + | + | + | + | + | + | + |
| 307865 | ins | G | GC | pseudo | + | NA | NA | intragenic_variant n.307865_307866insC | PP_0253 | pck (pseudogene) | + | + | + | + | + | + | + | + | + | + | + | + |
| 336124 | ins | A | AT | CDS | + | 211/237 | 71/78 | frameshift_variant c.210dupT p.Thr71fs | PP_0278 | hypothetical protein | + | + | + | + | + | + | + | + | + | + | + | + |
| 353067 | snp | A | G | NA | NA | NA | NA | NA | NA | NA | + | + | + | + | + | + | + | + | + | + | + | + |
| 499203 | snp | A | G | NA | NA | NA | NA | NA | NA | NA | + | + | + | + | + | + | + | + | + | + | + | + |
| 1070246 | ins | T | TGA | NA | NA | NA | NA | NA | NA | NA | + | + | + | + | + | + | + | + | + | + | + | + |
| 1126645 | ins | A | AC | NA | NA | NA | NA | NA | NA | NA | + | + | - | + | + | + | + | - | + | + | + | + |
| 1419548 | ins | G | GC | NA | NA | NA | NA | NA | NA | NA | + | + | + | + | + | + | + | + | + | + | + | + |
| 1499480 | mnp | CA | AC | NA | NA | NA | NA | NA | NA | NA | + | + | + | + | + | + | + | + | + | + | + | - |
| 1499480 | complex | CA | AC | NA | NA | NA | NA | NA | NA | NA | - | - | - | - | - | - | - | - | - | - | - | + |
| 1499498 | complex | T | CG | NA | NA | NA | NA | NA | NA | NA | + | + | + | + | + | + | + | + | + | + | + | + |
| 1499506 | ins | G | GC | NA | NA | NA | NA | NA | NA | NA | + | + | + | + | + | + | + | + | + | + | + | + |
| 1932638 | ins | G | GC | NA | NA | NA | NA | NA | NA | NA | + | + | + | + | + | + | + | + | + | + | + | + |
| 2327922 | snp | A | G | CDS | - | 407/924 | 136/307 | missense_variant c.407T>C p.Val136Ala | PP_2046 | LysR family transcriptional regulator | - | - | - | - | - | - | + | + | + | + | - | - |
| 2958523 | snp | C | T | CDS | + | 1283/1494 | 428/497 | missense_variant c.1283C>T p.Ala428Val | PP_2589 | aldehyde dehydrogenase family protein | + | + | + | + | + | + | + | + | + | + | + | + |
| 3319377 | snp | C | T | CDS | - | 769/918 | 257/305 | missense_variant c.769G>A p.Glu257Lys | PP_2919 | LysR family transcriptional regulator | - | - | - | - | + | - | - | - | - | - | - | - |
| 3786089 | snp | G | C | CDS | - | 1120/1242 | 374/413 | missense_variant c.1120C>G p.Leu374Val | PP_3349 | 3-(3-hydroxy-phenyl)propionate transporter MhpT | - | - | + | - | - | - | + | + | + | + | + | + |
| 3799247 | snp | C | T | CDS | + | 151/471 | 51/156 | missense_variant c.151C>T p.Arg51Cys | PP_3359 | hydroxycinnamic acid degradation regulator | - | - | + | - | - | - | + | + | + | + | + | + |
| 3845703 | ins | C | CG | NA | NA | NA | NA | NA | NA | NA | + | + | + | + | + | + | + | + | + | + | + | + |
| 3951159 | complex | TTA | CTT | NA | NA | NA | NA | NA | NA | NA | + | + | + | + | + | + | + | + | + | + | + | + |
| 4586030 | complex | CTGC | TCGCG | NA | NA | NA | NA | NA | NA | NA | + | + | - | + | + | + | + | + | + | + | + | + |
| 4586056 | ins | A | AC | NA | NA | NA | NA | NA | NA | NA | + | + | - | + | + | + | + | + | + | + | + | + |
| 4657540 | snp | G | A | CDS | + | 176/1782 | 59/593 | missense_variant c.176G>A p.Arg59His | PP_4121 | NADH-quinone oxidoreductase subunit C/D | - | - | - | + | - | - | - | - | - | - | - | - |
| 4740804 | del | CT | C | NA | NA | NA | NA | NA | NA | NA | - | + | - | - | - | - | - | - | - | - | - | - |
| 4740811 | complex | TATTTTTCTG | ATTTGTCCT | NA | NA | NA | NA | NA | NA | NA | + | - | + | + | + | + | + | + | + | + | + | + |
| 4740816 | complex | TTCTG | GTCCT | NA | NA | NA | NA | NA | NA | NA | - | + | - | - | - | - | - | - | - | - | - | - |
| 4874145 | snp | C | A | CDS | - | 388/756 | 130/251 | stop_gained c.388G>T p.Glu130* | PP_4283 | GntR family transcriptional regulator | - | - | - | - | - | - | - | - | - | - | + | + |
| 4978562 | del | GC | G | CDS | - | 37/696 | 13/231 | frameshift_variant c.37delG p.Ala13fs | PP_4384 | flagellar L-ring protein | - | - | - | - | - | - | - | - | - | - | + | + |
| 4980585 | ins | G | GGGC | NA | NA | NA | NA | NA | NA | NA | + | + | + | + | + | + | + | + | + | + | + | + |
| 5317300 | ins | G | GT | CDS | - | 347/492 | 116/163 | frameshift_variant c.347_348insA p.Ala117fs | PP_4679 | acetolactate synthase small subunit | - | - | - | - | - | - | - | - | - | - | - | + |
| 5555608 | ins | G | GGCC | NA | NA | NA | NA | NA | NA | NA | + | + | + | + | + | + | + | + | + | + | + | + |
| 5674752 | complex | CG | GCC | NA | NA | NA | NA | NA | NA | NA | + | + | + | + | + | + | + | + | + | + | + | + |
| 5681415 | ins | C | CCGGG | NA | NA | NA | NA | NA | NA | NA | + | + | + | + | + | + | + | + | + | + | + | + |
| 5871461 | ins | T | TA | CDS | - | 1191/2280 | 397/759 | frameshift_variant c.1191_1192insT p.Lys398fs | PP_5145 | phosphoenolpyruvate-dependent regulator | - | - | - | + | - | - | - | - | - | - | - | - |
| 6013885 | snp | T | C | NA | NA | NA | NA | NA | NA | NA | + | + | + | + | + | + | + | + | + | + | + | + |

**1** *P. putida* KT2440; **2** *P. putida* TA; **3** *P. putida* TA7; **4** *P. putida* KT2440 $P_{IbpA}$-bglx-HiC; **5** *P. putida* KT2440 $P_{IbpA}$-EhaA-LCC; **6** *P. putida* KT2440 $P_{IbpA}$-lasb-IsP$^{Dura}$; **7** *P. putida* TA7-BD; **8** *P. putida* TA7-BD $P_{IbpA}$-bglx-HiC; **9** *P. putida* TA7-BD $P_{IbpA}$-EhaA-LCC; **10** *P. putida* TA7-BD $P_{IbpA}$-lasb-IsP$^{Dura}$; **11** *P. putida* TA7-EG $P_{IbpA}$-bglx-HiC; **12** *P. putida* TA7-EG $P_{IbpA}$-EhaA-LCC

All mutations present in the reference strain **1** and all other strains (compared against the reference genome) are greyed out; random mutations (potentially sequencing artefacts) are shown in yellow; functional mutations are shown in red. + / - signs indicate presence or absence, respectively, of a given mutation.

#### Supplementary Table 3: Signal peptide oligonucleotide library.

| Oligonucleotide sequence (5’ → 3’)[a] | Signal peptide protein sequence[b] | Origin (GenBank Nr.), 4-letter code[c] |
|---|---|---|
| catgctaaggaggtCttctaATGCAGTCGAAAACCACCCGTCGTTCGTTCGTGAAGGGTCTGGCGGCAACGGGTTTGTTGGGCGGGCTCGGTATGTGGCGTGCCCCAGTGTGGGCAGTCGCTcctgcaggcCAATCCAATC | MQSKTTRRSFVKGLAATGLLGGLGMWRAPVWAVA | *P. putida*, copper resistance protein A (AAN70945.1)<br>**copa** |
| catgctaaggaggtCttctaATGGGGGAGCATGATCTGAATCGCCGTCAGTTTATTAAAACCGTCGGTGTGGCCTCGGTCGCCGCTGCCGCGATGTCGCTCCCATTTGTGAAAGCAAATGCAcctgcaggcCAATCCAATC | MGEHDLNRRQFIKTVGVASVAAAAMSLPFVKANA | *P. putida*, HA62_13390 (KEX93498.1)<br>**spat** |
| catgctaaggaggtCttctaATGTGTATGGACGATTGCTGTTCCTCCAGCTCGCGCCGCGACTTTCTCAAACTGGGTGCAATGTTGACGGCTGCCGGCGCTCTGCCGCTGCTCTCGTCCCTCCAGGCACGTGCCcctgcaggcCAATCCAATC | MCMDDCCSSSSRRDFLKLGAMLTAAGALPLLSSLQARA | *P. putida*, ABC transporter substrate binder (AJA16455.1)<br>**atsb** |
| catgctaaggaggtCttctaATGCCACCTACCCCAACGCGCCGCACGTTCGTCAAAGGCTTGGGTGCAGCTACCGCACTGGCAGGTCTGGGGCTCTGGCGCCCGTTGGCACAAGCCcctgcaggcCAATCCAATC | MPPTPTRRTFVKGLGAATALAGLGLWRPLAQA | *P. putida*, multicopper oxidase (AMK30412.1)<br>**muox** |
| catgctaaggaggtCttctaATGTCCAATCGTGAAATTTCGCGCCGGTCCTTTTTGCAAGGTGGGTTGGTGGCTGGTGTGTCCGTGACCCTCACGCCACTGTCCTCGCAAGCACTGGCGcctgcaggcCAATCCAATC | MSNREISRRSFLQGGLVAGVSVTLTPLSSQALA | *P. putida*, Tat pathway signal sequence (SDB06516.1)<br>**tpss** |
| catgctaaggaggtCttctaATGAATGACTCCGAACATTTCAACCTCCAACGGCGGCGGGTGCTGATGGGTATGGGGGCAGCCGGTGTGGCCTTGGCTGGTACGGCTTTGTCGTGTCCTGCAATGGCTcctgcaggcCAATCCAATC | MNDSEHFNLQRRRVLMGMGAAGVALAGTALSCPAMA | *P putida*, deferrochelatase / peroxidase EfeB (SDB47198.1)<br>**efeb** |
| catgctaaggaggtCttctaATGCCATCGCGCCGCACCTTCTTGCAGCTCTCCCTGGCTGCGGGTCTCGGTGCTAGCCTGGGGCTCCCACTGGGTCTCGCTCGTGCAcctgcaggcCAATCCAATC | MPSRRTFLQLSLAAGLGASLGLPLGLARA | *P. putida*, agmatine deiminase (APO84336.1)<br>**agde** |
| catgctaaggaggtCttctaATGGCCGATGCCATGGCGAGCAGCGTGCATCGCCGTGAAGTGATTGGGTGGATGGGTATCGGCGCATTGGCGcctgcaggcCAATCCAATC | MADAMASSVHRREVIGWMGIGALA | *P. putida*, uxpA (AAN66669.1)<br>**uxpa** |
| catgctaaggaggtCttctaATGAGCCGGGACACGGGGGACAATTTGGACCGTAATCACAGCGGTAACATGCCTATGGCCAATGTCATGGACGCCTATCTCTCCCGCCGGAGCATTATGCGCGGCTCCCTCGGGGCTGCTATCGCAcctgcaggcCAATCCAATC | MSRDTGDNLDRNHSGNMPMANVMDAYLSRRSIMRGSLGAAIA | *P. putida*, uxpB (CAA56976.1)<br>**uxpb** |

| | | |
|---|---|---|
| catgctaaggaggtCttctaATGAAAACGCCAGCGGTCTCCCATGCTTCCCGTCGTGCATTTCAACTGACCAGCGTGACCGCGTTGATGATTAGCTTGGGGCTCATTACCGCTATGGCCcctgcaggcCAATCCAATC | MKTPAVSHASRRAFQLTSVTALMISLGLITAMA | *P. putida*, Mn(II) copper oxidase A, McoA (AAN68792.1) **mcoa** |
| catgctaaggaggtCttctaATGAGCAATCGGGATATTTCCCGTCGGGCGTTCCTCCAAGGTGGCCTGATTGCGGGCGTGGGGGTGACGATGGCGCCTCTGGGTAGCCAAGCCTTTGCTcctgcaggcCAATCCAATC | MSNRDISRRAFLQGGLIAGVGVTMAPLGSQAFA | *P. putida*, molybdopterin binding subunit (WP_159411419.1) **mobs** |
| catgctaaggaggtCttctaATGATGAAATTGAGCTTGCTCGGTCTCGCGATGGGTCTGGCGTCCCAAGCTGCACTCGCTGCTcctgcaggcCAATCCAATC | MMKLSLLGLAMGLASQAALAA | *Pseudomonas*, betaglucosidase BglX (WP_010952511.1) **bglx** |
| catgctaaggaggtCttctaATGACCACGAAAACGTCCATTGCAAAAGCCTTGACGCTCGCAGCAGGGTTGTCGCTGGCTTCGATGCAAGCGTTTGCGcctgcaggcCAATCCAATC | MTTKTSIAKALTLAAGLSLASMQAFA | *P. putida*, alginate biosynthesis protein AlgF (AAN66902.1) **algf** |
| catgctaaggaggtCttctaATGCTCCGTGCCTTGATCTTGTTGGCCTTGAGCTGTCTCCTGAGCCCCGCATTCGCGcctgcaggcCAATCCAATC | MLRALILLALSCLLSPAFA | *Pseudomonas*, murein hydrolase activator EnvC (WP_010955607.1) **envc** |
| catgctaaggaggtCttctaATGCTCCGTGCTACGGCTTTGGCCCTCGCATGCTTGGCTAGCTCGAGCGCAATCGCTGCCcctgcaggcCAATCCAATC | MLRATALALACLASSSAIAA | *Pseudomonas*, alpha/beta hydrolase (WP_010953184.1) **abhy** |
| catgctaaggaggtCttctaATGACGAATTCGTTGGCTCGTCCTAGCTTGTTGGCCCTGACCGTGTCGTTTTCGATGCTGGGGGCGGCAGCACCATCGTTTGCGGCCcctgcaggcCAATCCAATC | MTNSLARPSLLALTVSFSMLGAAAPSFAA | *Pseudomonas*, copper resistance protein B (WP_049588384.1) **copb** |
| catgctaaggaggtCttctaATGAATCTGTTGAAACCCTTGACGCCTAGCCTCCTCGCATTGGCCTTGGCTGCAcctgcaggcCAATCCAATC | MNLLKPLTPSLLALALAA | *Pseudomonas*, TolC family (WP_010954293.1) **tolc** |
| catgctaaggaggtCttctaATGTTTGCAAAAGCCGTGGCAGTGTCCCTGTTGACCCTGGCAAGCGCAAGCGTGTTCGCCGCAcctgcaggcCAATCCAATC | MFAKAVAVSLLTLASASVFAA | *Pseudomonas*, azurin (WP_003249580.1) **azur** |
| catgctaaggaggtCttctaATGAACATCGTCCACAAAGCATTGACGACCTCGTTGTTGGCTCTGTCGGTGTCCTCGGCTTTTGCCGCTGGTTCCCCTGGCcctgcaggcCAATCCAATC | MNIVHKALTTSLLALSVSSAFAAGSPG | *P. putida*, lipase (putative) (Q88GB2) **lipu** |
| catgctaaggaggtCttctaATGAAACTGAAAAACACCTTGGGCTTGGCCATTGGTTCGCTTGTAGCCGCCACTTCGATTGGCGCTATGGCAcctgcaggcCAATCCAATC | MKLKNTLGLAIGSLVAATSIGAMA | *P. putida*, OmpA family protein (WP_119713436.1) **ompa** |
| catgctaaggaggtCttctaATGCCCTTTCATATCCATCGTGCGACCTGTTGGGCACTGACCTGCTGCCTCTCGGGGCTCTTGGCTGCACCTCAAGCATTGGCAcctgcaggcCAATCCAATC | MPFHIHRATCWALTCCLSGLLAAPQALA | *P. putida*, secretion protein (AHZ75903.1) **secp** |

| | | |
|---|---|---|
| catgctaaggaggtCttcta<u>ATGGCGAGCACCACCGTCTCCGGCATTGGCTCGGGTGTCGATACGCAATCGATTGTCAAAGCGTTGGCC</u>cctgcaggcCAATCCAATC | MASTTVSGIGSGVDTQSIVKALA | *P. putida*, flagellar filament capping protein FliD (AAN69954.1) **flid** |
| catgctaaggaggtCttcta<u>ATGAATCAATCGCTCGGCGTGCTGCGTCTCACCCGTGGCGTCATTGCTCTCGCTTTGGCATCGGTGGCT</u>cctgcaggcCAATCCAATC | MNQSLGVLRLTRGVIALALASVA | *Pseudomonas*, oxidized polyvinyl alcohol hydrolase PvaB (BAA94192.1) **pvab** |
| catgctaaggaggtCttcta<u>ATGCAACAAAACATCGAACGCAACCAAGTGAGCATGACCACGTCCCGTTTTGTGTGGGGCGCTGTGATGGCGCTGGTCGCCCTCGGTTCCGCGTCCGCTGCT</u>cctgcaggcCAATCCAATC | MQQNIERNQVSMTTSRFVWGAVMALVALGSASAA | *Pseudomonas*, PQQ-dependent polyvinyl alcohol dehydrogenase PvaA (BAA09321.1) **pvaa** |
| catgctaaggaggtCttcta<u>ATGAATACCTTTTCCAAGGTCCTCACCGGCACGTTGTTGGCGATGAGCATCAGCAACGCGTTTGCC</u>cctgcaggcCAATCCAATC | MNTFSKVLTGTLLAMSISNAFA | *Pseudomonas mandelii*, EstK (AEW10549.2) **estk** |
| catgctaaggaggtCttcta<u>ATGAACAAGAACAAAACCTTCCTCGCGGCAGCGTTGGTGGCATTGGCAGCGTCGTTTCCTGTCCACGCTGCC</u>cctgcaggcCAATCCAATC | MNKNKTFLAAALVALAASFPVHAA | *Pseudomonas aeruginosa*, lipase lipC (AAG08198.1) **lipc** |
| catgctaaggaggtCttcta<u>ATGAAAAAGGTCTCGACGCTGGACCTGTTGTTTGTGGCTATCATGGGTGTCTCCCCGGCGGCCTTCGCGGCT</u>cctgcaggcCAATCCAATC | MKKVSTLDLLFVAIMGVSPAAFAA | *Pseudomonas aeruginosa*, elastase (AFM37281.1) **lasb** |
| catgctaaggaggtCttcta<u>ATGCACAAACGCACCTATTTGAATGCGTGCTTGGTGCTGGCGTTGGCTGCTGGTGCCTCCCAAGCCTTGGCTGCTCCTGGCGCA</u>cctgcaggcCAATCCAATC | MHKRTYLNACLVLALAAGASQALAAPGA | *Pseudomonas aeruginosa*, protease IV (ABC73074.1) **prpl** |
| catgctaaggaggtCttcta<u>ATGAACTTTCCCCGCGCTTCCCGCCTGATGCAGGCCGCCGTTCTCGGCGGGCTGATGGCCGTGTCGGCCGCCGCCACCGCC</u>cctgcaggcCAATCCAATC | MNFPRASRLMQAAVLGGLMAVSAAATA | *Ideonella sakaiensis*, PETase (GAP38373.1) **ispe** |
| catgctaaggaggtCttcta<u>ATGCAAACCACGGTCACCACCATGTTGTTGGCTTCGGTCGCTCTCGCTGCG</u>cctgcaggcCAATCCAATC | MQTTVTTMLLASVALAA | *Ideonella sakaiensis*, MHETase (GAP38911.1) **ismh** |

[a] The SP coding sequence is underlined; the remaining overhangs were included for cloning purposes.

[b] The first 11 SPs are classified as Tat-dependent, the remaining 19 SPs as Sec-dependent.

[c] The provided 4-letter code is not the official gene name, but a custom naming to identify the different SPs within this manuscript.

**Supplementary Table 4: SP library screening data.**

**LCC**

| Clone | pNPB | OD600 | SP |
|---|---|---|---|
| A4 | 3.28 | 0.471 | **lipu** |
| F1 | 3.277 | 0.256 | **ompa** |
| E1 | 3.256 | 0.286 | **ispe** |
| H9 | 3.23 | 0.235 | **ispe** |
| D3 | 3.181 | 0.327 | **ispe** |
| C9 | 3.177 | 0.283 | **flid** |
| B10 | 3.135 | 0.268 | |
| D8 | 3.087 | 0.243 | |
| F8 | 2.958 | 0.266 | |
| E7 | 2.947 | 0.265 | |
| G11 | 2.934 | 0.251 | |
| H12 | 2.866 | 0.277 | |
| F10 | 2.856 | 0.262 | |
| C11 | 2.854 | 0.542 | **flid** |
| A3 | 2.784 | 0.45 | **efeb** |
| C2 | 2.613 | 0.346 | **efeb** |
| G2 | 2.569 | 0.255 | |
| F7 | 2.553 | 0.255 | |
| D7 | 2.542 | 0.23 | |
| B12 | 2.429 | 0.278 | |
| F4 | 2.426 | 0.289 | |
| F12 | 2.339 | 0.248 | |
| A9 | 2.181 | 0.259 | |
| D11 | 2.04 | 0.264 | |
| G1 | 1.992 | 0.251 | |
| A5 | 1.988 | 0.238 | |
| G8 | 1.93 | 0.191 | |
| C12 | 1.897 | 0.391 | **copa** |
| B9 | 1.884 | 0.253 | |
| C4 | 1.769 | 0.25 | |
| C5 | 1.749 | 0.263 | |
| G4 | 1.712 | 0.268 | |
| A12 | 1.674 | 0.258 | |
| D1 | 1.648 | 0.262 | |
| A7 | 1.608 | 0.247 | |
| G6 | 1.573 | 0.299 | |
| D2 | 1.505 | 0.267 | |
| E8 | 1.475 | 0.253 | |
| D5 | 1.441 | 0.268 | |
| E4 | 1.423 | 0.278 | |
| H7 | 1.422 | 0.269 | |
| A10 | 1.413 | 0.273 | |
| D9 | 1.366 | 0.268 | |
| G3 | 1.348 | 0.278 | |
| F3 | 1.337 | 0.288 | |
| C7 | 1.323 | 0.253 | |
| A2 | 1.305 | 0.292 | |
| F2 | 1.303 | 0.275 | |
| H5 | 1.276 | 0.285 | |
| C10 | 1.271 | 0.259 | |
| H10 | 1.235 | 0.258 | |
| F9 | 1.227 | 0.289 | |

**HiC**

| Clone | pNPB | OD600 | SP |
|---|---|---|---|
| C8 | 2.324 | 0.181 | **bglx** |
| A6 | 2.281 | 0.183 | **bglx** |
| H11 | 2.265 | 0.185 | **azur** |
| E12 | 2.239 | 0.177 | **lipc** |
| H10 | 2.212 | 0.189 | **bglx** |
| C7 | 2.199 | 0.194 | **bglx** |
| B12 | 2.182 | 0.273 | **bglx** |
| F1 | 2.145 | 0.2 | **n.a.** |
| C11 | 2.103 | 0.175 | |
| C4 | 2.083 | 0.188 | |
| C10 | 2.047 | 0.174 | |
| G12 | 2.041 | 0.18 | |
| A8 | 1.98 | 0.176 | |
| H4 | 1.948 | 0.189 | |
| H12 | 1.944 | 0.222 | **mcoa** |
| E9 | 1.9 | 0.191 | |
| F12 | 1.845 | 0.193 | |
| E11 | 1.828 | 0.205 | |
| D3 | 1.767 | 0.19 | |
| H2 | 1.753 | 0.203 | |
| H1 | 1.739 | 0.197 | |
| G4 | 1.72 | 0.189 | |
| B1 | 1.713 | 0.179 | |
| A10 | 1.651 | 0.195 | |
| F5 | 1.582 | 0.192 | |
| G7 | 1.549 | 0.203 | |
| F4 | 1.546 | 0.189 | |
| C2 | 1.54 | 0.187 | |
| B3 | 1.535 | 0.186 | |
| E2 | 1.499 | 0.205 | |
| G6 | 1.494 | 0.189 | |
| D6 | 1.487 | 0.2 | |
| E10 | 1.447 | 0.19 | |
| F6 | 1.438 | 0.201 | |
| E8 | 1.41 | 0.182 | |
| B4 | 1.385 | 0.306 | **prpl** |
| C9 | 1.373 | 0.185 | |
| G8 | 1.346 | 0.201 | |
| G9 | 1.311 | 0.199 | |
| E5 | 1.308 | 0.199 | |
| D9 | 1.234 | 0.183 | |
| F10 | 1.178 | 0.185 | |
| C1 | 1.116 | 0.215 | |
| G1 | 1.066 | 0.222 | |
| D1 | 0.988 | 0.229 | |
| F7 | 0.98 | 0.26 | |
| E1 | 0.884 | 0.239 | |
| F9 | 0.877 | 0.228 | |
| E4 | 0.843 | 0.24 | |
| G10 | 0.84 | 0.253 | |
| F2 | 0.835 | 0.198 | |
| C6 | 0.831 | 0.227 | |

**IsP**

| Clone | pNPB | OD600 | SP |
|---|---|---|---|
| B6 | 2.773 | 0.434 | **pvaa** |
| C12 | 2.763 | 0.404 | **lipc** |
| B8 | 2.716 | 0.361 | |
| B4 | 2.705 | 0.359 | |
| C7 | 2.694 | 0.396 | |
| C11 | 2.676 | 0.371 | |
| A12 | 2.673 | 0.391 | |
| E11 | 2.589 | 0.387 | |
| B11 | 2.576 | 0.599 | **bglx** |
| C1 | 2.571 | 0.432 | **prpl** |
| B10 | 2.505 | 0.617 | **n.a.** |
| A11 | 2.502 | 0.844 | **agde** |
| A4 | 2.495 | 0.652 | **lasb** |
| A5 | 2.49 | 0.324 | |
| A6 | 2.478 | 0.892 | **copb** |
| F8 | 2.472 | 0.337 | |
| C6 | 2.458 | 0.518 | **lasb** |
| H6 | 2.458 | 0.308 | |
| G12 | 2.457 | 0.355 | |
| B9 | 2.447 | 0.561 | **copb** |
| B12 | 2.447 | 0.771 | **copa** |
| D12 | 2.446 | 0.507 | |
| B5 | 2.437 | 0.331 | |
| A8 | 2.43 | 0.697 | **pvab** |
| D5 | 2.427 | 0.374 | |
| A10 | 2.423 | 0.355 | |
| G2 | 2.419 | 0.267 | |
| D8 | 2.401 | 0.843 | **lasb** |
| C8 | 2.4 | 0.661 | |
| A7 | 2.399 | 0.337 | |
| E12 | 2.393 | 0.871 | **agde** |
| A3 | 2.389 | 1.258 | |
| A9 | 2.389 | 0.5 | |
| C4 | 2.386 | 0.494 | |
| D7 | 2.376 | 0.381 | |
| G10 | 2.376 | 0.348 | |
| E10 | 2.374 | 0.711 | |
| C10 | 2.372 | 0.768 | |
| C2 | 2.37 | 0.317 | |
| D2 | 2.365 | 0.434 | |
| E9 | 2.354 | 0.689 | |
| D6 | 2.349 | 0.371 | |
| C9 | 2.348 | 0.352 | |
| F6 | 2.343 | 0.389 | |
| B3 | 2.336 | 0.862 | |
| B7 | 2.334 | 0.851 | |
| D10 | 2.328 | 0.771 | |
| F9 | 2.328 | 0.855 | |
| D11 | 2.322 | 0.342 | |
| F12 | 2.315 | 0.861 | |
| E7 | 2.311 | 0.803 | |
| E6 | 2.307 | 0.547 | |

| | | | |
|---|---|---|---|
| B11 | 1.202 | 0.209 | |
| H3 | 1.162 | 0.259 | |
| B5 | 1.121 | 0.268 | |
| B4 | 1.118 | 0.302 | **uxpb** |
| E2 | 1.115 | 0.29 | |
| A11 | 1.087 | 0.327 | **uxpb** |
| G12 | 1.077 | 0.252 | |
| F11 | 1.064 | 0.299 | |
| E3 | 1.015 | 0.287 | |
| B6 | 0.958 | 0.296 | |
| B3 | 0.935 | 0.256 | |
| H4 | 0.853 | 0.255 | |
| E10 | 0.694 | 0.321 | |
| H1 | 0.647 | 0.311 | |
| H2 | 0.628 | 0.354 | |
| H6 | 0.574 | 0.347 | |
| B7 | 0.542 | 0.365 | |
| B1 | 0.54 | 0.581 | |
| G9 | 0.526 | 0.315 | |
| H11 | 0.519 | 0.362 | |
| D10 | 0.515 | 0.364 | |
| C1 | 0.507 | 0.378 | |
| E12 | 0.501 | 0.341 | |
| G5 | 0.492 | 0.329 | |
| A8 | 0.489 | 0.304 | |
| B8 | 0.482 | 0.283 | |
| F5 | 0.478 | 0.353 | |
| A6 | 0.456 | 0.54 | |
| C8 | 0.456 | 0.334 | |
| G10 | 0.454 | 0.335 | |
| G7 | 0.437 | 0.387 | |
| C6 | 0.428 | 0.37 | |
| D4 | 0.388 | 0.752 | |
| E9 | 0.387 | 0.612 | |
| A1 | 0.38 | 0.816 | |
| D12 | 0.38 | 0.763 | |
| F6 | 0.376 | 0.777 | |
| D6 | 0.374 | 0.607 | |
| C3 | 0.372 | 0.73 | |
| E5 | 0.367 | 0.733 | |
| B2 | 0.364 | 0.772 | |
| E11 | 0.358 | 0.756 | |
| E6 | 0.278 | 0.206 | |
| H8 | 0.267 | 0.288 | |

| | | |
|---|---|---|
| E3 | 0.812 | 0.194 |
| B9 | 0.731 | 0.224 |
| D2 | 0.662 | 0.154 |
| G5 | 0.649 | 0.163 |
| H8 | 0.624 | 0.385 |
| B5 | 0.554 | 0.207 |
| A3 | 0.551 | 0.234 |
| A12 | 0.543 | 0.171 |
| H3 | 0.541 | 0.222 |
| H7 | 0.526 | 0.407 |
| E6 | 0.511 | 0.231 |
| A5 | 0.483 | 0.2 |
| D10 | 0.453 | 0.222 |
| G3 | 0.451 | 0.229 |
| H9 | 0.449 | 0.23 |
| B11 | 0.437 | 0.225 |
| G2 | 0.437 | 0.536 |
| D5 | 0.432 | 0.225 |
| C5 | 0.425 | 0.224 |
| B7 | 0.413 | 0.23 |
| F3 | 0.407 | 0.22 |
| F8 | 0.341 | 0.32 |
| H6 | 0.334 | 0.632 |
| D11 | 0.327 | 0.309 |
| A9 | 0.323 | 0.721 |
| A4 | 0.321 | 0.272 |
| A7 | 0.315 | 0.501 |
| D7 | 0.314 | 0.541 |
| B2 | 0.31 | 0.741 |
| E7 | 0.308 | 0.656 |
| B6 | 0.305 | 0.617 |
| D8 | 0.3 | 0.667 |
| H5 | 0.3 | 0.546 |
| B10 | 0.298 | 0.509 |
| A11 | 0.296 | 0.419 |
| C12 | 0.292 | 0.761 |
| C3 | 0.291 | 0.731 |
| B8 | 0.289 | 0.642 |
| A1 | 0.288 | 0.75 |
| D4 | 0.287 | 0.722 |
| A2 | 0.286 | 0.638 |
| F11 | 0.285 | 0.56 |
| G11 | 0.279 | 0.996 |
| D12 | 0.269 | 0.735 |

| | | |
|---|---|---|
| E8 | 2.304 | 0.397 |
| C5 | 2.3 | 0.893 |
| F11 | 2.289 | 0.836 |
| D9 | 2.264 | 0.5 |
| G11 | 2.262 | 0.625 |
| H10 | 2.262 | 0.362 |
| B1 | 2.249 | 0.876 |
| F10 | 2.245 | 0.504 |
| E4 | 2.2 | 0.301 |
| A2 | 2.199 | 0.763 |
| F1 | 2.199 | 0.861 |
| F7 | 2.192 | 0.858 |
| D4 | 2.188 | 0.872 |
| F4 | 2.185 | 0.573 |
| F2 | 2.182 | 0.791 |
| F5 | 2.18 | 0.363 |
| G9 | 2.179 | 0.885 |
| D1 | 2.174 | 0.311 |
| G6 | 2.174 | 0.885 |
| E2 | 2.168 | 0.897 |
| A1 | 2.156 | 0.782 |
| C3 | 2.149 | 0.894 |
| E5 | 2.141 | 0.296 |
| H12 | 2.127 | 0.355 |
| D3 | 2.126 | 0.294 |
| G8 | 2.119 | 0.864 |
| G3 | 2.112 | 0.777 |
| G5 | 2.093 | 0.844 |
| E3 | 2.083 | 0.284 |
| G7 | 2.078 | 0.366 |
| G1 | 2.072 | 0.666 |
| B2 | 2.061 | 0.887 |
| E1 | 2.058 | 0.269 |
| G4 | 2.056 | 0.833 |
| H11 | 2.054 | 0.624 |
| H1 | 2.042 | 0.815 |
| F3 | 2.038 | 0.308 |
| H7 | 2.025 | 0.871 |
| H2 | 1.955 | 0.356 |
| H4 | 1.955 | 0.851 |
| H9 | 1.9 | 0.374 |
| H3 | 1.896 | 0.863 |
| H8 | 1.832 | 0.375 |
| H5 | 1.813 | 0.341 |

pNPB esterase activity was measured 3 h post-induction; OD600 was measured 20 h post-induction. The clones marked red were selected for follow-up analysis and sequencing.

n.a.: ambigous sequencing results, typically duplicate sequence traces (non-clonal cultures).

#### Supplementary Table 5: Plasmids used in this work.

| Plasmid name[a] | Plasmid features[b] | Reference |
|---|---|---|
| pBAMD1-2 | $Ap^R$, $Km^R$, *oriV*(R6K), mini-Tn5 transposon | [1] |
| pBAMD-tphII | pBAMD1-2 derivative; $P_{EM7}\rightarrow$tphA1$_{II}$, tphA2$_{II}$, tphA3$_{II}$, tphB$_{II}$ | This work |
| pBAMD-$P_{IbpA}$-EhaA-LCC | pBAMD1-2 derivative; $P_{IbpA}$-EhaA-LCC knock-in cassette, $Km^R$ | This work |
| pBAMD-$P_{IbpA}$-bglx-HiC | pBAMD1-2 derivative; $P_{IbpA}$-bglx-HiC knock-in cassette, $Km^R$ | This work |
| pBAMD-$P_{IbpA}$-lasb-IsP$^{Dura}$ | pBAMD1-2 derivative; $P_{IbpA}$-lasb-IsP$^{Dura}$ knock-in cassette, $Km^R$ | This work |
| pPS39 | Sm/Sp$^R$, *oriV*(pRO1600/ColE1), rhaS-rhaR-$P_{rhaB}$-MCS | [6] |
| pSEVA328 | $Cm^R$, *oriV*(RK2), XylS-$P_m$ | [7] |
| pSEVA338 | $Cm^R$, *oriV*(pBBR1), XylS-$P_m$ | [7] |
| pHi-$P_{RhaB}$ | pPS39 derivative; Sm/Sp$^R$ switched to $Km^R$ | This work |
| pSEVA338-$P_{rhaB}$ | pSEVA338 derivative; XylS-$P_m$ switched to rhaS-rhaR-$P_{rhaB}$-MCS | This work |
| pSEVA328-$P_{rhaB}$ | pSEVA328 derivative; XylS-$P_m$ switched to rhaS-rhaR-$P_{rhaB}$-MCS | This work |
| pMe-$P_{rhaB}$ | pSEVA338-$P_{rhaB}$ derivative; $Cm^R$ switched to $Km^R$ | This work |
| pLo-$P_{rhaB}$ | pSEVA328-$P_{rhaB}$ derivative; $Cm^R$ switched to $Km^R$ | This work |
| pLo-$P_{rhaB}$-LCC | pLo-$P_{rhaB}$ derivative, LCC ORF under control of $P_{rhaB}$ | This work |
| pLo-$P_{rhaB}$-HiC | pLo-$P_{rhaB}$ derivative, HiC ORF under control of $P_{rhaB}$ | This work |
| pLo-$P_{rhaB}$-IsP | pLo-$P_{rhaB}$ derivative, IsP ORF under control of $P_{rhaB}$ | This work |
| pHi-$P_{rhaB}$-OprF | pHi-$P_{rhaB}$ derivative; OprF membrane display anchor | This work |
| pHi-$P_{rhaB}$-OprF-LCC | pHi-$P_{rhaB}$-OprF with LCC cargo | This work |
| pHi-$P_{rhaB}$-OprF-HiC | pHi-$P_{rhaB}$-OprF with HiC cargo | This work |
| pMe-$P_{rhaB}$-OprF | pMe-$P_{rhaB}$ derivative; OprF membrane display anchor | This work |
| pMe-$P_{rhaB}$-OprF-LCC | pMe-$P_{rhaB}$-OprF with LCC cargo | This work |
| pMe-$P_{rhaB}$-OprF-HiC | pMe-$P_{rhaB}$-OprF with HiC cargo | This work |
| pLo-$P_{rhaB}$-OprF | pLo-$P_{rhaB}$ derivative; OprF membrane display system | This work |
| pLo-$P_{rhaB}$-OprF-LCC | pLo-$P_{rhaB}$-OprF with LCC cargo | This work |
| pLo-$P_{rhaB}$-OprF-HiC | pLo-$P_{rhaB}$-OprF with HiC cargo | This work |
| pLo-$P_{rhaB}$-OprF-IsP | pLo-$P_{rhaB}$-OprF with IsP cargo | This work |
| pHi-$P_{rhaB}$-EhaA-LCC | pHi-$P_{rhaB}$ derivative with EhaA membrane display system and LCC cargo | This work |
| pHi-$P_{rhaB}$-EhaA-HiC | pHi-$P_{rhaB}$ derivative with EhaA membrane display system and HiC cargo | This work |

| | | |
|---|---|---|
| pMe-$P_{rhaB}$-EhaA-LCC | pMe-$P_{rhaB}$ derivative with EhaA membrane display system and LCC cargo | This work |
| pMe-$P_{rhaB}$-EhaA-HiC | pMe-$P_{rhaB}$ derivative with EhaA membrane display system and HiC cargo | This work |
| pLo-$P_{rhaB}$-EhaA-LCC | pLo-$P_{rhaB}$ derivative with EhaA membrane display system and LCC cargo | This work |
| pLo-$P_{rhaB}$-EhaA-HiC | pLo-$P_{rhaB}$ derivative with EhaA membrane display system and HiC cargo | This work |
| pLo-$P_{rhaB}$-EhaA-IsP | pLo-$P_{rhaB}$ derivative with EhaA membrane display system and IsP cargo | This work |
| pHi-$P_{rhaB}$-InaV | pHi-$P_{rhaB}$ derivative; InaV membrane display anchor | This work |
| pHi-$P_{rhaB}$-InaV-LCC | pHi-$P_{rhaB}$-InaV with LCC cargo | This work |
| pHi-$P_{rhaB}$-InaV-HiC | pHi-$P_{rhaB}$-InaV with HiC cargo | This work |
| pMe-$P_{rhaB}$-InaV | pMe-$P_{rhaB}$ derivative; InaV membrane display anchor | This work |
| pMe-$P_{rhaB}$-InaV-LCC | pMe-$P_{rhaB}$-InaV with LCC cargo | This work |
| pMe-$P_{rhaB}$-InaV-HiC | pMe-$P_{rhaB}$-InaV with HiC cargo | This work |
| pLo-$P_{rhaB}$-InaV | pLo-$P_{rhaB}$ derivative; InaV membrane display anchor | This work |
| pLo-$P_{rhaB}$-InaV-LCC | pLo-$P_{rhaB}$-InaV with LCC cargo | This work |
| pLo-$P_{rhaB}$-InaV-HiC | pLo-$P_{rhaB}$-InaV with HiC cargo | This work |
| pLo-$P_{rhaB}$-SP(OprF)-LCC | pLo-$P_{rhaB}$-EhaA-LCC derivative; EhaA coding region and GS linker removed | This work |
| pLo-$P_{rhaB}$-SPswitch-LCC | pLo-$P_{rhaB}$-SP(OprF)-LCC derivative; BbsI site introduced in BCD2 translational coupler | This work |
| pLo-$P_{rhaB}$-SPswitch-HiC | pLo-$P_{rhaB}$-SPswitch-LCC derivative; LCC cargo replaced with HiCut | This work |
| pLo-$P_{rhaB}$-SPswitch-IsP | pLo-$P_{rhaB}$-SPswitch-LCC derivative; LCC cargo replaced with IsP | This work |
| pLo-$P_{rhaB}$-ispe-LCC | pLo-$P_{rhaB}$-SPswitch-LCC derivative with ispe SP | This work |
| pLo-$P_{rhaB}$-flid-LCC | pLo-$P_{rhaB}$-SPswitch-LCC derivative with flid SP | This work |
| pLo-$P_{rhaB}$-efeb-LCC | pLo-$P_{rhaB}$-SPswitch-LCC derivative with efeb SP | This work |
| pLo-$P_{rhaB}$-bglx-HiC | pLo-$P_{rhaB}$-SPswitch-HiC derivative with bglx SP | This work |
| pLo-$P_{rhaB}$-mcoa-HiC | pLo-$P_{rhaB}$-SPswitch-HiC derivative with mcoa SP | This work |
| pLo-$P_{rhaB}$-lasb-IsP | pLo-$P_{rhaB}$-SPswitch-IsP derivative with lasb SP | This work |
| pLo-$P_{rhaB}$-agde-IsP | pLo-$P_{rhaB}$-SPswitch-IsP derivative with agde SP | This work |
| pLo-$P_{rhaB}$-copb-IsP | pLo-$P_{rhaB}$-SPswitch-IsP derivative with copb SP | This work |
| pLo-$P_L$ | pLo-$P_{rhaB}$ derivative; rhaS-rhaR-$P_{RhaB}$ switched to cI857/$P_L$ | This work |

| | | |
|---|---|---|
| pLo-$P_L$-EhaA-LCC | pLo-$P_L$ derivative with EhaA-LCC membrane display cargo | This work |
| pLo-$P_L$-bglx-HiC | pLo-$P_L$ derivative with bglx-HiC secretion construct cargo | This work |
| pLo-$P_{IbpA}$ | pLo-$P_{rhaB}$ derivative; rhaS-rhaR-$P_{rhaB}$ switched to $P_{IbpA}$ | This work |
| pLo-$P_{IbpA}$-EhaA-LCC | pLo-$P_{IbpA}$ derivative with EhaA-LCC membrane display cargo | This work |
| pLo-$P_{IbpA}$-bglx-HiC | pLo-$P_{IbpA}$ derivative with bglx-HiC secretion construct cargo | This work |
| pLo-$P_{IbpA}$-lasb-IsP$^{Dura}$ | pLo-$P_{IbpA}$ derivative with lasb-IsP$^{Dura}$ secretion construct cargo | This work |
| pMe-$P_{rhaB}$-mhpT | pMe-$P_{rhaB}$ derivative with mhpT$^{wt}$ ORF | This work |
| pMe-$P_{rhaB}$-mhpT$^{L374V}$ | pMe-$P_{rhaB}$ derivative with mhpT$^{L374V}$ ORF | This work |

[a] Naming of expression plasmids generated in this work follows cellular copy number: Hi, high copy number, *oriV*(pRO1600/ColE1); Me, medium copy number, *oriV*(pBBR1), Lo, low copy number, *oriV*(RK2).

[b] MCS multiple cloning site; Ap ampicillin; Cm chloramphenicol; Km kanamycin; Sm/Sp streptomycin/spectinomycin.

[c] The SEVA plasmid database: http://seva-plasmids.com/

All plasmid maps and sequences are available on request.

**Supplementary Table 6: Bacterial strains used in this work.**

| Strain name | Strain features[a] | Source / Reference |
|---|---|---|
| *E. coli* NEB 5-alpha | DH5α derivative; general cloning strain | NEB |
| *E. coli* CC118 *λpir* | Cloning and maintenance of *oriV*(R6K) plasmids | SEVA |
| *P. putida* KT2440 | *P. putida* reference strain, ATCC #47054 | ATCC |
| *P. putida* TA | *P. putida* KT2440 with *tphII* operon knock in | This work |
| *P. putida* TA7 | *P. putida* TA derivative, adapted for growth in M9-TA medium at pH ≥ 7; $mhpT^{L374V}$, $PP\_3359^{R51C}$ | This work |
| *P. putida* TA7-EG | *P. putida* TA7 derivative, adapted for growth in M9-EG medium; $gclR^{E130*}$, $PP\_4384^{A13fs}$ | This work |
| *P. putida* TA7-BD | *P. putida* TA7 derivative, adapted for growth in M9-BD medium; $PP\_2046^{V136A}$ | This work |
| *P. putida* KT2440 $P_{IbpA}$-EhaA-LCC | *P. putida* KT2440 with $P_{IbpA}$-EhaA-LCC knock-in | This work |
| *P. putida* KT2440 $P_{IbpA}$-bglx-HiC | *P. putida* KT2440 with $P_{IbpA}$-bglx-HiC knock-in | This work |
| *P. putida* KT2440 $P_{IbpA}$-lasb-$IsP^{Dura}$ | *P. putida* KT2440 with $P_{IbpA}$-lasb-$IsP^{Dura}$ knock-in | This work |
| *P. putida* TA7-EG $P_{IbpA}$-EhaA-LCC | *P. putida* TA7-EG with $P_{IbpA}$-EhaA-LCC knock-in | This work |
| *P. putida* TA7-EG $P_{IbpA}$-bglx-HiC | *P. putida* TA7-EG with $P_{IbpA}$-bglx-HiC knock-in | This work |
| *P. putida* TA7-BD $P_{IbpA}$-EhaA-LCC | *P. putida* TA7-BD with $P_{IbpA}$-EhaA-LCC knock-in | This work |
| *P. putida* TA7-BD $P_{IbpA}$-bglx-HiC | *P. putida* TA7-BD with $P_{IbpA}$-bglx-HiC knock-in | This work |
| *P. putida* TA7-BD $P_{IbpA}$-lasb-$IsP^{Dura}$ | *P. putida* TA7-BD with $P_{IbpA}$-lasb-$IsP^{Dura}$ knock-in | This work |

[a] See Supplementary Tables 1 and 2 for details of whole-genome sequencing and landing sites of *tphII* or $P_{IbpA}$-PET hydrolase knock-in cassettes.

**Supplementary Table 7: Oligonucleotides used in this work.**

| Name | Sequence (5' → 3')[a] |
|---|---|
| OB40 | GCCACTGGATGATGGACTCCTGCATTAGAAAACCTCCTTAGCATG |
| OB51 | GCCTAGGCCGCGGCCGCGCGAATTCTTGTTGACAATTAATCATCGGCATAGTATATCGGC |
| OB39 | CATGCTAAGGAGGTTTTCTAATGCAGGAGTCCATCATCCAGTGGC |
| OB54 | GTAGATCCCTCGTGTGTGCGCCTGAAGTCTTATCAGGCTTGCATCTCTTGACCCATG |
| OB55 | TAAGACTTCAGGCGCACACACGAGGGATCTACATGATCAATGAGATTCAAATTGCAGCG |
| OB56 | GATAGTCCCTCGTGTGTGGAGCCCAGTTACTATCACAGCGGCAAAGCCATGAGGG |
| OB57 | TAGTAACTGGGCTCCACACACGAGGGACTATCATGACCATTGTGCATCGGCGCC |
| OB58 | AGTGTACCCTCGTGTGTGGGTCTATCGGATCATCAGACGGGCTGTGCCCCCAG |
| OB59 | CCGATAGACCCACACACGAGGGTACACTATGAACCACCAAATCCATATCCATGATAGCG |
| OB53 | GGGTCCGCAATTAATTAGACAAGGGTCTCAGTGATGACCCGTTGGCCCGGTAGCAAAGAC |
| OB179 | CGACTGAGCCTTTCGTTTTATTTGATGCCTTTAATTAAGAACGGTCG |
| OB192 | CGGCCTAGGCATATGGAATCTCCTGAATTATCATCAAGTAC |
| OB191 | CAGGAGATTCCATATGCCTAGGCCGCGGCCGCGCG |
| OB189 | CTACACGAAATTCATCTGGGAGAAGTACCGC |
| OB178 | CGACCGTTCTTAATTAAAGGCATCAAATAAAACGAAAGGCTCAGTCG |
| OB190 | GCGGTACTTCTCCCAGATGAATTTCGTGTAG |
| OB182 | GATGCCTTTAATTAATCAGCCAAACGTCTCTTCAG |
| OB183 | GGGCCCCCTAGGTAATCCGGAATCGCACTTACG |
| OB85 | GACTTCCCTAGGGGGCCCAAGTTCACTTAAAAAGGAGATCAACAATGAAAGCAATTTTCG |
| OB91 | CATTAGAAAACCTCCTTAGCATGATTAAGATG |
| OB92 | CATCTTAATCATGCTAAGGAGGTTTTCTAATGAAACTGAAAAACACCTTGGGCTTGGC |
| OB93 | CCTGAAAAGCTTTCAGTGGTGGTGGTGATGATGC |
| OB86 | GCCACCCGAACCACCCGAGCCGCCCCCGCCCCCATGGCCCTGACAATGC |
| OB94 | GGCTCCCCTGCAGGCCAACTCGGTGCG |
| OB95 | ATGATGCCCATGGCCCGCCCGAATCCG |
| OB132 | TATATACCTGCAGGCCAGACCAACCC |
| OB133 | TATATACCCATGGCCCGAACAGTTCGC |
| OB87 | GCGGGGGCGGCTCGGGTGGTTCGGGTGGCGGTGGTTCCGGTGG |
| OB88 | CCTGAAAAGCTTTTAGAACTGCCACTTGATGCCCACCATCGCGGAGGTG |
| OB89 | CCTGAAAAGCTTTCAGTGGTGGTGGTGATGATGCCCATGGCCGGGTCTATCGGATGAGC |
| OB90 | CATCTTAATCATGCTAAGGAGGTTTTCTAATGAACATCGACAAGGCACTCGTCCTG |
| OB121 | CCAGTCACGACGCTGAGGTGCGATCGCGCGGCCGCAAGCTTTTACCC |
| OB122 | GCGACTTTCGCACGAATAACCGGCATTGTCAGGGCCATGGGTAAAAGCTTGCGGCCGCGC |

| | |
|---|---|
| OB134 | CATCTTAATCATGCTAAGGAGGTCTTCTAATGAAACTGAAAAACACCTTGG |
| OB135 | CCAAGGTGTTTTTCAGTTTCATTAGAAGACCTCCTTAGCATGATTAAGATG |
| OB136 | CATGCTAAGGAGGTCTTC |
| OB137 | GATTGGATTGGCCTGCAGG |
| OB139 | ACGCTGAGGTGCGATCGCGCGGCCGC |
| OB211 | ATCATGCTAAGGAGGTCTTCTAATGCAATCCAATCCATACCAGCGGGG |
| OB210 | ATCATGCTAAGGAGGTCTTCTAATGCAACTCGGTGCGATTGAAAATGGTC |
| OB209 | ATCATGCTAAGGAGGTCTTCTAATGCAGACCAACCCATACGCACGTG |
| OB201 | CAGAGTACTTGATGATAATTCAGGAGATTCCATATGAAACTGAAAAACACCTTGGGCTTG |
| OB197 | GATTCCATATGATGAAATTGAGCTTGCTCGGTCTC |
| OB195 | GATTCCATATGAAAAAGGTCTCGACGCTGGACCTG |
| OB220 | CGGCCGCAAGCTTTTACGAACAGTTCGCGGTGCG |
| OB221 | GATGCCTTTAATTAAGAACGGTCGTTCTCAAGGCTCTTG |
| OB143 | CCTTTCGGCGGGCTTTGGCGGCCGCAAGCTTTTACCCATGGCCCGCCCGAATCC |
| OB205 | CGAATTCGAGCTCGAACGGTCGTTCTCAAGGCTCTTG |
| OB119 | CCTTTCGGCGGGCTTTGGCGGCCGCAAGCTTTTAGAACTGCCACTTGATGCCCACC |
| OB214 | CGCCTAGGCCGCGGCCGCGCGAATTCGAGCTCGAACGGTCGTTCTCAAGGCTCTTG |
| OB109 | CGATCGCCCTAGGACGAGGGATCTACATGCATCATTCATGCGCGTCATCGG |
| OB110 | CGGCCGCAAGCTTCTATGCAGGGTCCTGGGCTGCAG |
| OB96 | GGCACGCGTCGACTAGTACNNNNNNNNNNACGCC |
| OB105 | GTAAATACGCTGCTGCTCTTGGTC |
| OB97 | GGCACGCGTCGACTAGTAC |
| OB104 | CACTGGATGATGGACTCCTGC |
| OB102 | ACCCATTTCGGGAATTTGCATGAC |
| OB103 | CAGGACTTCCATGCCGACGTC |

[a] Oligos are not ordered by name, but chronologically as discussed in the *Materials and Methods* section.

#### III Synthetic DNA sequences

**Supplementary sequence 1:** $P_{EM7}$-BCD2 promoter-bicistronic translational coupler fragment.

CTGTCTCTTATACACATCTTTGTGTCTCAGGCCGTTGTTGACAATTAATCATCGGCATAGTA
TATCGGCATAGTATAATACGACAAGGTGAGGAACTAAACCCCTAGGGCCCAAGTTCACTTA
AAAAGGAGATCAACAATGAAAGCAATTTTCGTACTGAAACATCTTAATCATGCTAAGGAGGT
TTTCTAATG

**Supplementary sequence 2:** *Comamonas sp*. E6, *tphA2*$_{II}$, Genbank AB238679.1 and BAE47085.1, oxygenase large subunit of terephthalate 1,2-dioxygenase, codon-optimized for *P. putida.*

ATGCAGGAGTCCATCATCCAGTGGCATGGGGCGACCAATACCCGGGTGCCGTTCGGCATT
TACACCGACACGGCGAATGCTGACCAAGAGCAGCAGCGTATTTACCGTGGTGAAGTCTGG
AACTACCTCTGCCTGGAAAGCGAAATCCCTGGTGCGGGGGACTTCCGTACGACCTTCGCG
GGCGAAACGCCCATTGTGGTGGTCCGCGACGCTGACCAGGAAATTTATGCATTCGAAAAT
CGGTGCGCTCATCGTGGGGCGCTGATCGCTCTCGAAAAAAGCGGCCGTACGGACTCGTTT
CAATGCGTCTACCATGCCTGGAGCTATAATCGTCAGGGGGATTTGACCGGGGTCGCTTTT
GAAAAGGGGGTGAAGGGGCAAGGTGGTATGCCGGCGTCCTTTTGTAAAGAGGAACATGGT
CCTCGTAAGTTGCGCGTGGCCGTCTTTTGTGGTTTGGTCTTTGGCTCCTTCTCGGAGGATG
TGCCATCGATTGAGGACTATCTGGGGCCAGAAATCTGCGAGCGGATCGAACGTGTCCTCC
ATAAGCCAGTCGAAGTCATTGGCCGTTTCACGCAAAAGCTCCCGAACAACTGGAAGCTGTA
TTTCGAAAACGTGAAGGATAGCTATCATGCCTCCCTCTTGCACATGTTTTTTACGACGTTCG
AGTTGAATCGGCTCTCCCAGAAGGGGGGCGTGATCGTCGATGAATCGGGTGGTCACCAC
GTCTCGTATAGCATGATCGACCGGGGGGCGAAGGATGACTCGTACAAGGATCAGGCCATC
CGTTCCGACAATGAACGTTATCGTCTGAAAGACCCGTCGCTCCTCGAGGGCTTTGAGGAAT
TTGAGGACGGTGTCACCCTCCAGATCTTGTCGGTGTTCCCGGGTTTTGTGCTCCAGCAGAT
CCAAAATAGCATCGCCGTGCGTCAATTGCTGCCTAAGAGCATCAGCTCGTCGGAATTGAAT
TGGACCTATCTGGGGTATGCGGACGATTCGGCGGAACAACGGAAGGTGCGCTTGAAACAG
GCCAATCTCATCGGTCCCGCCGGTTTCATCTCCATGGAAGATGGGGCGGTGGGGGGCTTC
GTCCAACGGGGGATTGCCGGTGCAGCGAATTTGGACGCCGTGATCGAGATGGGTGGGGA

CCATGAAGGGTCCTCCGAGGGCCGCGCCACGGAGACCAGCGTGCGTGGGTTTTGGAAGG
CGTACCGTAAGCACATGGGTCAAGAGATGCAAGCCTGA

**Supplementary sequence 3:** *Comamonas sp*. E6, *tphA3*$_{II}$, Genbank AB238679.1 and BAE47086.1, oxygenase small subunit of terephthalate 1,2-dioxygenase, codon-optimized for *P. putida.*

ATGATCAATGAGATTCAAATTGCAGCGTTTAACGCTGCGTATGCCAAAACCATCGACTCCG
ATGCTATGGAACAGTGGCCGACCTTCTTCACGAAAGATTGCCATTATTGCGTGACCAACGT
CGATAACCACGATGAGGGTCTGGCCGCTGGCATTGTCTGGGCTGACTCCCAAGATATGTT
GACCGACCGCATTTCCGCTTTGCGCGAGGCCAATATCTACGAACGGCACCGCTACCGTCA
TATCCTGGGTCTGCCGTCCATTCAGTCGGGGGACGCAACGCAAGCGTCCGCCTCCACGC
CGTTCATGGTCCTCCGCATTATGCACACGGGCGAAACGGAGGTGTTCGCGTCGGGCGAAT
ACCTCGACAAATTCACGACCATCGATGGTAAACTGCGCCTCCAGGAGCGGATCGCAGTCT
GCGACTCCACCGTGACCGATACCCTCATGGCTTTGCCGCTGTGA

**Supplementary sequence 4:** *Comamonas sp*. E6, *tphB*$_{II}$, Genbank AB238679.1 and BAE47087.1, 1,2-dihydroxy-3,5-cyclohexadiene-1,4-dicarboxylate dehydrogenase, codon-optimized for *P. putida.*

ATGACCATTGTGCATCGGCGCCTCGCACTCGCAATTGGTGATCCGCACGGTATTGGTCCA
GAGATTGCGCTCAAAGCGCTGCAACAACTGAGCGTCACGGAACGCTCGCTGATTAAAGTG
TATGGCCCATGGTCCGCGTTGGAACAAGCCGCTCGTGTCTGCGAAATGGAACCTTTGTTG
CAAGACATTGTCCATGAAGAGGCAGGGACCCTGACGCAGCCCGTGCAGTGGGGCGAAAT
CACGCCACAGGCTGGCCTCTCCACGGTGCAATCCGCGACGGCCGCGATCCGCGCTTGTG
AGAACGGCGAAGTGGATGCCGTGATTGCCTGCCCGCATCACGAGACGGCCATCCACCGT
GCAGGCATCGCGTTCTCCGGGTACCCGTCCCTGTTGGCGAACGTCCTCGGTATGAATGAG
GACCAAGTGTTTCTGATGCTGGTGGGCGCCGGTCTCCGCATTGTCCACGTCACGCTCCAC
GAGAGCGTGCGTAGCGCATTGGAGCGTTTGTCGCCGCAGCTGGTGGTCAATGCTGCGCA
AGCAGCGGTCCAAACGTGCACCCTCTTGGGTGTCCCTAAGCCCAAAGTCGCAGTCTTCGG
TATTAATCCTCATGCAAGCGAGGGCCAGCTGTTCGGCCTGGAGGATTCGCAGATCACCGT
CCCGGCAGTCGAAACGTTGCGCAAACGCGGCTTGGCGGTGGATGGTCCAATGGGCGCCG
ACATGGTGCTCGCGCAGCGGAAACATGATCTGTATGTGGCAATGCTGCACGACCAAGGCC

ACATCCCCATTAAATTGCTCGCACCCAATGGCGCTAGCGCCCTCTCCATTGGGGGTCGCG
TCGTCCTGTCGTCCGTCGGCCATGGGTCGGCGATGGATATCGCTGGTCGTGGTGTCGCTG
ATGCCACCGCGCTGCTGCGTACCATTGCATTGCTGGGGGCACAGCCCGTCTGA

**Supplementary sequence 5:** *Comamonas sp*. E6, *tphA1*$_{II}$, Genbank AB238679.1 and BAE47088.1, reductase component of terephthalate 1,2-dioxygenase, codon-optimized for *P. putida.*

ATGAACCACCAAATCCATATCCATGATAGCGACATCGCGTTCCCGTGCGCTCCTGGTCAGA
GCGTCCTCGATGCAGCACTCCAGGCAGGGATTGAATTGCCGTACTCCTGTCGTAAAGGCT
CGTGCGGGAACTGCGCCTCGACCTTGTTGGACGGTAATATTGCATCCTTTAATGGCATGG
CAGTCCGTAATGAGTTGTGTGCCTCCGAGCAAGTGCTCCTGTGTGGGTGTACCGCCGCGA
GCGATATCCGGATTCACCCATCCTCGTTCCGCCGGCTGGACCCAGAAGCTCGCAAACGGT
TCACCGCGAAAGTCTATTCGAATACGCTCGCCGCGCCCGATGTCTCCTTGTTGCGGCTGC
GGCTGCCAGTGGGTAAACGCGCAAAATTTGAAGCAGGTCAGTACCTCTTGATCCACCTCG
ATGATGGCGAATCGCGGTCGTACTCCATGGCGAATCCCCCTCATGAGTCGGATGGTATCA
CGCTGCACGTGCGGCACGTCCCGGGTGGGCGTTTCAGCACCATCGTGCAACAACTGAAAA
GCGGGGATACGTTGGACATCGAATTGCCGTTCGGTTCCATCGCCTTGAAGCCGGATGATG
CTCGTCCTCTCATCTGTGTGGCCGGTGGTACCGGGTTTGCACCTATTAAGTCGGTCCTCGA
CGACTTGGCGAAACGGAAAGTGCAACGGGATATTACGCTGATTTGGGGCGCTCGTAATCC
TAGCGGCCTCTACTTGCCAAGCGCGATCGACAAATGGCGCAAAGTCTGGCCACAGTTCCG
TTACATCGCAGCTATCACGGATCTCGGGGATATGCCGGCTGATGCTCACGCTGGCCGCGT
GGACGATGCATTGCGCACCCATTTCGGGAATTTGCATGACCACGTGGTGCACTGCTGTGG
TTCCCCCGCGTTGGTCCAATCGGTGCGGACGGCCGCGAGCGACATGGGGTTGCTGGCTC
AGGACTTCCATGCCGACGTCTTTGCTACCGGGCCAACGGGTCATCACTGA

**Supplementary sequence 6:** $P_{IbpA}$ oligo (ssDNA) fragment, including parts of the T1 terminator and PacI site at the 5’ end for cloning purposes (the actual $P_{IbpA}$ fragment is underlined).

gatgcctttaatTAAGAACGGTCGTTCTCAAGGCTCTTGAAATACTTCCGACGATCCCTATCTAGT
TCGTGCGGGATGCCGAAGACGGGTTCCGCATCAGAGTACTTGaTGATAATTCAGGAGATTC

**Supplementary sequence 7:** cI857/$P_L$ fragment, adapted for the SEVA plasmid system (based on Aparicio *et al*., 2019).[8]

TCAGCCAAACGTCTCTTCAGGCCACTGACTAGCGATAACTTTCCCCACAACGGAACAACTC
TCATTGCATGGGATCATTGGGTACTGTGGGTTTAGTGGTTGTAAAAACACCTGACCGCTAT
CCCTGATCAGTTTCTTGAAGGTAAACTCATCACCCCCAAGTCTGGCTATGCAGAAATCACC
TGGCTCAACAGCCTGCTCAGGGTCAACGAGAATTAACATTCCGTCAGGAAAGCTCGGCTT
GGAGCCTGTTGGTGCGGTCATGGAATTACCTTCAACCTCAAGCCAGAATGCAGAATCACTG
GCTTTTTTGGTTGTGCTTACCCATCTCTCCGCATCACCTTTGGTAAAGGTTCTGAGCTTAGG
TGAGAACATCCCTGCCTGAACATGAGAAAAAACAGGGTACTCATACTCACTTCTAAGTGAC
GGCTGCATACTAACCGCTTCATACATCTCGTAGATTTCTCTGGCGATTGAAGGGCTAAATT
CTTCAACGCTAACTTTGAGAATTTTTGTAAGCAATGCGGCGTTATAAGCATTTAATGCATTG
ATGCCATTAAATAAAGCACCAACGCCTGACTGCCCCATCCCCATCTTGTCTGCGACAGATT
CCTGGGATAAGCCAAGTTCATTTTTCTTTTTTTCATAAATTGCTTTAAGGCGGCGTGCGTCC
TCAAGCTGCTCTTGTGTTAATGGTTTCTTTTTTGTGCTCATGCTAGCTTTTTCCTCCTTATAA
AGTTAATCAACACCCCTTGTATTACTGTTTATGTAAGCAGACAGTTTTATTGTTCATGATGAT
ATATTTTTATCTTGTGCAATGTAACATCAGAGATTTTGAGACACAAGACGTGGCAAAAAACA
TTATCCAGAACGGGAGTGCGCCTTGAGCGACACGAATTATGCAGTGATTTACGACCTGCAC
AGCCATACCACAGCTTCCGATGGCTGCCTGACGCCAGAAGCATTGGTGCACCGTGCAGTC
GATGATAAGCTGTCAAACAGCGGATAACAATTTCACACAGGATAACCCGGCCTCAGCGCC
GGGTTTTCTTTGCCTCACGATCGCCCCCAAAACACATAACCAATTGTATTTATTGAAAAATA
AATAGATACAACTCACTAAACATAGCAATTCAGATCTCTCACCTACCAAACAATGCCCCCCT
GCAAAAAATAAATTCATATAAAAAACATACAGATAACCATCTGCGGTGATAAATTATCTCTG
GCGGTGTTGACATAAATACCACTGGCGGTGATACTGAGCACATCAGCAGGACGCACTGAC
CACCATGAAGGTGACGCTCTTAAAAATTAAGCCCTGAAGAAGGGCAGCATTCAAAGCAGAA
GGCTTTGGGGTGTGTGATACGAAACGAAGCATTGGCCGTAAGTGCGATTCCGGATTA

**Supplementary sequence 8:** BCD2-Leaf compost cutinase (LCC) fragment (LCC: residues 35-293, Uniprot G9BY57, GenBank AEV21261.1), including the N-terminal BCD2 bicistronic linker, followed by residues 1-37 of *P. putida* OprF (as described by Tozakidis *et al*., 2019)[9], a 6xHis tag, and a PstI restriction site, codon-optimized for *P. putida.*

cacttaaaaaggagatcaacaatgaaagcaattttcgtactgaaacatcttaatcatgctaaggaggttttctaATGAAACTGAA
AAACACCTTGGGCTTGGCCATTGGTTCGCTTGTAGCCGCCACTTCGATTGGCGCTATGGC

ACAAGGTCAAGGCGCCGTCGAGACTGAAATCTTCTACAAGCATCATCACCACCACCACcctg
caggcCAATCCAATCCATACCAGCGGGGTCCAAATCCTACGCGTTCCGCGCTGACGGCAGA
CGGGCCGTTCTCCGTCGCTACCTATACCGTCTCCCGTCTCTCCGTGTCCGGTTTCGGCGG
CGGTGTCATCTACTATCCCACGGGTACCAGCCTGACCTTCGGGGGGATTGCCATGTCGCC
AGGCTATACGGCAGATGCCAGCTCCTTGGCCTGGCTCGGCCGTCGTCTCGCTTCCCACGG
TTTCGTCGTCCTCGTCATCAACACGAACTCGCGTTTTGATTATCCTGATTCGCGGGCGTCG
CAGCTGAGCGCTGCGCTGAACTATTTGCGGACGAGCTCCCCGTCCGCAGTGCGTGCACG
TCTCGACGCAAATCGTTTGGCCGTCGCAGGGCACAGCATGGGTGGCGGGGGTACGTTGC
GGATCGCAGAGCAGAATCCCTCGCTCAAAGCGGCAGTCCCGCTCACGCCGTGGCACACC
GACAAGACCTTCAATACGTCCGTCCCAGTCCTCATCGTCGGCGCTGAGGCCGATACGGTG
GCCCCTGTGTCCCAGCACGCAATTCCATTTTACCAGAACCTGCCCTCCACCACGCCTAAAG
TCTATGTCGAACTGGACAACGCTTCGCATTTTGCCCCTAATAGCAACAATGCAGCAATTTCC
GTGTACACGATTTCGTGGATGAAACTGTGGGTCGACAACGATACGCGGTATCGGCAGTTC
CTCTGCAATGTCAATGACCCCGCTCTGAGCGACTTTCGCACGAATAACCGGCATTGTCAGg
gccatgggGG

**Supplementary sequence 9:** *Pseudomonas putida* KT2440 OprF (Uniprot Q88L46, GenBank AAN67703.1), residues 1-206, including a C-terminal $G_4SG_2S(G_4S)_3$ linker, PstI and NcoI restriction sites, 6xHis tag, and HindIII restriction site.

ATGAAACTGAAAAACACCTTGGGCTTGGCCATTGGTTCGCTTGTAGCCGCCACTTCGATTG
GCGCTATGGCACAAGGTCAAGGCGCCGTCGAGACTGAAATCTTCTACAAGAAAGAgTTCTT
CGACAGCCAGCGCGACTTCAAGAACGACGGCAACCTGTTCGGCGGCTCGATCGGTTACTT
CCTGACCGACGACGTTGAGCTGCGTCTGGGCTACGACGAAGTGCACAACGCTCGTGGCG
AAGACGGCAAGAACATCAAGGGCTCGAACACTGCCCTGGACGCCGTTTACCACTTCAACA
ACCCGTACGACGCTATCCGTCCATACGTTTCCGCTGGTTTCTCGCACCAGTCGCTGGGtCA
GACCGGCCGTGGCGGTCGTGACCACTCCACCTTCGCCAACGTTGGCGCTGGCGCCAAGT
GGTACATCACCGACATGTTCTATGCCCGTGCCGGCGTAGAAGCTCAGTACAACATCGACC
AGGGCGACACCGAGTGGGCCCCGAGCGTTGGTGTTGGCCTGAACTTCGGCGGTAGCCCG
AAGCAAGCTGAAGCTGCTCCTGCTCCAGTTGCTGAAGTCTGCTCCGACTCCGACAACGAC
GGCGTtTGCGAtAAtGTCGGcGGtGGtGGCagcGGTGGcTCcGGcGGcGGTGGTagtGGTGGcG
GtGGTTCCGGCGGtGGCGGCagccctgcaggcTCATCCGATAGACCCggccatgggCATCATCACC
ACCACCACtgaaagctt

**Supplementary sequence 10:** *Humicola insolens* cutinase (HiC) (Uniprot A0A075B5G4, GenBank ASK40094.1), including a N-terminal PstI and a C-terminal NcoI restriction site, codon-optimized for *P. putida.*

cctgcaggcCAACTCGGTGCGATTGAAAATGGTCTCGAAAGCGGCTCGGCGAATGCGTGCCC
AGACGCCATTCTGATTTTCGCTCGTGGTTCGACCGAACCCGGTAACATGGGGATCACGGT
GGGCCCAGCTTTGGCCAACGGTTTGGAATCGCACATCCGTAACATTTGGATCCAAGGTGT
GGGTGGCCCCTACGACGCGGCACTCGCTACGAATTTTCTCCCACGTGGTACCAGCCAGGC
GAATATTGACGAAGGTAAACGTCTGTTTGCATTGGCGAATCAGAAGTGTCCCAATACCCCT
GTCGTCGCGGGGGGCTATTCCCAAGGTGCGGCGCTCATCGCTGCGGCTGTGTCCGAATT
GTCCGGCGCCGTGAAGGAGCAAGTGAAAGGTGTGGCTCTCTTCGGGTATACCCAGAATCT
CCAAAATCGGGGTGGCATCCCTAATTACCCGCGTGAACGGACGAAGGTGTTTTGCAACGT
CGGGGATGCCGTCTGCACCGGCACCCTGATTATTACCCCTGCTCATCTGTCGTATACCATT
GAGGCTCGCGGTGAGGCAGCCCGGTTCTTGCGCGATCGGATTCGGGCGggccatggg

**Supplementary sequence 11:** *Ideonella sakaiensis* PETase, residues 28-290 (Uniprot A0A0K8P6T7, GenBank BBYR01000074.1), codon-optimized for *P. putida.*

CAGACCAACCCATACGCACGTGGTCCCAATCCTACCGCTGCCAGCCTCGAAGCCAGCGCG
GGCCCATTTACGGTGCGTTCGTTTACGGTGTCCCGGCCTAGCGGTTATGGGGCGGGCAC
CGTCTACTATCCAACGAATGCTGGCGGGACGGTCGGCGCcATCGCAATTGTCCCTGGTTA
CACGGCTCGCCAGTCCAGCATCAAATGGTGGGGGCCGCGGCTCGCATCCCACGGTTTTG
TGGTGATTACGATTGACACGAACTCGACCCTCGACCAGCCTAGCTCGCGGTCGTCGCAAC
AAATGGCCGCGCTGCGCCAAGTGGCATCCCTCAATGGCACCAGCTCcAGCCCGATCTATG
GCAAAGTCGATACGGCACGTATGGGGGTGATGGGTTGGTCGATGGGCGGTGGCGGTTCG
TTGATCTCCGCAGCGAACAATCCGTCCTTGAAGGCAGCTGCACCACAAGCGCCCTGGGAT
TCGTCGACGAACTTTTCGAGCGTCACGGTGCCGACCTTGATTTTCGCCTGTGAGAACGATT
CCATTGCACCCGTGAACTCGTCCGCCCTGCCAATCTACGATTCCATGTCGCGTAATGCAAA
GCAGTTCTTGGAGATTAATGGGGGTTCGCACAGCTGCGCCAACTCCGGGAACAGCAACCA
GGCCTTGATTGGTAAGAAAGGGGTCGCTTGGATGAAGCGTTTCATGGACAATGACACCCG
TTATAGCACGTTCGCGTGTGAGAATCCGAACTCGACCCGcGTgAGCGATTTTCGCACCGCG
AACTGTTCG

**Supplementary sequence 12:** *Escherichia coli* EhaA, residues 840-1323 (Uniprot A0A1Y2Y633, GenBank WP_023307657.1), including parts of an N-terminal $G_4SG_2S(G_4S)_3$ linker. Note: the last four 3'-terminal bases of EhaA were omitted to keep the fragment at 1500 bases to reduce synthesis cost and were re-introduced with primer OB88 during PCR amplification of this fragment; codon-optimized for *P. putida*.

CGGGTGGCGGTGGTTCCGGTGGcGGGGGTTCCGGCGGGGGCGGCTCCCTCACCAATAAT
GGTACgCTGATGACCGGCATGTCCGGTCAGCAAGCCGGTAACGTCTTGGTCGTGAAGGGC
AACTACCACGGTAACAATGGGCAACTCGTGATGAATACCGTGTTGAATGGTGATGATAGCG
TCACCGATAAACTCGTCGTGGAAGGCGACACCTCCGGCACCACGGCGGTCACGGTCAATA
ACGCTGGCGGTACgGGCGCTAAGACGCTGAACGGGATTGAACTGATTCATGTGGATGGGA
AGTCCGAAGGTGAATTTGTCCAGGCTGGTCGGATTGTGGCAGGGGCTTACGACTATACGC
TCGCTCGCGGCCAGGGCGCGAACTCGGGGAACTGGTATCTGACCTCGGGTTCGGATTCG
CCAGAGCTGCAACCCGAGCCAGATCCGATGCCGAATCCAGAGCCTAACCCAAATCCGGAA
CCAAACCCTAATCCCACCCCGACGCCGGGTCCAGATTTGAACGTCGATAATGATCTCCGC
CCAGAGGCTGGCTCCTACATTGCGAACCTCGCTGCTGCGAATACGATGTTCACCACCCGC
CTCCACGAACGGCTCGGCAACACGTACTACACGGACATGGTCACGGGCGAACAAAAGCAA
ACCACGATGTGGATGCGGCACGAAGGTGGGCACAACAAATGGCGTGACGGGAGCGGTCA
GCTGAAGACCCAAAGCAATCGCTATGTGTTGCAACTCGGGGGCGATGTGGCGCAGTGGA
GCCAAAATGGGTCGGACCGTTGGCACGTCGGGGTCATGGCCGGGTATGGGAATAGCGAC
TCCAAAACGATCAGCAGCCGCACCGGCTATCGCGCTAAGGCCTCGGTCAACGGTTACTCC
ACGGGCTTGTATGCGACGTGGTATGCCGACGATGAGTCGCGCAACGGCGCTTACCTCGAC
AGCTGGGCGCAATACTCCTGGTTCGACAACACGGTCAAGGGTGATGATCTGCAAAGCGAG
TCCTACAAGTCCAAAGGGTTCACCGCAAGtTTGGAAGCAGGGTATAAGCATAAATTGGCGG
AgTTCAACGGcTCCCAGGGcACCCGGAATGAGTGGTACGTCCAGCCTCAAGCGCAGGTGA
CGTGGATGGGCGTCAAAGCCGACAAACACCGGGAGAGCAATGGTACGCTCGTCCACAGC
AACGGCGACGGCAATGTCCAAACGCGTCTCGGCGTGAAGACCTGGTTGAAGAGCCATCAC
AAGATGGACGACGGGAAATCGCGTGAGTTTCAACCCTTCGTGGAGGTGAATTGGCTGCAC
AACAGCAAAGACTTTTCGACGTCgATGGACGGTGTCTCCGTGACGCAGGACGGGGCCCGC
AATATCGCTGAAATTAAAACGGGTGTGGAAGGTCAGCTCAATGCTAACCTCAACGTGTGGG
GCAATGTCGGGGTGCAGGTCGCGGACCGGGGCTACAACGACACCTCCGCgATGGTGGGC
ATCA

**Supplementary sequence 13:** *Pseudomonas syringae* InaV (Uniprot O33479, GenBank AJ001086), encompassing the N-terminal 609 bases followed by the C-terminal 291 bases; the fragment also encodes a C-terminal $G_4SG_2S(G_4S)_3$ linker and PstI / NcoI restriction sites, codon-optimized for *P. putida.*

ATGAACATCGACAAGGCACTCGTCCTGCGTACCTGCGCGAACAACATGGCGGACCACTGC
GGCCTGATTTGGCCCGCCAGCGGTACGGTGGAGTCCAAGTACTGGCAATCCACGCGGCG
CCATGAAAACGGGCTCGTGGGCTTGCTGTGGGGTGCAGGGACCTCGGCGTTTTTGAGCG
TGCACGCGGATGCCCGCTGGAAAGTGTGTGAAGTCGCGGTGGCCGACATCATTGGGCTG
GAAGAACCAGGCATGGTCAAATTTCCACGGGCGGAAGTCGTCCATGTCGGTGACCGCATT
TCGGCGAGCCATTTCATTAGCGCGCGGCAGGCCGATCCTGCCAGCACGCCAACCCCTAC
CCCAACGCCGATGGCGACCCCCACCCCAGCCGCAGCCAATATTGCTTTGCCAGTCGTCGA
GCAGCCGTCCCACGAAGTCTTTGACGTCGCCCTGGTGAGCGCAGCTGCTCCGTCCGTCAA
CACCCTCCCTGTCACGACGCCTCAAAACTTGCAAACCGCCACGTACGGTTCGACGTTGTC
GGGCGACAATAATTCCCGCCTGATTGCCGGCTACGGGTCCAATGAGACCGCGGGCAATCA
TTCCGATCTGATCGGCGGCCACGACTGTACCCTGATGGCGGGCGACCAGAGCCGCCTCA
CGGCTGGCAAAAACAGCATCCTGACGGCAGGGGCGCGTAGCAAACTCATCGGCTCGGAG
GGTTCCACGTTGTCCGCTGGGGAGGATTCGACGTTGATCTTCCGCCTCTGGGACGGGAAA
CGCTACCGTCAACTCGTGGCTCGTACGGGCGAGAACGGTGTGGAGGCAGACATTCCGTA
CTACGTCAACGAGGATGACGACATCGTCGATAAGCCGGACGAAGATGACGACTGGATCGA
AGTGGAGGGtGGtGGCGGCTCcGGTGGTTCGGGcGGcGGTGGTTCCGGTGGcGGtGGTTCC
GGCGGtGGCGGCTCCcctgcaggcTCATCCGATAGACCCggccatg

**Supplementary sequence 14:** $P_{IbpA}$-$IsP^{Dura}$ fragment, including the N-terminal $P_{IbpA}$ promoter-RNAT unit, followed by the ORF of DuraPETase as described by Cui et al., 2021;[10] codon-optimized for *P. putida*.

GAACGGTCGTTCTCAAGGCTCTTGAAATACTTCCGACGATCCCTATCTAGTTCGTGCGGGA
TGCCGAAGACGGGTTCCGCATCAGAGTACTTGaTGATAATTCAGGAGATTCcatATGAAAAA
GGTCTCGACGCTGGACCTGTTGTTTGTGGCTATCATGGGTGTCTCCCCGGCGGCCTTCGC
GGCTcctgcaggcCAGACCAACCCATACGCACGTGGTCCCAATCCTACCGCTGCCAGCCTCG
AAGCCAGCGCGGGCCCATTTACGGTGCGTTCGTTTACGGTGTCCCGGCCTAGCGGTTATG
GGGCGGGCACCGTCTACTATCCAACGAATGCTGGCGGGACGGTCGGCGCcATCGCAATT
GTCCCTGGTTACACGGCTCGCCAGTCCAGCATCAAATGGTGGGGGCCGCGGCTCGCATC

CCACGGTTTTGTGGTGATTACGATTGACACGAACTCGACCttcGACtacCCTAGCTCGCGGTC
GTCGCAACAAATGGCCGCGCTGCGCCAAGTGGCATCCCTCAATGGCgacAGCTCcAGCCC
GATCTATGGCAAAGTCGATACGGCACGTATGGGGGTGATGGGTcacTCGATGGGCGGTGG
CgccTCGTTGcgcTCCGCAGCGAACAATCCGTCCTTGAAGGCAGCTatcCCACAAGCGCCCTG
GGATTCGcagACGAACTTTTCGAGCGTCACGGTGCCGACCTTGATTTTCGCCTGTGAGAAC
GATTCCATTGCACCCGTGAACTCGcacGCCCTGCCAATCTACGATTCCATGTCGCGTAATGC
AAAGCAGTTCTTGGAGATTAATGGGGGTTCGCACAGCTGCGCCAACTCCGGGAACAGCAA
CCAGGCCTTGATTGGTAAGAAAGGGGTCGCTTGGATGAAGCGTTTCATGGACAATGACAC
CCGTTATAGCACGTTCGCGTGTGAGAATCCGAACTCGACCgccGTgAGCGATTTTCGCACC
GCGAACTGTTCGTAAa
